# PINDER: The protein interaction dataset and evaluation resource

**DOI:** 10.1101/2024.07.17.603980

**Authors:** Daniel Kovtun, Mehmet Akdel, Alexander Goncearenco, Guoqing Zhou, Graham Holt, David Baugher, Dejun Lin, Yusuf Adeshina, Thomas Castiglione, Xiaoyun Wang, Céline Marquet, Matt McPartlon, Tomas Geffner, Emanuele Rossi, Gabriele Corso, Hannes Stärk, Zachary Carpenter, Emine Kucukbenli, Michael Bronstein, Luca Naef

**Affiliations:** VantAI; NVIDIA Corporation; Massachusetts Institute of Technology; University of Oxford

## Abstract

Protein-protein interactions (PPIs) are fundamental to understanding biological processes and play a key role in therapeutic advancements. As deep-learning docking methods for PPIs gain traction, benchmarking protocols and datasets tailored for effective training and evaluation of their generalization capabilities and performance across real-world scenarios become imperative. Aiming to overcome limitations of existing approaches, we introduce PINDER, a comprehensive annotated dataset that uses structural clustering to derive non-redundant interface-based data splits and includes *holo* (bound), *apo* (unbound), and computationally predicted structures. PINDER consists of 2,319,564 dimeric PPI systems (and up to 25 million augmented PPIs) and 1,955 high-quality test PPIs with interface data leakage removed. Additionally, PINDER provides a test subset with 180 dimers for comparison to AlphaFold-Multimer without any interface leakage with respect to its training set. Unsurprisingly, the PINDER benchmark reveals that the performance of existing docking models is highly overestimated when evaluated on leaky test sets. Most importantly, by retraining DiffDock-PP on PINDER interface-clustered splits, we show that interface cluster-based sampling of the training split, along with the diverse and less leaky validation split, leads to strong generalization improvements.

## 1. Introduction

Proteins orchestrate numerous cellular processes, many of which are intricately tied to the way proteins interact with each other. The 3D structures of individual proteins and protein complexes inherently dictate their cellular functions, underscoring their critical role in drug design and therapeutic strategies (Lu et al., 2013). Deep learning (DL)-based methods for PPI complex modeling and docking have gained significant traction in recent years. However, the effective evaluation of these methods requires robust benchmarking protocols and datasets that can accurately assess their generalization capabilities and performance across diverse realworld scenarios. Existing benchmarking approaches often suffer from limitations such as data leakage between training and test sets, limited diversity in protein structures and interaction modes, and the lack of *apo* or predicted structures to evaluate docking performance under realistic conditions. To address these limitations, we introduce PINDER, the **P**rotein **IN**teraction **D**ataset and **E**valuation **R**esource, designed to facilitate the development and evaluation of next-generation computational protein docking methods (Figure 1).

**Figure 1.**
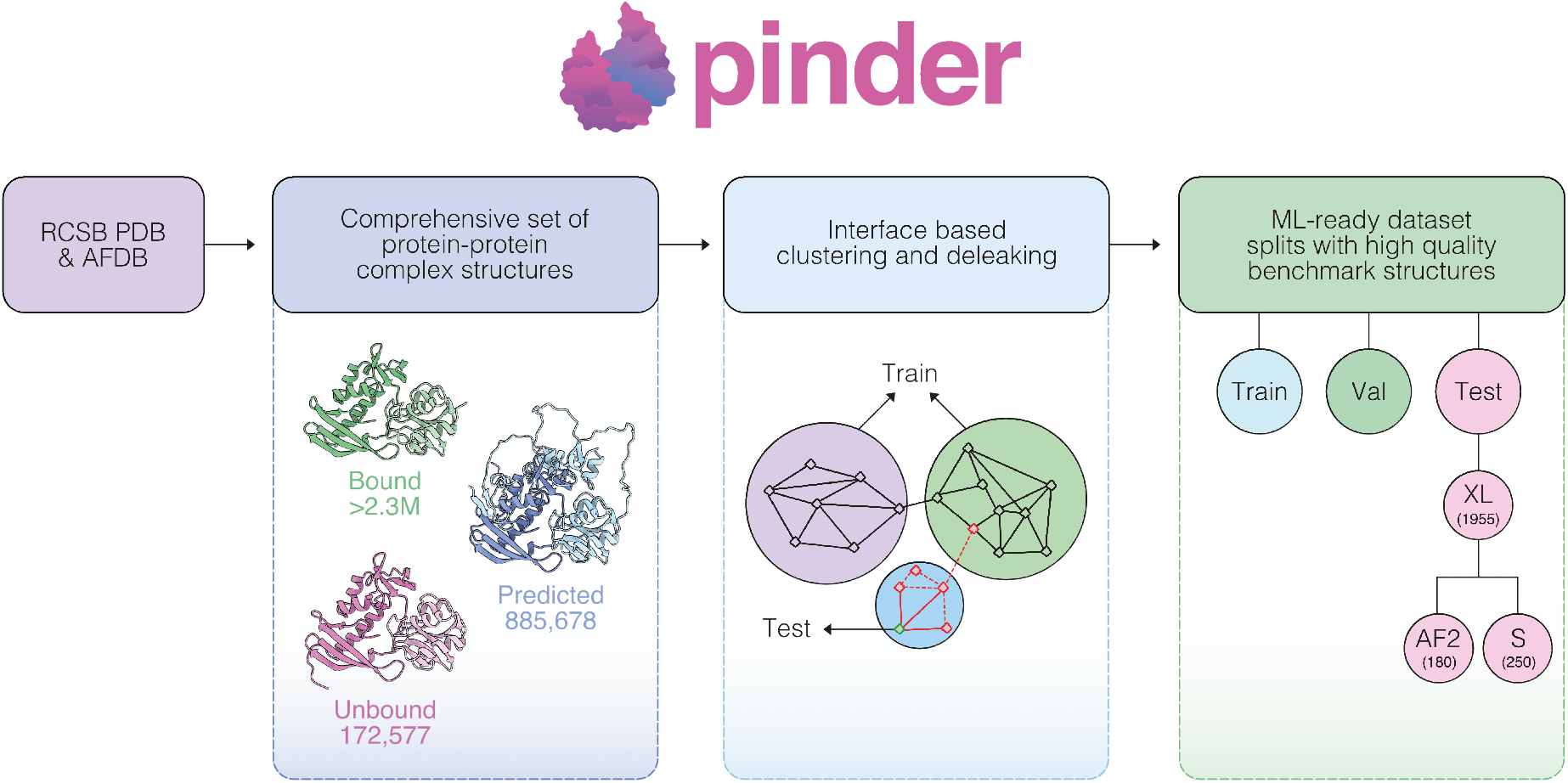
Schematic overview of PINDER. It provides ML-ready dataset with splits tailored to real-world use cases with high-quality benchmark structures.

Contrary to previous benchmarking efforts, such as the Database of Interacting Protein Structures (DIPS-Plus) (Morehead et al., 2023), ProteinFlow (Bio, 2023), DIPS-Plus based EquiDock splits (Ganea et al., 2021) and PPIRef (Bushuiev et al., 2023), PINDER mitigates potential leakage between training and test sets by utilizing structure and sequence similarity-based clustering, specifically targeting interface residues (Figure 2A). Moreover, PINDER emphasizes testing performance on a diverse set of physiological dimers. We found that PINDER’s interface-based splitting method, which uses structural similarity, is superior to the splits obtained using sequence similarity. We demonstrate this by training the state-of-the-art docking model DiffDock-PP on both splits. The model trained on the sequence split overestimated performance on a test set with information leakage and underperformed on a de-leaked test set, while the model trained on the structure split did not. Additionally, the PINDER benchmark revealed that the performance of AlphaFold-Multimer significantly drops when evaluated on a de-leaked test set, further highlighting the importance of using interface-based splitting for accurate benchmarking.

**Figure 2.**
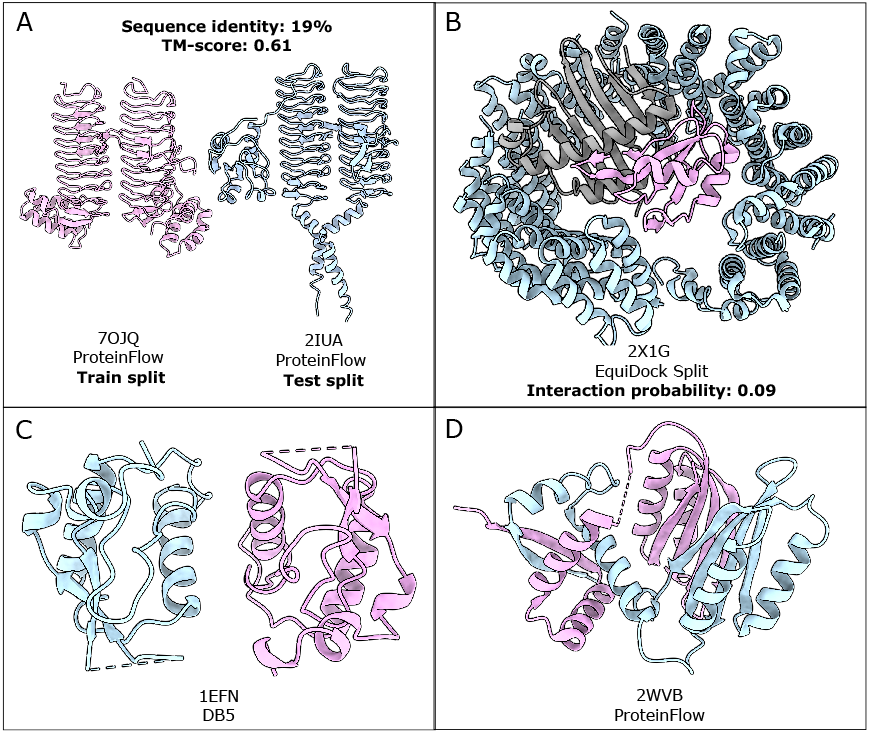
Visual illustrations of selected problematic test systems from existing datasets which are filtered out by PINDER’s benchmark quality criteria. (A) Structural leakage: Test structure example 2IUA from ProteinFlow falls into the same PINDER cluster as a different structure, 7OJQ, from the train split. This is due to the use of sequence identity threshold (40%) in ProteinFlow’s de-leaking workflow. (B) Spurious interaction: Trimeric structure (2X1G) in EquiDock test split contains interaction site shared by three chains. The PPI structure used as the test structure corresponds to the blue and pink chains, however the shared interface arrangement leads to an “incomplete” binding mode between the two chains, leading to a low predicted biological interaction probability (p = 0.09) by PRODIGY-cryst. (C) Interfacial gap: Example of a structure (1EFN) from DB5 which contains gaps within the interfacial area. Detached parts: A structure (2WVB) from ProteinFlow which contains a chain (pink) detached into two parts.

**Figure 3.**
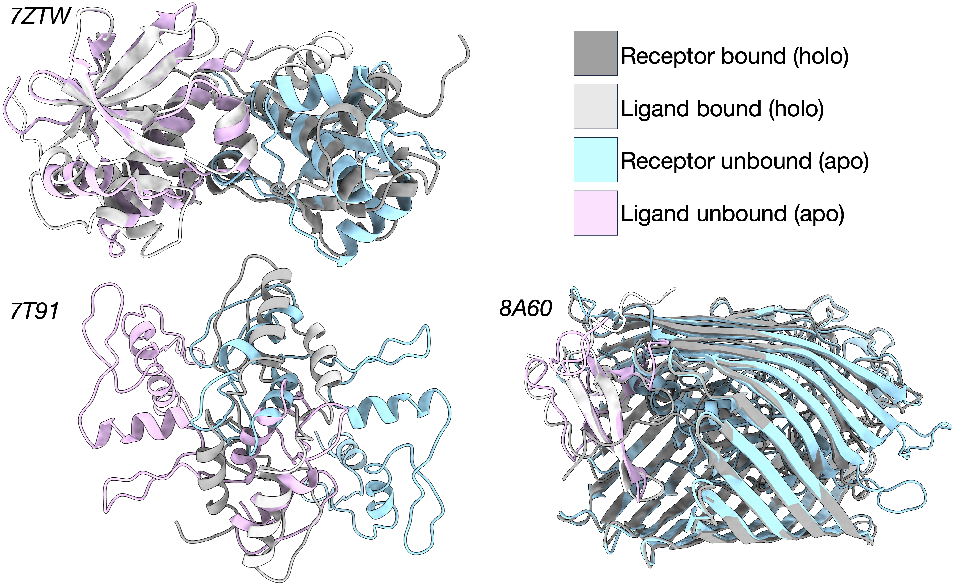
Examples of *apo* structures from the PINDER-AF2 set superimposed on the ground truth structures. All these examples are categorized as “flexible” cases, with no docking tools able to predict with Acceptable or higher quality (more details on flexibility in Figure 10).

## 2. Methods

### 2.1. Data and code availability

The first release of PINDER database was constructed by downloading all assemblies from the RCSB NextGen database (as of 01.29.2024, (Choudhary et al., 2023)). The NextGen database was developed to centralize and streamline access to 3D protein structures from the PDB with enriched structural annotations. PINDER is end-to-end reproducible and intended for online periodic updates. PINDER’s source code is available at https://github.com/pinder-org/pinder as a Python library.

### 2.2. Dataset processing and generation

After the mmCIF files were obtained from NextGen, the first biological assembly was generated. As a result of applying crystallographic symmetry operators, some chains were removed and some added. To resolve chain naming in copies of chains, we added the entity ID as a suffix to each mmCIF chain name, e.g. *A 1, A 2*.

#### *holo* (bound) structures

Binary protein-protein interactions (PPIs) were identified between pairs of chains in a biological assembly with at least one pair of backbone atoms (C, CA, N, O) in contact at a 10 Å threshold. In each binary PPI, the chains were assigned names R (receptor) and L (ligand); conventionally, the longer chain is assigned as the receptor. After the binary decomposition of each assembly, we obtained a total dataset comprising 2,319,564 *holo* PPI systems.

A subset of the most relevant annotations retained or calculated for these assemblies in PINDER is described below:

1. ***RCSB-derived annotations***, includes identification tags, oligomeric state, structure determination method, resolution, bio-assembly, chain information and ECOD domain annotations.
2. ***Gap-proximal interfacial atoms and residues***, defined as the number of interface atoms within a given radius (4 Å or 8 Å) of one of the residue gaps, i.e. residues missing in the determined structure.
3. ***Bio-relevant/crystal contact propensity***, defines the likelihood that an interface is a crystal contact, annotated using PRODIGY-cryst (Jiménez-García et al., 2019).
4. ***Planarity***, defined as deviation of interfacial C*α* atoms from the fitted plane. This quantifies interfacial shape complementarity. Transient complexes tend to have smaller and more planar interfaces (Goncearenco et al., 2015).
5. ***Interface residues***, indices of residues at the interface
6. ***Buried surface area***, defined as solvent accessible surface area change upon binding. (Jiménez-García et al., 2019; Elez et al., 2018).

#### *apo* (unbound) structures

To identify corresponding *apo* structures, all monomeric structures, i.e., chains without any protein interactions, matching the following criteria were paired with *holo* systems (pairs of chains) based on their UniProt accession:

- the number of structure atom types in a monomer *n*_atom_ ≥ 3, to exclude backbone-only structures.
- the number of residues in a monomer *n*_res_ ≥ 5, to exclude short peptides.

Furthermore, we evaluated each potential pairing between a dimer chain and an *apo* monomer. Pairings such as *apoR-holoL* or *holoR-apoL* are assessed against the *holo dimer* (*holoR-holoL*) using several metrics:

- The number of resolved residues must be at least 30% of the number of residues in the corresponding *holo* chain.
- Missing fraction of interface residues for both receptor and ligand chains (ℱ_miss,*R*_ and ℱ_miss,*L*_) should be ≤ 0.3 (F_miss,*max*_).
- RMSD after structural refinement should be below the maximum threshold: *RMSD*_refine_ ≤ 10*Å*.
- The fraction of structurally aligned residues after out-lier rejection 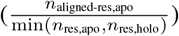 should be ≥ 70% and have at least 30% sequence identity.

The final canonical pairing is determined using a scaled score that equally weights interface RMSD, refinement RMSD, sequence identity, and fractions of native and non-native contacts (*F*_nat_ and *F*_nonnat_). The *apo* monomer with the highest score is selected. To minimize false positives in domain coverage, we calculate the ratios of *apo* to *holo* interface fractions for both ligand and receptor interfaces relative to the native *holo* dimer. These ratios help compare interfacial residues post-superposition to those in the native *holo* structure.

After defining the training split, we repeat the *apo* pairing algorithm for systems in the training split without a paired unbound structure, with a set of assessment thresholds relaxed by a factor of 2.5. For example, the 30% threshold became 12%. This increases the coverage of unbound structures for training while introducing a degree of “noise”, which can be useful for evaluating methods that can learn to ignore information. Dimers which have matches at these thresholds are assigned a “low” *apo* quality label. Overall, the procedure resulted in pairing 172,577 *holo* systems with a corresponding *apo* system and 335,709 *holo* systems with at least one chain paired to an *apo* monomer. A total of 47,717 unique *apo* monomer structures are included in the dataset.

#### Predicted structures

The AlphaFold Protein Structure Database (AFDB) provides structures for over 200 million proteins predicted with the highly successful AlphaFold2 (AF2) (Varadi et al., 2022; Jumper et al., 2021). The predictions in AFDB version 4 cover a majority of UniProt (21/04 release, (Consortium, 2019)), and are uniquely identifiable via the UniProt accession. Similar to the matching of *apo* with *holo* structures, we identified and superposed AFDB entries with the same UniProt accession as the *holo* PPI entries. This resulted in pairing 885,678 *holo* systems with a corresponding predicted system and 968,610 *holo* systems with at least one chain paired to a predicted monomer. A total of 42,827 unique AFDB monomer structures are included in the dataset.

#### Combinatorial structures

In restricted data spaces, data augmentation is critical to providing as much variability during training as possible. Therefore, PINDER is designed to switch between any available *apo* and predicted monomers while sampling the training dataset. PINDER expands the total training dataset to a size of 25 million PPI structures when all combinations of available alternative monomers are used.

#### Conformational changes upon binding

We annotate the flexibility of *apo* and predicted structures corresponding to the reference *holo* complex using the following criteria, similarly to how “difficulty” has been defined in DB4 (Hwang et al., 2010):

- “Rigid-body” if *iRMSD* ≤ 1.5*Å* and the fraction of non-native contacts *f*_nonnat_ ≤ 0.4.
- “Medium” if either of the following conditions is true: 1.5*Å* ≤ iRMSD ≤ 2.2*Å*, (b) iRMSD ≤ 1.5*Å* and *f*_nonnat_ ≥ 0.4.
- “Flexible” if none of the above conditions are true.

#### Interface clustering

To ensure diversification and eliminate redundancy, we employ an interface clustering scheme. As a pre-processing step, systems with fewer than 7 interface residues are excluded from consideration. Then, we computed all-vs-all alignments between all available chains using Foldseek (van Kempen et al., 2023). After extracting the scores and alignment indices, we construct an alignment graph in which each node represents a unique chain and each edge stores the local Distance Difference Test score (lDDT) between the matching chains, along with their respective start and end alignment indices. This graph is pruned by applying a mean lDDT threshold of 0.7, filtering edges to consider only structurally similar chains. Additionally, we remove edges where the alignment indices do not overlap with at least 50% of either chain’s interface residues, thus limiting the alignments to those with higher interface overlap.

Community clustering is then performed on the filtered graph using the label propagation algorithm to obtain cluster IDs for each chain. Chains with less than 40 residues are considered “peptides” and assigned to cluster ID “p”. The paired cluster IDs for each PPI system are referred to as PINDER clusters. These clusters are essential for sampling unique PPI interfaces, generating splits that maximize structural diversity and minimizing redundancy and data representation bias inherent to the RCSB database.

For the selection of high-quality dimer test systems, each PINDER system is initially labeled as *proto-test* if it meets our dimer quality criteria (see Appendix A.10.4). We then move the *proto-test* to the “leakage removal” step.

### 2.3. Split generation and leakage removal

The splitting algorithm for PINDER (Algorithm 1) is designed to achieve diversification, eliminate redundancy, prevent data leakage, and maximize the quality of the test and validation datasets.

Our splitting algorithm is configurable by a set of graphs *G*, here the Foldseek graph with an lDDT threshold of 0.55 and the MMseqs2 graph with a sequence identity threshold of 30%, and a neighbor depth (*D*) determining the maximum length of the shortest path between two systems to constitute leakage (here set to 2 for both graphs). Both the relatively low Foldseek threshold and the transitively captured hits are used to maximize recall, thereby limiting leakage. The minimum neighbors *m* helps to avoid singletons or sparsely connected, potentially unrealistic systems in the test set, while the maximum neighbors *M* = 1000 puts a cap on the number of systems removed from the train set by one test system, to maintain an acceptable training set size.

These clusters containing *proto-test* systems are then sorted by heterodimer state, availability of *apo* and predicted models, and by their release date, prioritizing the most recent. We then remove the redundancy of the *proto-test* by limiting it to the top *n* (*n* = 1) systems from each cluster. The obtained *proto-test* set is further randomly split into *test* and *validation* sets with equal fractions. Finally, “de-leaked” PPIs are masked out from the train set by masking all cluster members and transitive hits found at the de-leaking stage.

#### Algorithm 1

Splitting

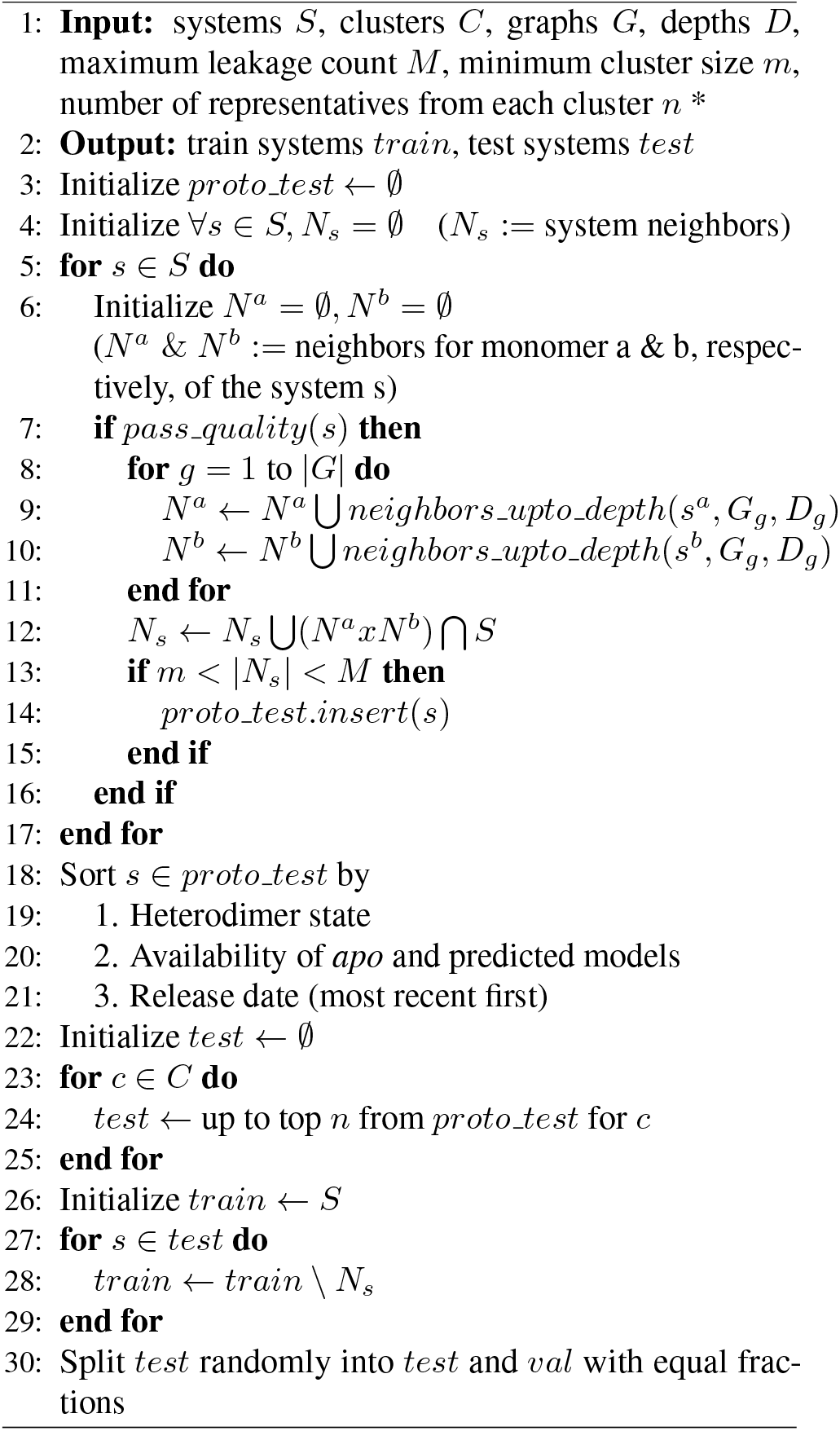

#### Test subsets

The total test set derived from the PINDER split generation process contains 1,955 dimers and is designated as the PINDER-XL benchmark set. From the PINDER-XL set, we sample a smaller subset of 250 test systems to form the PINDER-S benchmark, for ease of evaluation in computationally expensive inference settings. The selection process is biased towards systems with available *apo* structures, heterodimers, and diversity at the family level according to ECOD classification as well as UniProt accession diversity. Finally, the PINDER-AF2 dataset specifically targets systems added after the training cutoff date for AlphaFold-Multimer and is designed to evaluate methods against AlphaFold-Multimer.

After defining the initial splits, we further evaluate similarity of interfaces in the test and validation splits with respect to members of the training set using interface alignment metrics derived from iAlign (Gao & Skolnick, 2010). Any pair of dimers coming from different splits with at least one structural or sequence hit from Foldseek or MMseqs2 is assessed for interface similarity.

Interfaces are marked as similar when all of the following criteria are met:

- iRMS < 5.0.
- IS-score > 0.3.
- log(*P* -value) < −9.0.

Where iRMS is the interface RMSD after alignment, IS-score is the Interface Similarity score, and log(*P* -value) is the logarithm of the statistical significance of the IS-score as determined from the distribution of interface scores in the iAlign methodology (Gao & Skolnick, 2010).

For PINDER splits, the train split is the same as defined by the PINDER splitting methodology. For PINDER-AF2, members released after the training cutoff date are evaluated with respect to any systems released prior to the training cutoff date. Systems not meeting the interface similarity criterion are removed from the holdout set, implying that they have similar counterparts within the AlphaFold-Multimer training dataset. The final PINDER-AF2 subset was defined as the set of time-split members from PINDER-XL (180) with no similar interfaces in the train split at an iAlign log(*P* -value) threshold of −9.0. The PINDER-AF2 test set can be leveraged to evaluate and compare docking method performance against AlphaFold-Multimer, without a need for resource-intensive retraining. For PINDER-XL, the same process is applied, but any interfaces determined to be similar to the training set are left in the test subset and instead marked with a label to use as a quality control measure and for the stratification of performance. The selected thresholds, final selection of test subsets, and the impact of removing similar interfaces as determined by iAlign is described in Appendix A.4.1.

### 2.4. Evaluation

PINDER evaluation is performed for each predicted docking pose. We compute the following scores and metrics with respect to the reference ground truth systems:

- LRMS - Ligand Root Mean Square deviation (RMSD) over backbone atoms of the shorter chain (ligand) after superposition of the longer chain (receptor).
- iRMS - Interface RMSD over backbone atoms of receptor-ligand interface residues in the target (native) after superposition on their equivalents in the predicted complex (model). Interface residues are defined at a 10 A° atomic contact cutoff.
- Fnat - Fraction of native interfacial contacts preserved in the interface of the predicted complex. Interfacial contacts are pairs of heavy atoms from receptor and ligand within 5 Å.
- DockQ - Combines Fnat, LRMS, and iRMSD into one score:

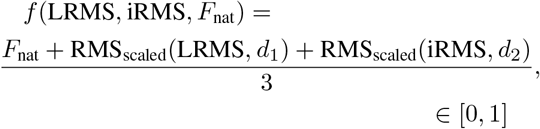

Where RMSscaled is defined as:

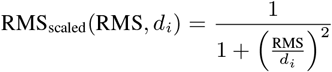

RMS_scaled_ represents the scaled RMS deviations for LRMS or iRMS, and *d*_*i*_ is a scaling factor (*d*_1_ = 8.5 Å, *d*_2_ = 1.5 Å).

We segregate model predictions into four quality classes using the CAPRI classification system of High, Medium, Acceptable and Incorrect, defined in Table 1. In order to make a fair comparison of methods across complete test sets, we penalize the missing predictions by registering the missing system as *iRMS* = 100.0, *LRMS* = 100.0, *Fnat* = 0.0, *DockQ* = 0.0, *CAPRI* = *Incorrect*.

**Table 1.** Definition of CAPRI quality classes according to CAPRI measure criteria.

| Ranking | Conditions based on CAPRI measures |
| --- | --- |
| High | $F_{\text{nat}} \geq 0.5$ AND $\text{LRMS} \leq 1.0$ OR $\text{iRMS} \leq 1.0$ |
| Medium | $(F_{\text{nat}} \geq 0.3 \text{ AND } F_{\text{nat}} < 0.5)$ AND $(\text{LRMS} \leq 5.0 \text{ OR } \text{iRMS} \leq 2.0)$ |
| Acceptable | $(F_{\text{nat}} \geq 0.1 \text{ AND } F_{\text{nat}} < 0.3)$ AND $(\text{LRMS} \leq 10.0 \text{ OR } \text{iRMS} \leq 4.0)$ |
| Incorrect | $F_{\text{nat}} < 0.1$ OR $(\text{LRMS} > 10.0 \text{ AND } \text{iRMS} > 4.0)$ |

For each docking tool, predictions are categorized based on the input monomer types (*holo, apo*, or predicted monomers). The evaluation output is then at the individual system level, with multiple poses, allowing for comprehensive assessment of individual system performance as well as model’s ranking capability. Additionally, the output is grouped by the test subsets (PINDER-XL, PINDER-AF2 and PINDER-S), flexibility difficulty, and monomer type to provide further insights into method performance across them.

## 3. Results & Discussion

### 3.1. pinder is a large and diverse PPI dataset

PINDER is the largest and most diverse structural PPI dataset to date, containing over 2 million PPIs derived from the RCSB NextGen database, and expanded into more than 25 million distinct PPI structures by PINDER’s augmentation dataloader (see section A.5 for details). As shown in Table 8, this is substantially larger than existing datasets like DIPS-Plus, ProteinFlow, PPIRef (Bushuiev et al., 2023), and EquiDock splits, which contain 42k, 169k, 837K, and 42k PPIs, respectively. The diversity of PINDER is evident in the distribution of its metadata annotations (Figure A.41), which cover a broad range of residue and chain lengths, buried surface areas, and interface characteristics. This diversity ensures that models trained on PINDER are exposed to a wide range of PPI examples, improving their ability to generalize to new and unseen interactions. Additionally, the test set was constructed by sampling a single high quality representative from each interface cluster to maximize the diversity of test interfaces (see Appendix Section A.2), specifically for the dimer modeling task.

### 3.2. pinder provides multiple starting conformations for realistic PPI docking

PINDER provides multiple starting conformations for each PPI, enabling a more realistic evaluation of protein docking methods. Unique to PINDER, 136,498 *apo* PPIs and 566,171 predicted PPIs are also included in the training set, allowing methods to learn docking under more diverse and challenging conditions, such as when the *apo* structures of the interacting proteins are known or when only predicted structures are available. The availability of multiple conformations also facilitates the development of docking methods that can leverage information from both *holo* and *apo* structures to improve prediction accuracy. Additionally, the PINDER test set includes 342 *apo*, in comparison to 257 in DB5.5, and more than 1,747 predicted structures unique to PINDER with a balanced range of difficulties (shown in Fig. A.43), allowing for a more comprehensive assessment of performance.

### 3.3. pinder splits by interface similarity

PINDER employs a novel splitting algorithm that ensures the diversification of test and validation sets, while preventing leakage between them and the large train set. In addition, it combines both structure (Foldseek) and sequence (MM-seqs2) similarity metrics and leverages similarity graphs to maximize recall. The interface-based splitting approach also allows for a more targeted evaluation of docking methods on specific types of interfaces, such as those with multiple modes of binding. Several quality control (QC) measures were implemented, listed in the Table A.1, to assess the diversity of the splits in terms of interface clusters, ECOD (Evolutionary Classification of protein Domains) annotations, and Pfam (Protein families database) clans, as well as the potential leakage between the splits based on UniProt pairs, ECOD pairs, and global sequence similarity.

Unlike other datasets that rely solely on either sequence similarity (DIPS, ProteinFlow) or structure similarity (PPIRef) for splitting, PINDER’s approach ensures that the test and validation sets contain interfaces with almost no similar examples present in the training set according to orthogonal quality control metrics. This reduced risk of information leakage leads to a more realistic evaluation. While PPIRef also clusters protein interfaces, we demonstrate that it is insufficient to prevent leakage across the splits. Specifically, PINDER exhibits notably lower ECOD pair leakage (6.34%) compared to DIPS (63.94%), ProteinFlow (26.07%), and PPIRef (45.17%), indicating that PINDER ‘s test and validation sets share fewer previously annotated domain pairs with its training set. In addition to preventing leakage, by clustering proteins based on interface similarity and selecting a single representative from each cluster for the test set, PINDER maximizes the diversity of test interfaces while minimizing redundancy (see Table A.1 for comparison).

### 3.4. Results on the pinder benchmarks

The PINDER leaderboard demonstrates docking results for three classical (HDOCK, PatchDock, and FRODOCK) and one DL-based (DiffDock-PP) protein-protein docking methods as well as the co-folding method AlphaFold-Multimer, with three separate test sets (PINDER-S, PINDER-XL, and PINDER-AF2), and three sets of systems, namely bound (*holo*), unbound (*apo*) and predicted AlphaFold2 monomers. Note that PINDER-AF2 and PINDER-S are subsets of PINDER-XL, hence only a single training run is required to report metrics on these benchmarks. DiffDock-PP was trained using *holo, apo*, and predicted structures to assess its ability to learn from augmented training data, despite the rigid-body nature of the model. AlphaFold2-Multimer inference produced 5 ranked poses, and Diffdock-PP inferred 40 poses for each test system. All classical docking methods were executed using their default settings (with template mode disabled), see A.7, producing a set of 40 poses ranked intrinsically by the respective methods. Definitions of reference-based evaluation metrics are provided in Methods 2.4.

#### Results on holo, apo, and predicted structures

As a general trend, all tools exhibit their best performance when interfaces have higher solvent-accessible surface area buried at the interaction site (dSASA) (see Appendix Figure A.13 and A.14), reflecting an easier solution with more interaction space. For the task of rigid docking, where the input structures are already in the correct bound conformations, the classical docking methods outperformed the Deep Learning methods by a large margin with HDOCK in the lead (Table 5). However, this is not reflective of performance with information available in most real usage scenarios. When *apo* or predicted structures were used as input, method performance dropped sharply across all methods (Table 5). Interestingly, classical methods outperformed AlphaFold-Multimer on all of the *apo* structures, while AlphaFold-Multimer leads on predicted structures. PINDER is the first dataset to provide paired *apo* and *predicted* structures. In the case of DiffDock-PP, which was trained on PINDER-provided *apo, holo* and *predicted* structures, this yields reasonable performance on medium/flexible systems - reaching approximately half of the performance on rigid-body systems (Figure 10). This underscores the importance of providing these variations during training.

#### Results by chain novelty

PINDER’s interface clustering approach allows us to categorize test systems into three different groups.

- Systems with **both** of the chains novel to test split, where none of the chains are clustered with chains found in train.
- Systems with a **single** novel, and the other chain cluster also present in train.
- Systems with **neither** of the chains unique to test, where both chains are clustered with the chains found in train.

This classification helps identify homologous chains, i.e. structurally similar chains that can, however, be involved in different binding modes. As depicted in Appendix Figures A.11 and A.12, both DL methods (DiffDock-PP and Alphafold-Multimer) show the highest success rates in the category “**both** novel” and the lowest in the category “**neither** novel”. This suggests that these methods may over-fit to the specific binding modes in the training data, failing to generalize to new binding modes for homologous chains in the test data.

#### Clashes

Figures 4 and 5 show the level of clashes in the docking results across PINDER-XL and PINDER-AF2, using the VoroMQA clash score (Olechnovič & Venclovas, 2017), as a complementary non-distance based measure of pose validity. While DiffDock-PP generally has poor scores with many clashes, AlphaFold-Multimer shows good (low) clash scores highlighting the advantages of incorporating flexibility. Note that this is expected, as the rigid-body docking method DiffDock-PP will naturally be trained to have clashing monomers for apo and predicted input structures to minimize distance to the interface defined by monomers.

**Figure 4.**
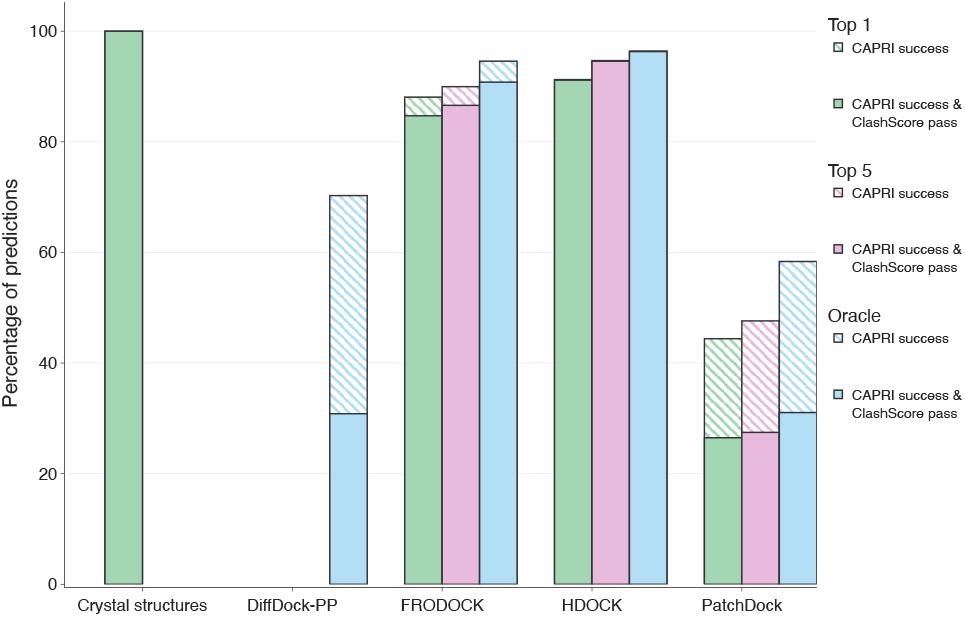
Bar-chart comparing the CAPRI success rates of bench-marked methods on *holo* systems across PINDER-XL, with and without low (< 0.1) VoroMQA clash scores for oracle, top-1 and best of top-5 poses.

**Figure 5.**
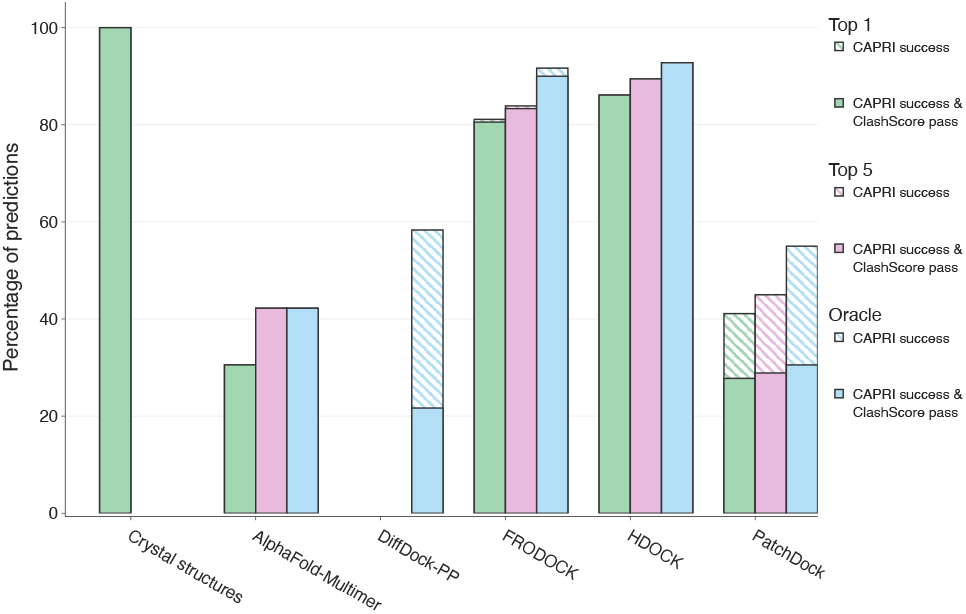
Bar-chart comparing the CAPRI success rates of bench-marked methods on *holo* systems across PINDER-AF2, with and without low (< 0.1) VoroMQA clash scores for oracle, top-1 and best of top-5 poses.

Results for *apo* and predicted input structures, the PINDER-S dataset and distributions of additional clash metrics are reported in Appendix A.3.1.

### 3.5. AlphaFold2-Multimer performance on structurally novel interfaces and on PINDER-AF2

To assess the impact of structural interface information leakage on AlphaFold-Multimer 2.3 performance, predictions were generated for a time-split subset of PINDER-XL consisting of 675 test systems. This time split contains a mixture of different similarity levels to train, allowing us to stratify performance by iAlign similarity to structures deposited before the AlphaFold2-Multimer training date cutoff. Note that due to the quality filtering and the simplification of the problem to the easier dimer (*versus* oligomer) prediction problem, we would expect to see a high CAPRI success rate compared to results originally reported. Indeed, we see a success rate of 80.6% on the 675 time-split systems. However when limiting to structurally novel interfaces, the success rate significantly drops to 57.8% (Figure 6). The final success rate is marginally lower than the reported value of 60.2% on the common subset of dimers evaluated in a previous benchmark (Zhu et al., 2023).

**Figure 6.**
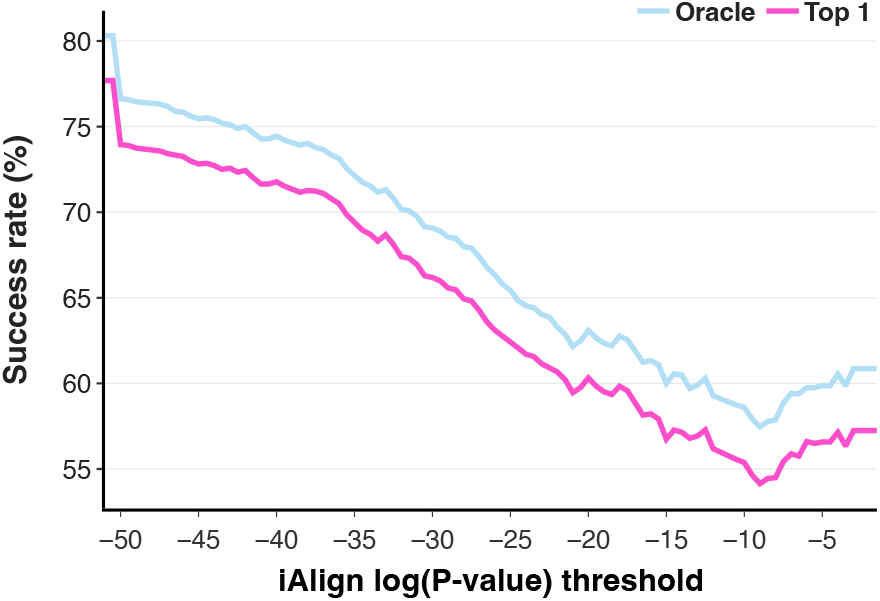
Percentage of AlphaFold-Multimer predictions from a time-split subset of PINDER-XL with CAPRI classification of acceptable or higher with and without de-leaking at varying iAlign log(*P*-value) thresholds. Success rates are reported for both Oracle and top ranked predictions. The figure is truncated to log(*P*-value) = −50, where values outside of the domain are represented by an initial drop from the maximum success rate to the first value in the domain.

Although the total number of systems in PINDER-AF2 with *apo* structures is small, AlphaFold-Multimer’s performance is substantially lower on this subset compared to the *holo* and predicted subsets, despite using identical sequence-only inputs for inference. Several factors likely contribute to this discrepancy.

Figure 9 shows that AlphaFold-Multimer’s performance is generally worse when predicted monomers are unavailable in AFDB. Performance also declines when the predicted biological interaction probability or interface surface area is lower. Furthermore, when provided with the full-length sequence, misfolded monomers can lead to incompatible interfaces due to binding site occlusion, which may not be observed even when using *apo* structures as input (Figure 9D-F).

**Figure 7.**
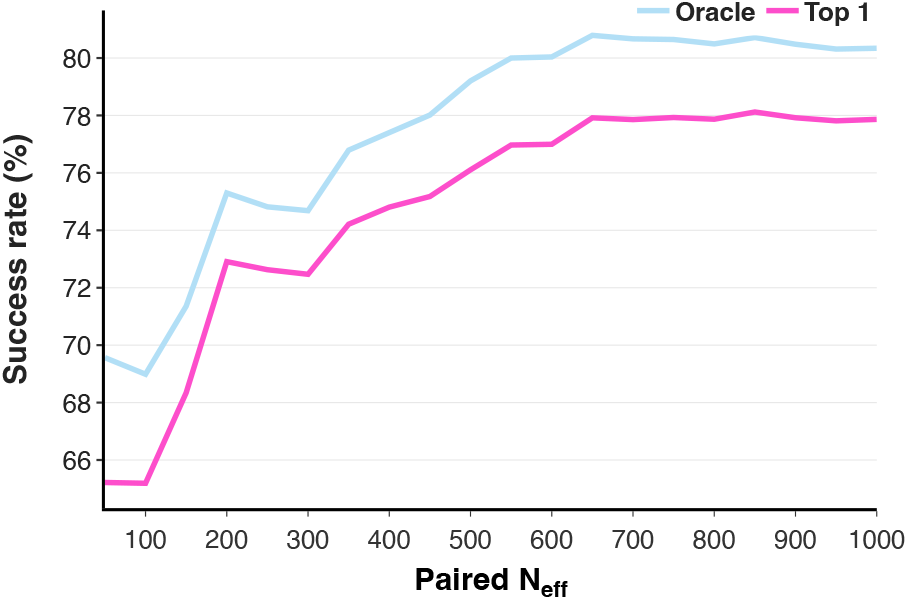
Percentage of AlphaFold-Multimer predictions from a time-split subset of PINDER-XL with CAPRI classification of acceptable or higher at varying thresholds on paired N_eff_. Success rates are reported for both Oracle and top ranked predictions. The figure is truncated to N_eff_ = 1000, where the success rate plateaus.

**Figure 8.**
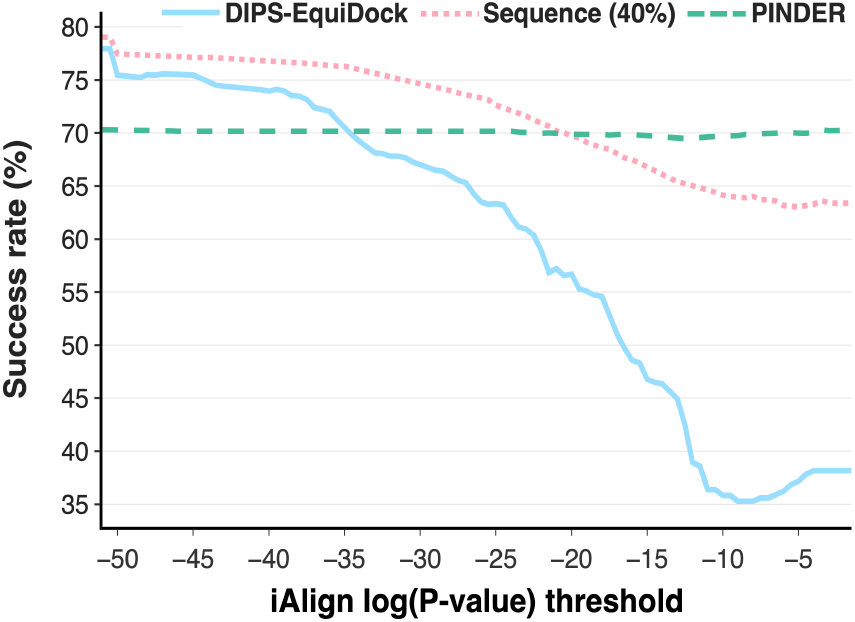
Percentage of DiffDock-PP predictions from models trained on PINDER, sequence and DIPS-EquiDock splits with CAPRI classification of acceptable or higher with and without de-leaking at varying iAlign log(P-value) thresholds. Success rates are reported for Oracle predictions.

**Figure 9.**
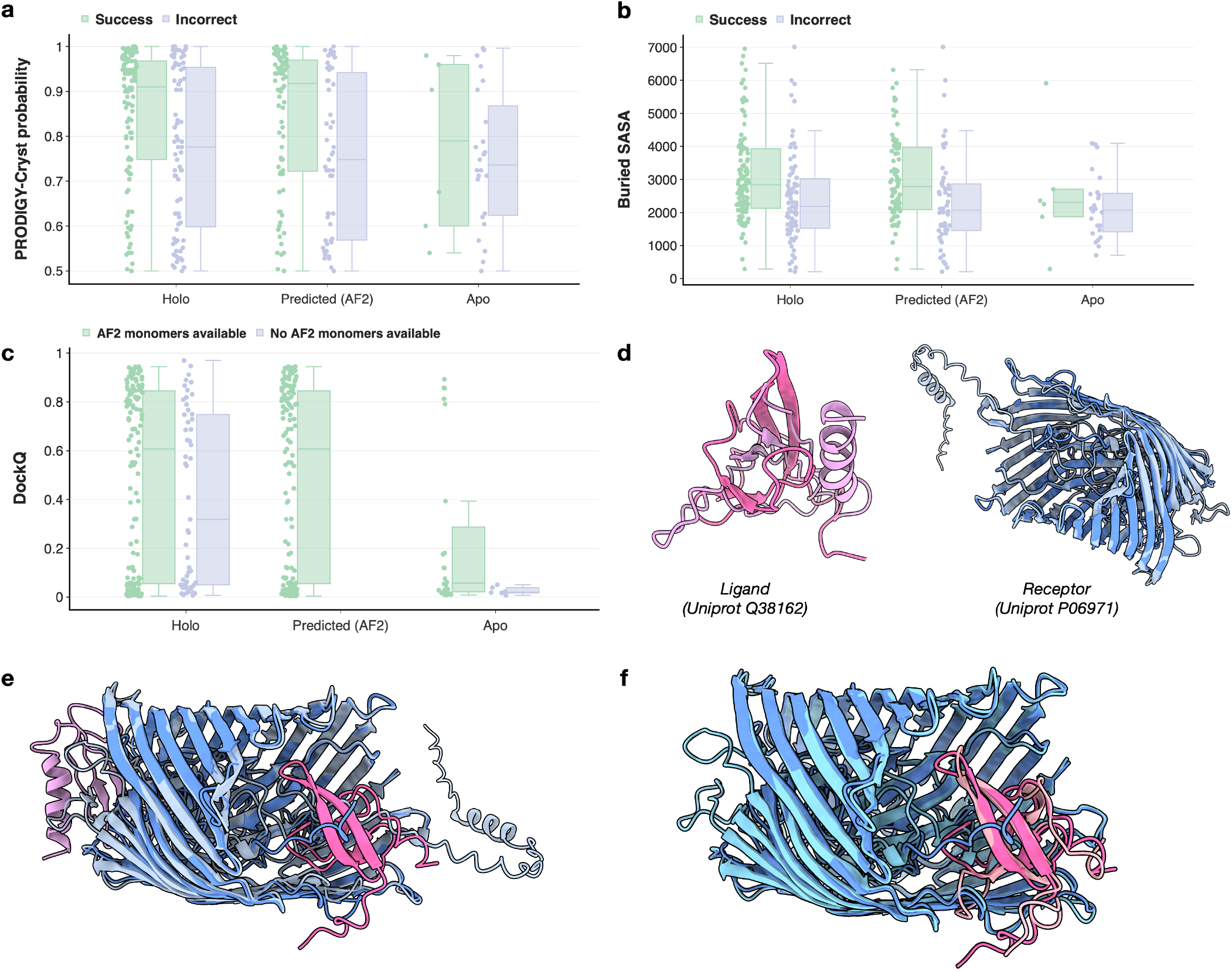
AlphaFold-Multimer performance on PINDER-AF2 from different monomer subsets. Figures **a-b**, property distributions illustrating dataset biases that impact CAPRI success rates in the limited *apo* subset. Points are colored by CAPRI success (green, acceptable or higher; light purple, incorrect) with higher average predicted interaction probability and buried SASA in cases with successful predictions and lower average values in the *apo* subset; **c**, DockQ distribution in each monomer subset of PINDER-AF2 colored by availability of predicted monomers in AFDB; **d**, Incorrect modeling of monomer chains leads to co-folding failure: predicted monomers aligned with ground truth; **e**, incorrectly folded truncated tail (light blue) blocks true binding site (dark pink), leading to wrong docking site (light pink); **f**, *apo* monomers superimposed to bound *holo* dimer illustrating challenging, but more compatible monomers than those predicted by AlphaFold-Multimer.

**Figure 10.**
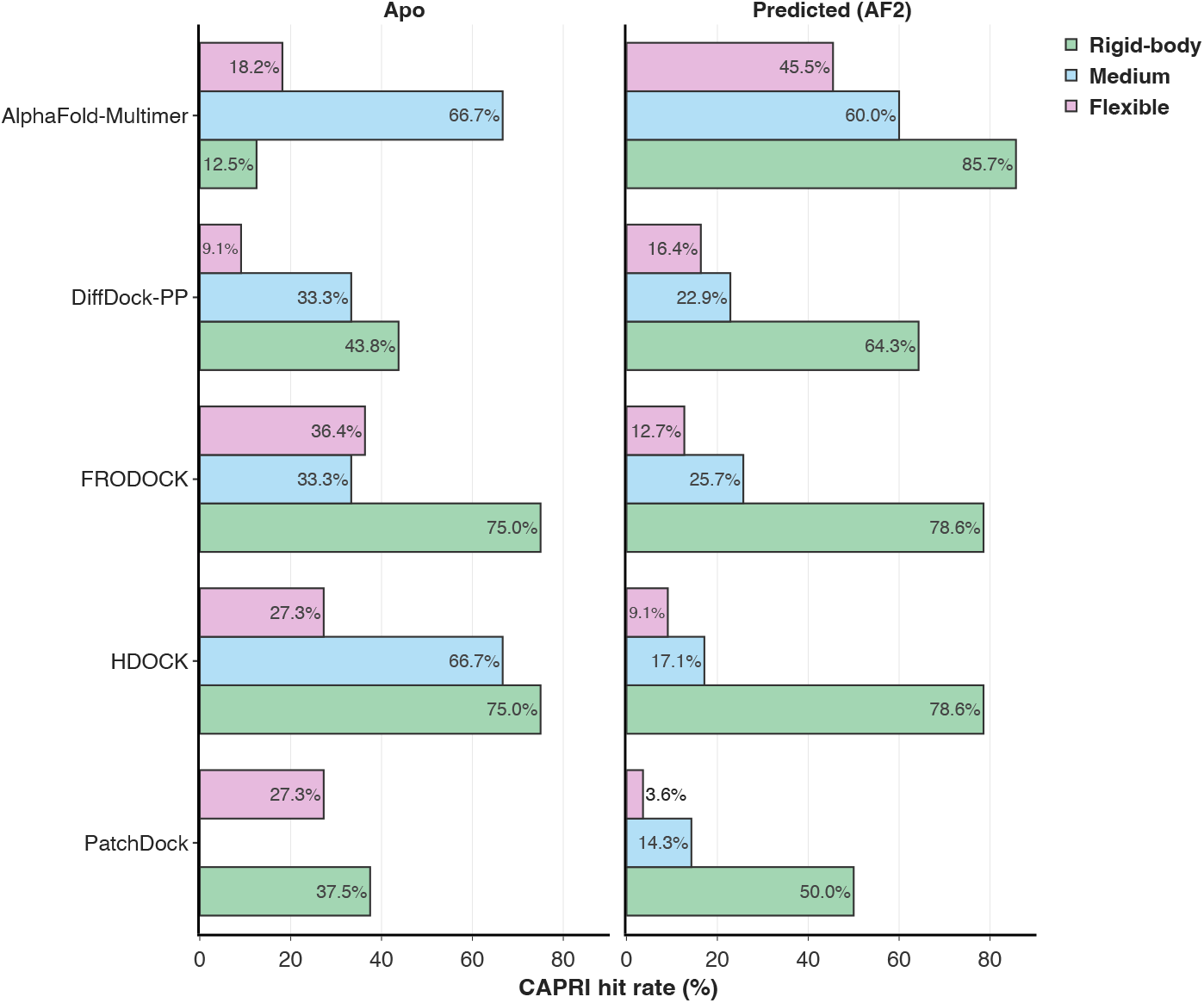
Bar-chart showing oracle CAPRI hit/success rates between benchmarked methods on PINDER-AF2 set using *apo* and predicted input structures at different “unbound” flexibility categories (Rigid-body, Medium and Flexible). Success rates are reported relative to the total number of systems in each flexibility category.

Moreover, due to the time split with interface de-leaking, medium and hard difficulty PINDER-AF2-*apo* interfaces are enriched for cases where AlphaFold2-Multimer was trained exclusively on monomers in *apo* conformations distinct from the holo conformations of these monomers. As shown in Figure A.43, PINDER-AF2-*apo* contains a higher proportion of medium and hard interfaces.

As previously reported (Saldaño et al., 2022), AlphaFold generally struggles to capture multiple conformations for a given protein. Since AlphaFold was specifically trained on monomers in conformations different from the *holo* conformation observed in these cases, it is more likely to predict the wrong *apo* conformation. This incorrect prediction can occlude the binding site, leading to the wrong site being chosen (Figure 9E).

#### Co-folding versus docking approaches

An important limitation of our splitting procedure and AlphaFold2-Multimer results reported here is that often dimers can be similar to interfaces occurring between domains within the monomers (Sprinzak & Margalit, 2001)(Alborzi et al., 2021). Indeed, we found many cases where a dimer was similar to interfaces within monomers, see Figure A.25. Cofolding methods trained on monomers may profit from having access to more supervision relevant for the PPI docking task, thus leading to unaccounted leakage and possibly over-estimated performance (AlphaFold2-Multimer performance in Table 7). While structural delineation of domain-domain interfaces can be considered an effective data-augmentation strategy (Hayes et al., 2024), these interactions are highly redundant and de-leaking of such interfaces leads to a significant increase in computational complexity. While domain-domain interfaces highlight a potential advantage co-folding methods have over docking-based methods, we did not consider this to be leakage for the docking task in this PINDER release.

Further, to avoid retraining DiffDock-PP for PINDER-AF2, we did not include any PINDER-XL clusters that are not part of PINDER-AF2 in the training data. This gave AF2-Multimer a small but tangible data advantage, despite isolating leakage from all models equally.

#### Reliance on evolutionary signals

Since some Deep Learning methods make use of co-evolutionary information, either explicitly by including multiple-sequence alignments (MSA), or implicitly by leveraging pre-trained sequence embeddings as features, they may be able to pick up co-evolutionary signals better than other methods for systems which have more homologous structures. The average number of effective paired chain sequences (*N*_eff_) in the MSA, a measure of PPI MSA depth, is often used to estimate the availability of co-evolutionary signals for a given protein interaction. Different types of protein-protein interactions, including antibody-antigen interactions (Gaudreault et al., 2023) and transient interactions (Mukherjee & Chakrabarti, 2021) tend to have low *N*_eff_ values and thus present considerable challenges in complex modeling and docking performance of methods which rely on sequence conservation and co-evolution signals. Therefore, we assess model performance at varying upper limits on *N*_eff_ as a proxy for the strength of detectable co-evolutionary signals. We show the CAPRI success rates of oracle poses from the time-split subset of PINDER-XL at varying paired *N*_eff_ in Figure 7. AlphaFold-Multimer performs significantly worse on systems with fewer *N*_eff_, i.e. those with hard-to-detect or missing co-evolutionary signals, yet this dependence is less pronounced than the impact of interface novelty.

### 3.6. pinder split allows model to generalize

To elucidate the positive impact of the PINDER split on model generalization, we performed retraining of a State-of-the-Art (SOTA) docking method on different splits. An additional sequence-based splitting method was implemented and the diffusion-based docking model DiffDock-PP (Ketata et al., 2023) was retrained using NVIDIA BioNeMo (bio) with both of these additional splits next to the PINDER split for a head-to-head comparison. The clustering for the sequence split was performed using an MMseqs2 cutoff of 40% sequence identity. In order to reduce the training time, we used PINDER’s filtering utilities to first remove structures which didn’t meet certain quality criteria (Appendix A.6.1), then we ranked cluster members in each cluster in terms of *apo* and predicted monomer availability, and finally sampled 3 systems from each cluster when available. In addition to the PINDER-derived (structure and sequence) splits, DiffDock-PP was re-trained from scratch and evaluated using the DIPS-Plus EquiDock splits used in the original paper (Ganea et al., 2021). The training processes on all three splits were stopped when the validation loss plateaued. Table 2 shows the number of systems in the train, val, and test sets for all splits. Tables 3 and 4 show percentage of test and validation systems that are leaky with respect to the training set for each of the splitting methods, as determined by the UniProt, ECOD and iAlign criteria.

**Table 2.** Split Sizes of Various DiffDock-PP Datasets. Cluster counts are reported based on PINDER structural clusters.

| Split method | Test | Val | Train |  |
| --- | --- | --- | --- | --- |
|  |  |  | Dimers | Clusters |
| Structure (PINDER-XL) | 1955 | 1958 | 53628 | 24689 |
| Sequence (40%) | 2536 | 2538 | 68245 | 24830 |
| DIPS-EquiDock | 757 | 745 | 30938 | 3899 |

**Table 3.** Test split UniProt, ECOD and interface leakage from different splitting methods.

| Split method | UniProt | ECOD | iAlign |
| --- | --- | --- | --- |
| Structure (PINDER-XL) | 1.74 | 5.42 | 12.69 |
| Sequence (40%) | 1.62 | 47.91 | 58.36 |
| DIPS-EquiDock | 42.93 | 63.94 | 83.36 |

**Table 4.** Validation split UniProt, ECOD and interface leakage to training set from different splitting methods.

| Split method | UniProt | ECOD | iAlign |
| --- | --- | --- | --- |
| Structure (PINDER-XL) | 2.04 | 7.51 | 9.45 |
| Sequence (40%) | 1.97 | 51.14 | 52.92 |
| DIPS-EquiDock | 47.45 | 75.54 | 84.03 |

**Table 5.** DockQ CAPRI classification evaluation metrics for the PINDER-XL test set across four evaluated docking methods. The leftmost column shows the input type (*holo*/*apo*/predicted) along with the number of evaluated systems. Methods are ranked alphabetically, results for the highest performing method are highlighted as bold. The rightmost column shows the number of systems not predicted by the respective method.

| PINDER-XL |  | DockQ CAPRI classification |  |  |  |  |  |  |  |  | Miss. Sys. |
| --- | --- | --- | --- | --- | --- | --- | --- | --- | --- | --- | --- |
|  |  | Top-1 |  |  | Top-5 |  |  | Oracle |  |  |  |
|  |  | Acceptable | Medium | High | Acceptable | Medium | High | Acceptable | Medium | High |  |
| Holo<br>(1,955) | DiffDock-PP | * | * | * | * | * | * | 70.74 | 50.95 | 23.38 | 0 |
|  | FRODOCK | 88.08 | 86.96 | 82.71 | 91.82 | 91.15 | 87.77 | 94.58 | 93.61 | 90.23 | 1 |
|  | HDOCK | **91.25** | **90.9** | **90.18** | **94.68** | **94.32** | **93.35** | **96.42** | **95.91** | **94.78** | 0 |
|  | PatchDock | 56.68 | 55.19 | 46.6 | 65.52 | 63.53 | 52.63 | 75.86 | 71.76 | 58.52 | 0 |
| Apo<br>(342) | DiffDock-PP | * | * | * | * | * | * | 41.52 | 21.05 | 4.39 | 0 |
|  | FRODOCK | 43.27 | 39.77 | 29.82 | **53.51** | **48.54** | 35.09 | **61.4** | **54.09** | 37.43 | 0 |
|  | HDOCK | **45.32** | **42.4** | **35.38** | 51.46 | 47.66 | **38.3** | 61.11 | 53.51 | **40.94** | 0 |
|  | PatchDock | 19.59 | 17.25 | 11.99 | 26.32 | 22.22 | 12.57 | 37.13 | 30.12 | 14.04 | 0 |
| Predicted<br>(1,747) | DiffDock-PP | * | * | * | * | * | * | 50.09 | 33.49 | 6.47 | 2 |
|  | FRODOCK | **49.34** | **44.71** | 20.89 | **54.89** | **50.2** | 23.7 | **59.59** | **54.32** | 24.96 | 1 |
|  | HDOCK | 48.25 | 44.02 | **26.33** | 53.86 | 49.17 | **28.85** | 58.79 | 52.89 | **29.59** | 0 |
|  | PatchDock | 28.05 | 25.3 | 10.07 | 34.97 | 31.08 | 12.25 | 44.42 | 38.29 | 13.74 | 1 |

**Table 6.** DockQ CAPRI classification evaluation metrics for the PINDER-S test set across four evaluated docking methods. The leftmost column shows the input type (*holo*/*apo*/predicted) along with the number of evaluated systems. Methods are ranked alphabetically, results for the highest performing method are highlighted as bold. The rightmost column shows the number of systems not predicted by the respective method.

| PINDER-S |  | DockQ CAPRI classification |  |  |  |  |  |  |  |  | Miss. Sys. |
| --- | --- | --- | --- | --- | --- | --- | --- | --- | --- | --- | --- |
|  |  | Top-1 |  |  | Top-5 |  |  | Oracle |  |  |  |
|  |  | Acceptable | Medium | High | Acceptable | Medium | High | Acceptable | Medium | High |  |
| Holo<br>(250) | DiffDock-PP | * | * | * | * | * | * | 60.0 | 38.0 | 10.0 | 0 |
|  | FRODOCK | 89.6 | 87.6 | 84.8 | 92.4 | 91.2 | 89.2 | 95.6 | 94.0 | 92.4 | 0 |
|  | HDOCK | **94.4** | **94.0** | **93.6** | **97.2** | **97.2** | **96.4** | **98.0** | **97.6** | **96.4** | 0 |
|  | PatchDock | 53.6 | 52.4 | 43.6 | 65.2 | 62.4 | 52.0 | 76.4 | 74.4 | 61.2 | 0 |
| Apo<br>(93) | DiffDock-PP | * | * | * | * | * | * | 36.56 | 18.28 | 1.08 | 0 |
|  | FRODOCK | 41.94 | 39.78 | 31.18 | **51.61** | **45.16** | 35.48 | **63.44** | 52.69 | 35.48 | 0 |
|  | HDOCK | **44.09** | **40.86** | **33.33** | 47.31 | 44.09 | **36.56** | 59.14 | **53.76** | **38.71** | 0 |
|  | PatchDock | 17.2 | 15.05 | 10.75 | 22.58 | 18.28 | 10.75 | 39.78 | 30.11 | 11.83 | 0 |
| Predicted<br>(250) | DiffDock-PP | * | * | * | * | * | * | 24.8 | 14.0 | 2.0 | 0 |
|  | FRODOCK | **24.8** | **21.6** | 8.0 | **30.4** | **26.8** | 9.6 | **39.6** | **32.4** | 11.6 | 0 |
|  | HDOCK | 23.2 | 19.6 | **11.2** | 28.4 | 25.6 | **13.2** | 35.6 | 30.4 | **14.4** | 0 |
|  | PatchDock | 9.6 | 9.2 | 3.6 | 12.4 | 11.6 | 4.8 | 22.4 | 17.2 | 5.2 | 0 |

**Table 7.** DockQ CAPRI classification evaluation metrics for the PINDER-AF2 test set across five evaluated docking methods. The leftmost column shows the input type (*holo*/*apo*/predicted) along with the number of evaluated systems. Methods are ranked alphabetically, results for the highest performing method are highlighted as bold. The rightmost column shows the number of systems not predicted by the respective method.

| PINDER-AF2 |  | DockQ CAPRI classification |  |  |  |  |  |  |  |  | Miss. Sys. |
| --- | --- | --- | --- | --- | --- | --- | --- | --- | --- | --- | --- |
| Input | Method | Top-1 |  |  | Top-5 |  |  | Oracle |  |  |  |
|  |  | Acceptable | Medium | High | Acceptable | Medium | High | Acceptable | Medium | High |  |
| Holo<br>(180) | AlphaFold-Multimer | 54.44 | 48.89 | 23.89 | 57.78 | 51.11 | 28.33 | 57.78 | 51.11 | 28.33 | 0 |
|  | DiffDock-PP | * | * | * | * | * | * | 58.33 | 37.78 | 15.56 | 0 |
|  | FRODOCK | 81.11 | 80.56 | 75.56 | 86.67 | 86.11 | 83.33 | 91.67 | 89.44 | 86.67 | 0 |
|  | HDOCK | **86.11** | **85.56** | **85.0** | **89.44** | **88.89** | **88.33** | **92.78** | **91.67** | **90.0** | 0 |
|  | PatchDock | 46.11 | 45.56 | 36.11 | 55.0 | 53.33 | 43.33 | 67.78 | 62.22 | 47.22 | 0 |
| Apo<br>(30) | AlphaFold-Multimer | 20.0 | 16.67 | 16.67 | 20.0 | 16.67 | 16.67 | 20.0 | 16.67 | 16.67 | 0 |
|  | DiffDock-PP | * | * | * | * | * | * | 30.0 | 10.0 | 0.0 | 0 |
|  | FRODOCK | 30.0 | 26.67 | 13.33 | **43.33** | **40.0** | 20.0 | **56.67** | **46.67** | 23.33 | 0 |
|  | HDOCK | **36.67** | **30.0** | **20.0** | 36.67 | 36.67 | **23.33** | 56.67 | 46.67 | **26.67** | 0 |
|  | PatchDock | 13.33 | 6.67 | 0.0 | 23.33 | 16.67 | 3.33 | 33.33 | 26.67 | 3.33 | 0 |
| Predicted<br>(127) | AlphaFold-Multimer | **57.48** | **51.97** | **26.77** | **59.84** | **54.33** | **32.28** | **59.84** | **54.33** | **32.28** | 0 |
|  | DiffDock-PP | * | * | * | * | * | * | 30.71 | 12.6 | 0.79 | 0 |
|  | FRODOCK | 16.54 | 13.39 | 5.51 | 24.41 | 18.9 | 6.3 | 31.5 | 24.41 | 7.87 | 0 |
|  | HDOCK | 16.54 | 12.6 | 4.72 | 18.9 | 14.17 | 4.72 | 26.77 | 17.32 | 4.72 | 0 |
|  | PatchDock | 7.87 | 6.3 | 1.57 | 12.6 | 11.02 | 2.36 | 16.54 | 13.39 | 2.36 | 0 |

**Table 8.**
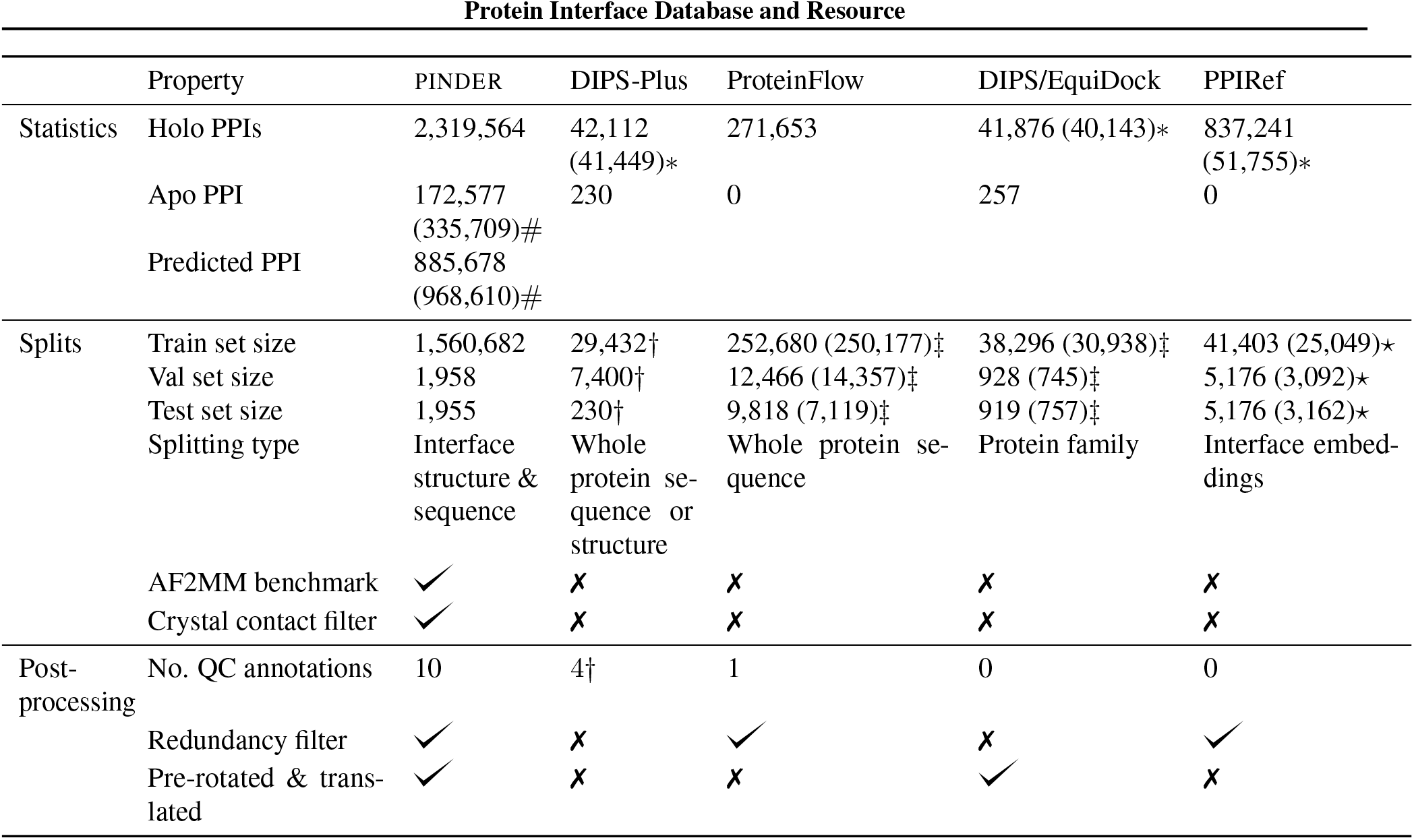
Properties of the different benchmark datasets reviewed or created in this work. # - The numbers PINDER systems with at least one chain mapped to an `apo` or `predicted` monomer. ∗-The numbers reported in the paper with the actual numbers of PPI in downloadable files in the brackets. Numbers for PPIRef500K (redundant) and PPIRef50K (non-redundant) datasets are provided. †-DIPS-Plus release 1.3.0 provides an option to use Foldseek for structural comparison. It uses DB5 dataset as the test set, where every PPI complex has a matching `apo` pair. DIPS-Plus provides 4 quality-filtered subsets and 9 residue-level feature annotations. ‡-PPIs matching PINDER IDs. ⋆ - PPIRef does not have a predefined split, we defined the splits as random samples with 80/20/20% ratios.

Both sequence and DIPS-EquiDock splits lead to models with over-estimated performance on their respective leaky test sets, while under-performing on structurally novel interfaces as shown by the strong dependence of performance on iAlign similarity to their respective training data (Figure 8). This is identical and even more pronounced to what has been observed for ALPHAFOLD 2-Multimer, as shown on the previous section. To isolate dataset size and diversity as the driving factors, the sampling process used for selecting diverse training data was identical for the sequence and PINDER splits, in fact resulting in *more* total training data and structural clusters in the sequence-split compared to the PINDER split, yet still yielding lower generalization (2). The increase in performance of the sequence-split with diversity sampling over the smaller, redundant DIPS-split, does indeed highlight the importance of diversity (Figure 8); however, its shortcoming compared to the PINDER splits on generalization indicates that the splits specifically can help models generalize (Figure 8). Excitingly, PINDER splits show no decrease in performance on structurally dissimilar interfaces despite not explicitly using iAlign to determine interface similarity clusters, indicating that the Foldseek-based protocol is robust. One possible explanation might be that over-fitting on train could encourage models to plateau after memorization of train/val leaked systems, rather than learning to generalize to novel interfaces by learning underlying physical or biological principles of interactions. Another factor could be better sampling via the interface-clusters during training. While further experiments are required to fully attribute these performance gains in the PINDER split to different reasons for why better clustering and splitting could help generalization, these experiments clearly demonstrate its positive impact on performance.

PINDER contains *apo* and predicted structures at 3 flexibility levels (described in section 2.2) with comparable number of examples from each flexibility category shown in Figure A.43. Leveraging this, we show in Figure 10, that predicted structures seem to have a better performance than *apo* structures, likely due to the fact that AlphaFold tends to predict structures closer to the *holo* conformation (Saldaño et al., 2022).

## 4. Conclusion

PINDER offers a substantial advancement in the field of Deep Learning-based protein-protein docking and complex modeling by addressing key limitations of existing training and benchmark datasets. PINDER’s emphasis on interface quality and diversity, achieved through strict quality filtering and clustering, ensures a more realistic evaluation of docking methods. The inclusion of *apo* and predicted structures, both in training and test, with different levels of difficulties is unique to PINDER, and enables researchers to push the boundaries of generalizable method development which can be applied to real-world scenarios. Additionally, diverse sampling through interface clustering contributes to both a balanced training scheme, and creation of a large and diverse test set. The boost in dataset size through our reproducible and automated curation method is expected to significantly improve the generalization capabilities of Deep Learning models trained on PINDER. Furthermore, the interface-based de-leaking method coupled with quality control based on ECOD classification, ensures robustness of the dataset splits. These methodological advancements collectively contribute to a more reliable and rigorous evaluation of protein-protein complex modeling, ultimately accelerating progress in this critical area of research.

While PINDER makes significant strides, several limitations highlight areas for future improvement. Most evidently, PINDER is currently focusing on biological dimers both to increase quality and since most methods currently work on dimer-based docking. As more and more methods will expand beyond these limits, such as via co-folding approaches, PINDER will be generalized to higher-order oligomers. Additionally, there are a few smaller methodological limitations - for instance, the reliance on single reference conformations and the inherent bias towards homodimers in the dataset can impact the accuracy and generalizability of the models. Finally, improvements in *apo* pairing and the integration of more advanced tools, such as iAlign, into the alignment methodology could enhance the dataset’s precision. Addressing these limitations could lead to even larger datasets, better performance and evaluation in future iterations of PINDER. We provide a more detailed discussion of the limitations of the PINDER dataset and methodology in the Appendix A.1.

