## Appendix for "PINDER: The protein interaction dataset and evaluation resource"

### A. Appendix

#### A.1. Detailed limitations

**Alignment methodology** Our primary focus was to minimize leakage within the test/validation splits. To achieve this, we employed low score thresholds in our all-vs-all score computations, specifically using Foldseek and MMseqs2. While alignments between monomers were required to have a minimum coverage of the interface for each monomer, the alignments themselves were not specific to the interfaces. Additionally, we masked training data with indirect (transitive) hits at a depth of two to enhance recall (see algorithm1 and methods2). Although this approach yielded a diverse set of clusters, it has limitations. Dissimilar interfaces may have been considered similar and consequently removed from the training data, and individual clusters can have high range of interfaces. Future iterations could address this by incorporating tools with greater precision, such as iAlign, into the splitting algorithm.

**PRODIGY-cryst** For the PINDER test, we selected protein complexes using multiple criteria to ensure quality, though each choice involved trade-offs. The threshold of 0.5 in the PRODIGY algorithm signifies a relatively low-quality interaction prediction, with scores between 0.5 and 0.6 falling into this category. Moreover, scores below 0.5 might still represent biologically relevant interfaces, potentially affecting the accuracy of interaction assessments.

**Dimers** PINDER exclusively utilizes interactions at dimeric scale for training and evaluation. Even though we utilize biological dimers in the test and validation sets, in training split they were often extracted from higher-order oligomeric complexes. This could limit their biological plausibility in isolation, for example, the binding of two monomers can create a neo-substrate for a third interacting chain that would otherwise not have the requisite binding affinity. Additionally, the classification of protein complexes into oligomeric states, such as homodimers and heterodimers, relies heavily on specific definitions and contextual interpretations. This variability can introduce inconsistencies in the analysis and interpretation of interaction data.

**Single reference conformation** Evaluating protein interactions using a single reference conformation restricts analysis flexibility. Proteins can adopt multiple conformations, and relying on one reference may not capture the full range of interactions, reducing accuracy. This is particularly important to note given the inclusion of *apo* and predicted monomers as augmentation. Incorporating multiple conformational states can provide a more accurate and comprehensive assessment of protein-protein interactions.

**AlphaFold2 monomers** While the PINDER dataset includes many predicted AlphaFold2 monomers, it is limited to only those present in AFDB 4. For structures with undefined UniProt IDs, we could have inferred sequences to broaden the dataset.

**Homodimer bias** The PINDER dataset exhibits a heavy bias towards homodimers, which is reflective of this bias as found in the PDB source data. This potentially limits the generalizability of the models trained on it.

***Apo* pairing** Pairing *apo* forms could be improved by integrating Foldseek and MMseqs2 on monomers, as the current approach is primarily UniProt-based.

#### A.2. Diversity and leakage

To assess the quality of splits generated by our approach (See Algorithm 1), we compute several metrics of split diversity and split leakage for the PINDER-XL and PINDER-Val sets (Table 3 and Table 4). We additionally include these metrics for the training and validation sets of three existing datasets: DIPS-EquiDock, ProteinFlow and PPIRef. In all cases, we restrict our analysis to consider only systems for which both of the binding partners are longer than 40 residues. Additionally, for the existing datasets we restrict our analysis to consider only systems for which both binding partners are unambiguously contained within a PINDER dimer. The mapping procedure from existing dataset to PINDER dimers is described in A.4.3.

To ensure that test and validation splits cover a sufficiently broad subspace of protein-protein interactions we compute:

- **Cluster diversity**: The ratio of the number of unique PINDER clusters in the split of interest to the total number of unique PINDER clusters in the union of the Test, Val, and Train sets.

- **ECOD single-chain diversity**: The ratio of the number of unique ECOD (Cheng et al., 2014) domain labels in the split of interest to the total number of unique ECOD domain labels in the union of the Test, Val, and Train sets. Note that each PINDER system will have two ECOD domain labels, corresponding to the receptor and ligand chains.

- **ECOD chain-pair diversity**: The ratio of the number of unique ECOD (Cheng et al., 2014) domain *pair labels* in the split of interest to the total number of unique ECOD domain *pair labels* in the union of the Test, Val, and Train sets. Note that each PINDER system will have one ECOD domain pair label.

- **Pfam clan diversity**: The ratio of the number of unique Pfam (Mistry et al., 2020) clans in the split of interest to the total number of unique Pfam clans in the union of the Test, Val, and Train sets. Note that each PINDER system will have two Pfam clan labels.

- **Unique ECOD chain pairs**: The number of systems in the split of interest that have an ECOD pair label that does not appear in the Train set, expressed as a ratio with the total size of the split of interest.

To ensure that test and validation splits are sufficiently distinct from the training set, we compute:

- **UniProt pair leakage**: The number of systems in the split of interest that have a UniProt pair label that also appears in the PINDER-Train set, expressed as a ratio with the total size of the split of interest. Note that we exclude systems that contain a UniProt label of 'UNDEFINED' from this analysis. Note that UniProt pair leakage can result in false positives as the sequence the sequences may not necessarily overlap, in case of partially-resolved proteins, for instance.

- **ECOD pair leakage**: The number of systems in the split of interest that have an ECOD pair label that also appears in the PINDER-Train set, expressed as a ratio with the total size of the split of interest.

- **iAlign interface pair leakage**: The number of systems in the split of interest that have at least one similar interface meeting the iAlign criteria that appears in the PINDER-Train set, expressed as a ratio with the total size of the split of interest.

**Protein family and domain diversity** In addition, we delved into the of protein family and domain diversity, focusing on chains from cluster representatives across PINDER's train, validation, and test datasets. We first filtered any peptide chains out, specifically those less than 40 amino acids in length, and subsequently matched interface region to Pfam (Protein families database) domain-level classification that correspond to The F-group level (family level) in ECOD (Liao et al., 2018) as well as globally to PANTHER (Protein ANalysis THrough Evolutionary Relationships) global family classifications (Thomas et al., 2022). Our findings revealed a notable degree of diversity within the test and validation data, with the median count for each family being 2, and matching to our test sampling scheme of two representatives per cluster. To provide a comprehensive overview, we have included an appendix, showcasing the top 50 most abundant families and ECOD domains.

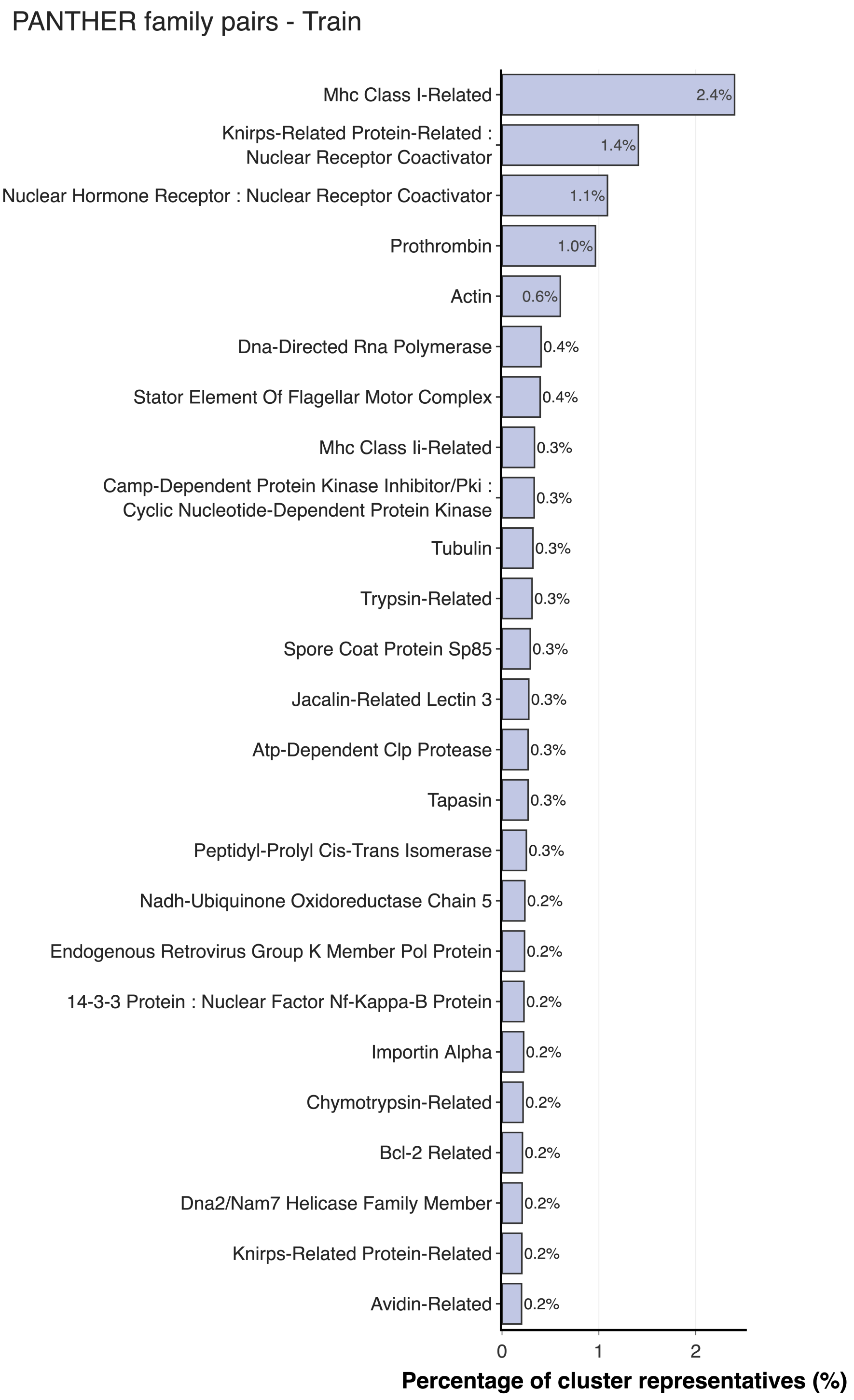


*Figure A.1.* **PANTHER Classes - PINDER-Train**. Top 50 panther classes and paired panther classes for interacting chains, ranked and represented by the ratio present in different splits.

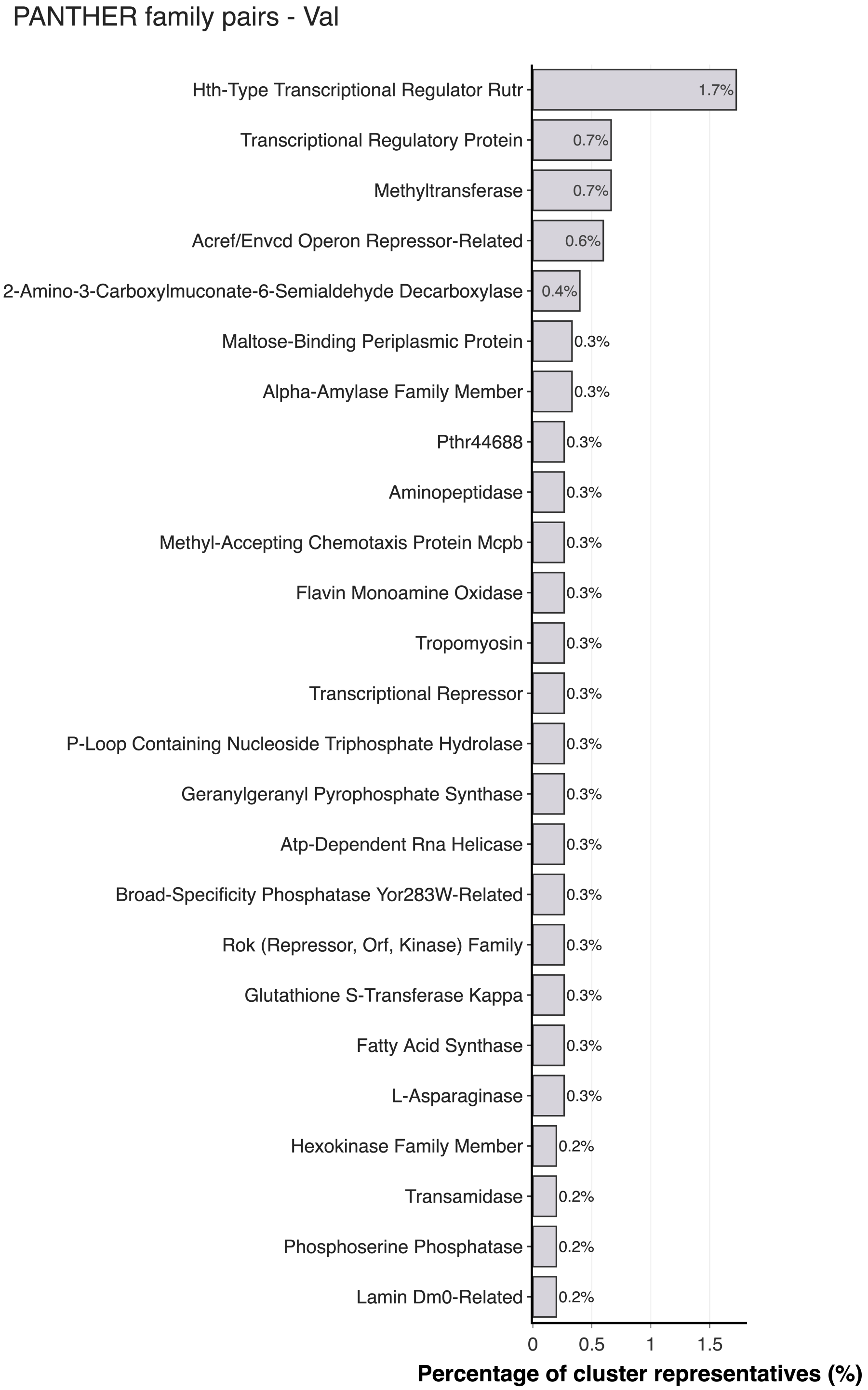


*Figure A.2.* **PANTHER Classes - PINDER-Val**. Top 50 panther classes and paired panther classes for interacting chains, ranked and represented by the ratio present in different splits.

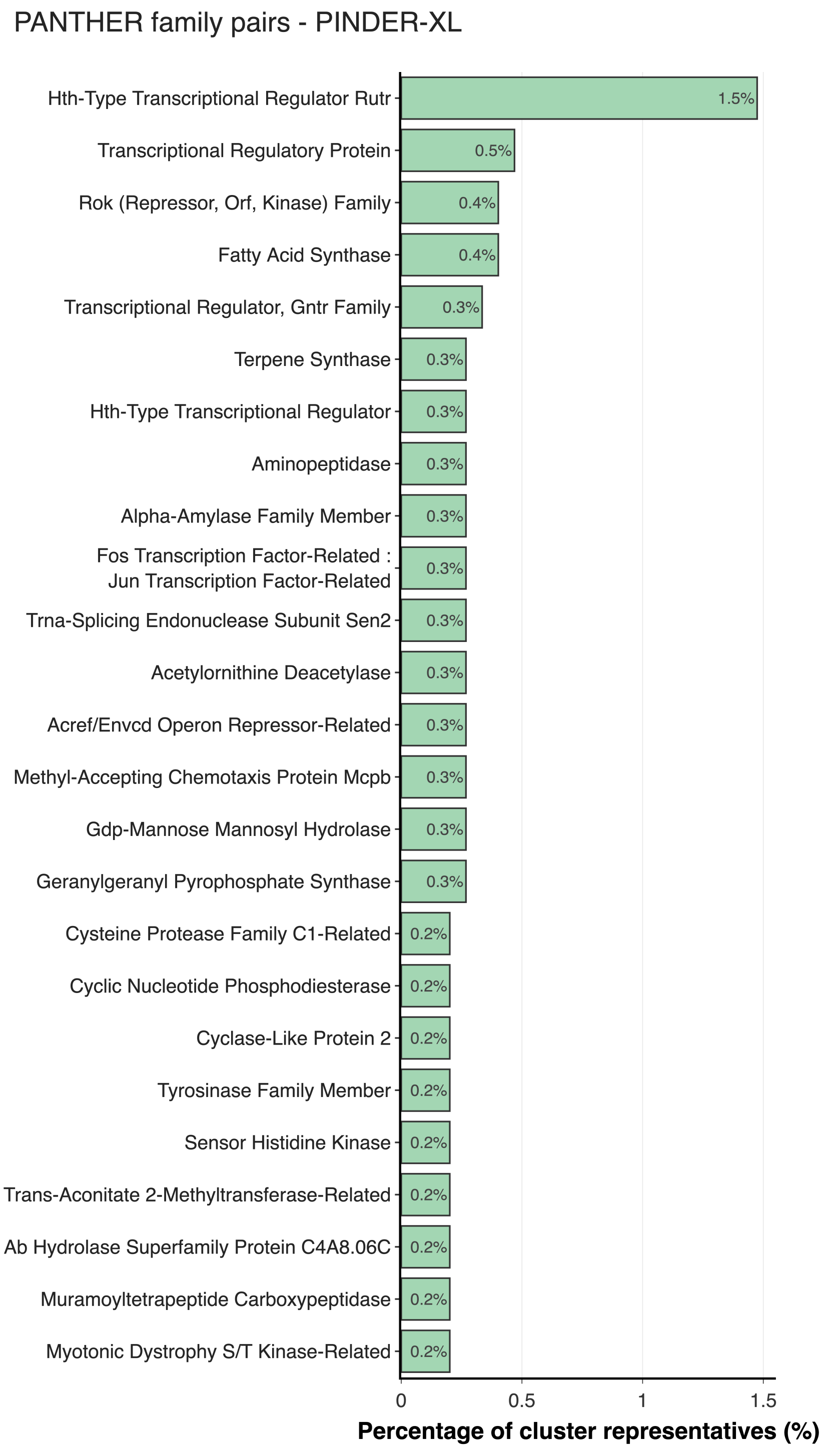


*Figure A.3.* **PANTHER Classes - PINDER-XL**. Top 50 panther classes and paired panther classes for interacting chains, ranked and represented by the ratio present in different splits.

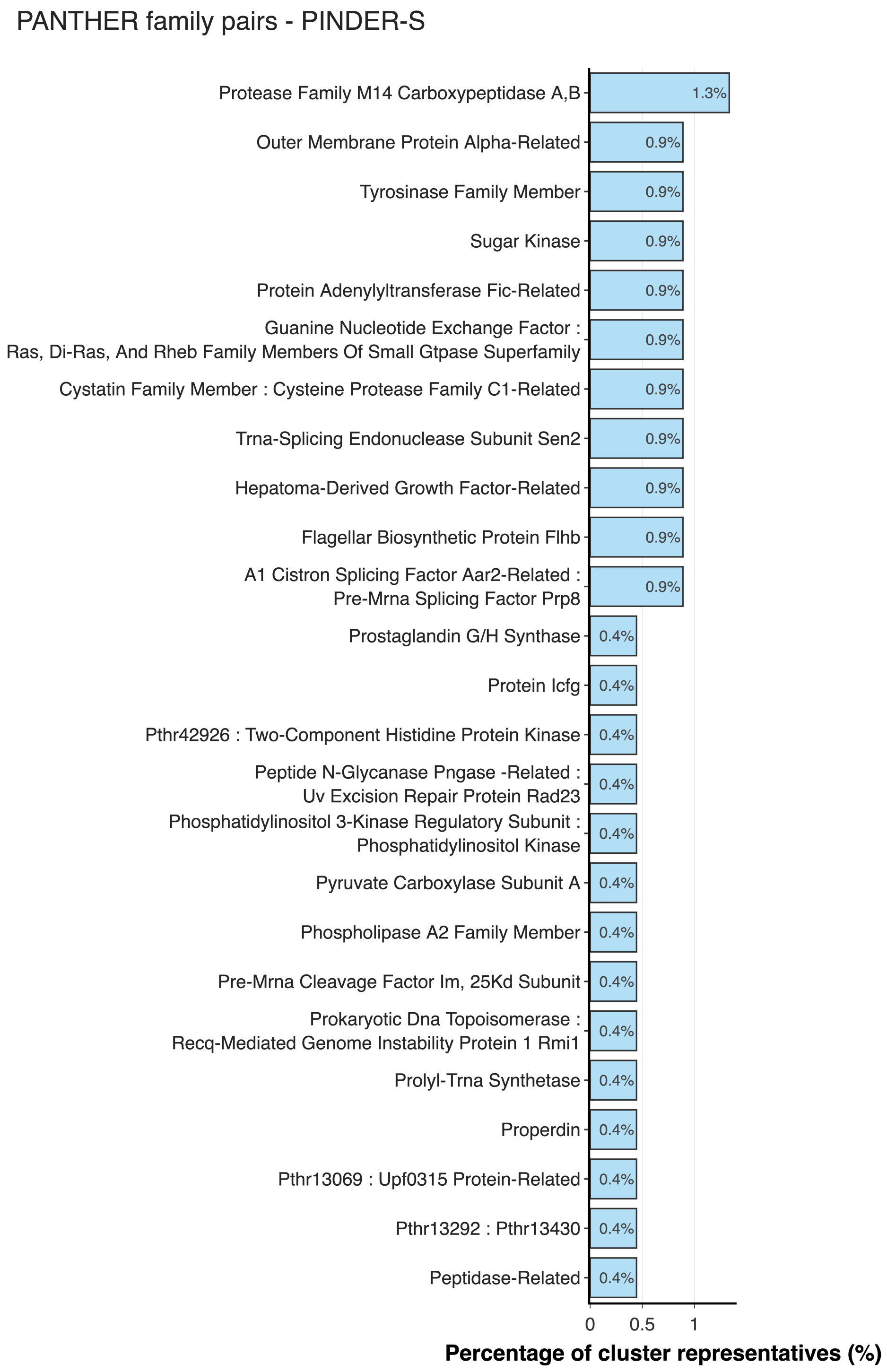


*Figure A.4.* **PANTHER Classes - PINDER-S**. Top 50 panther classes and paired panther classes for interacting chains, ranked and represented by the ratio present in different splits.

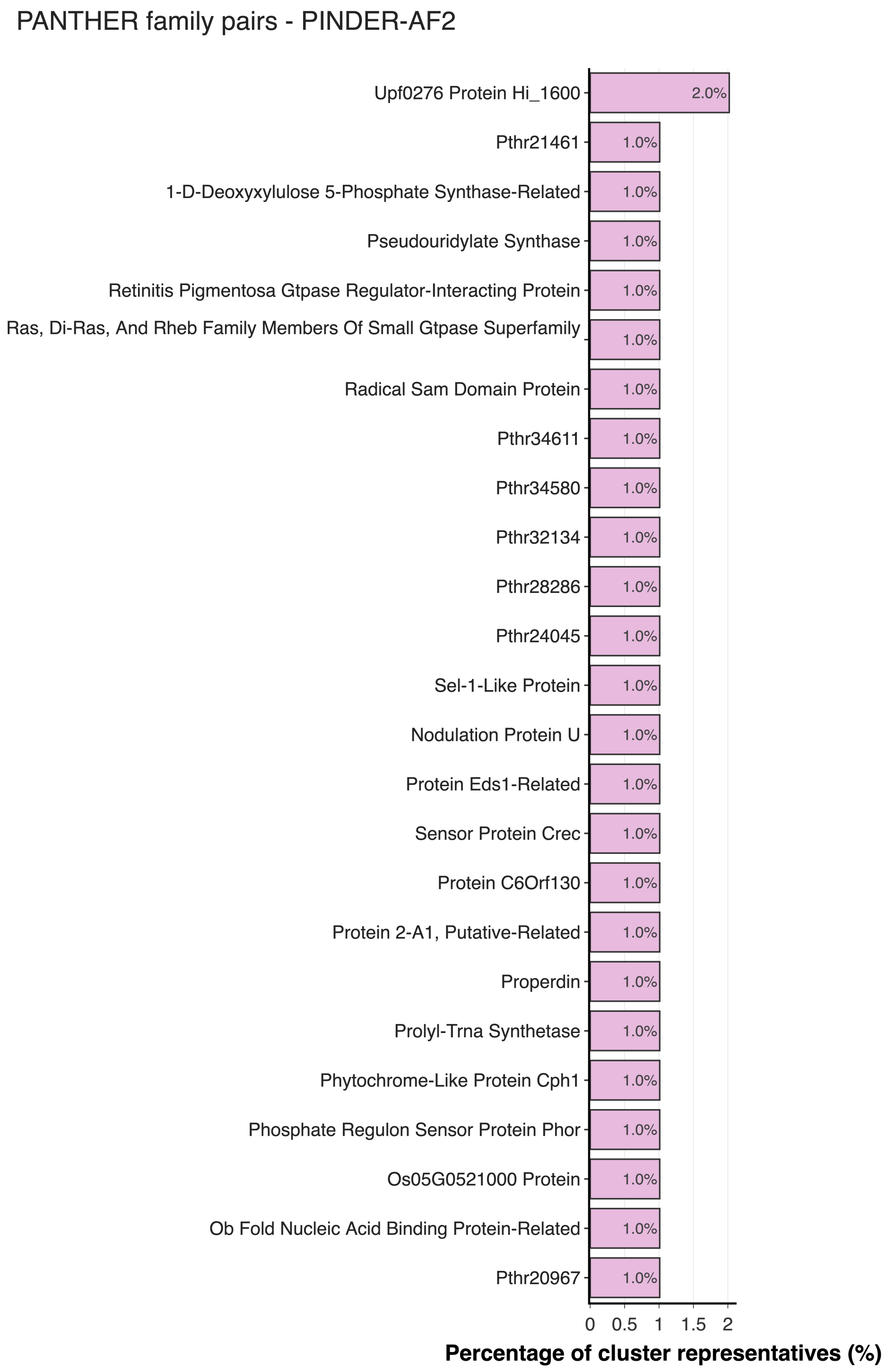


*Figure A.5.* **PANTHER Classes - PINDER-AF2**. Top 50 panther classes and paired panther classes for interacting chains, ranked and represented by the ratio present in different splits.

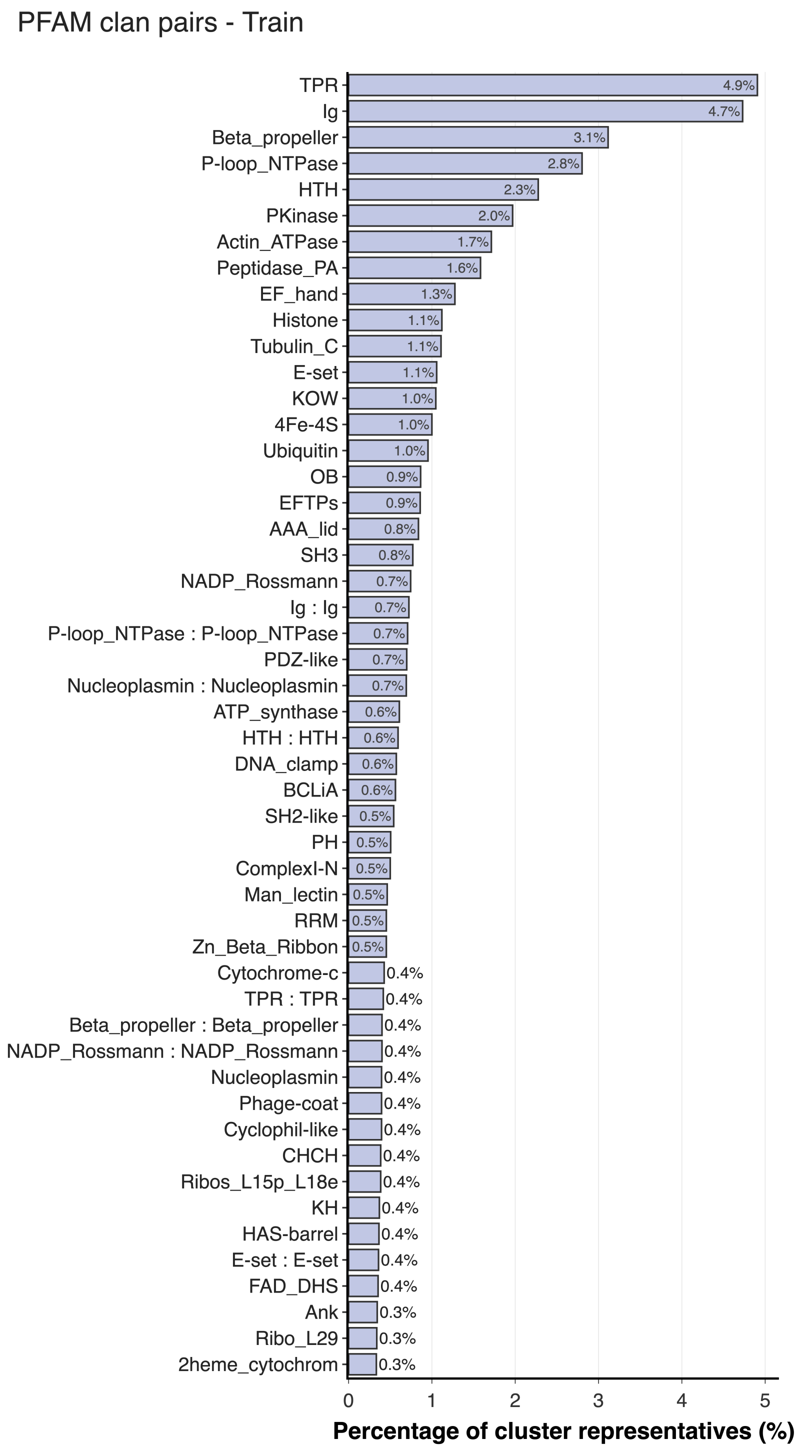


*Figure A.6.* **PFAM Classes - PINDER-Train**. Top 50 Pfam classes and paired Pfam classes for interacting chains, ranked and represented by the ratio present in different splits.

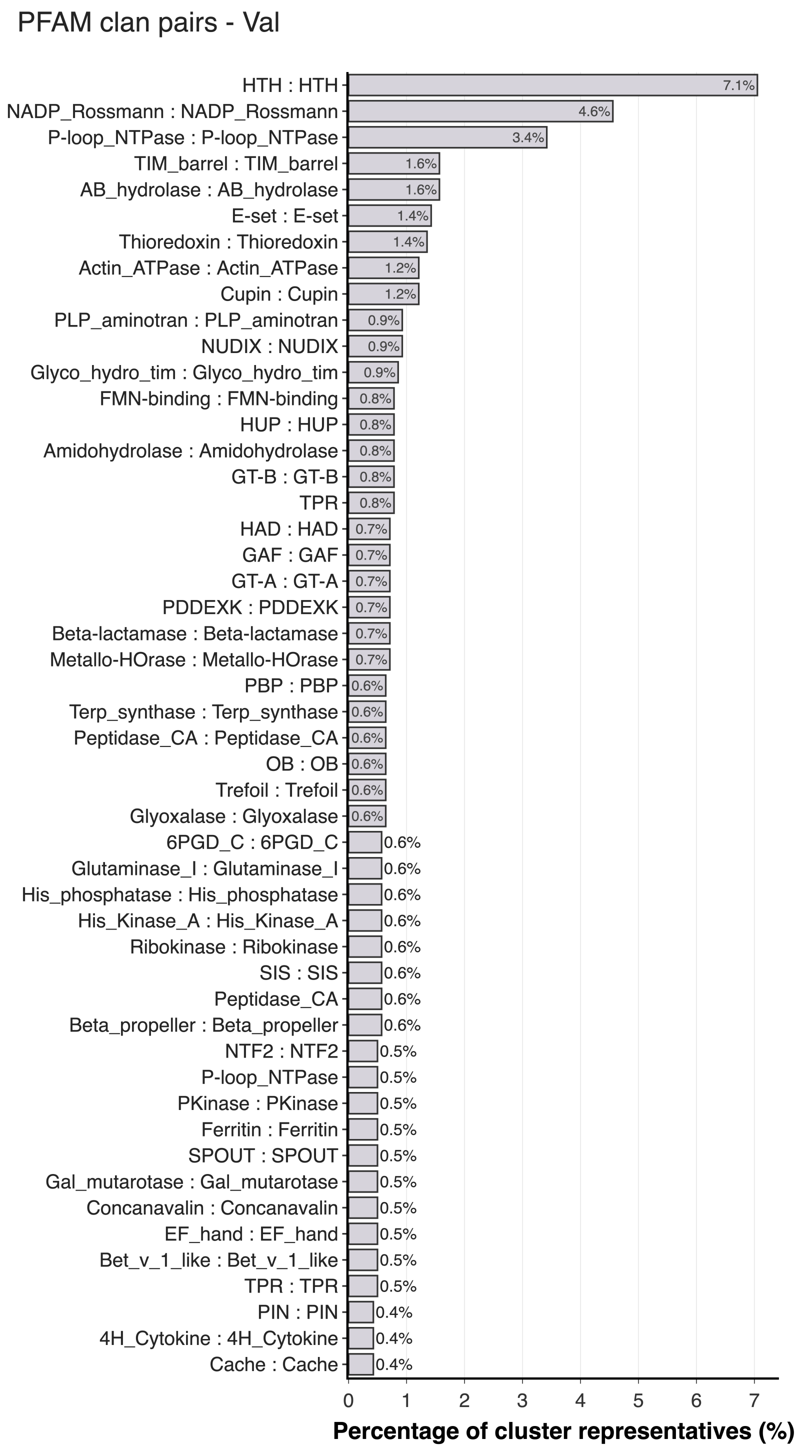


*Figure A.7.* **PFAM Classes - PINDER-Val**. Top 50 Pfam classes and paired Pfam classes for interacting chains, ranked and represented by the ratio present in different splits.

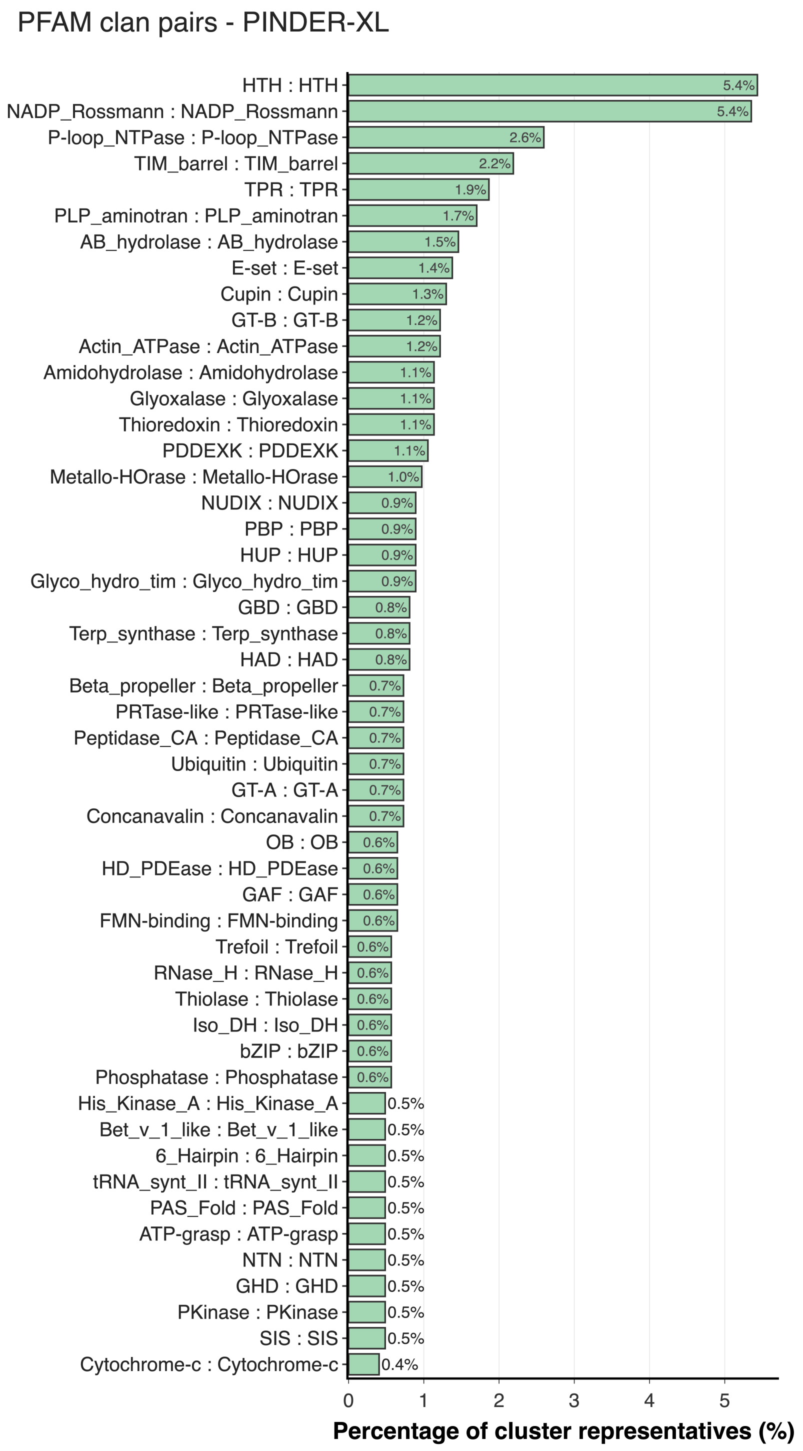


*Figure A.8.* **PFAM Classes - PINDER-XL**. Top 50 Pfam classes and paired Pfam classes for interacting chains, ranked and represented by the ratio present in different splits.

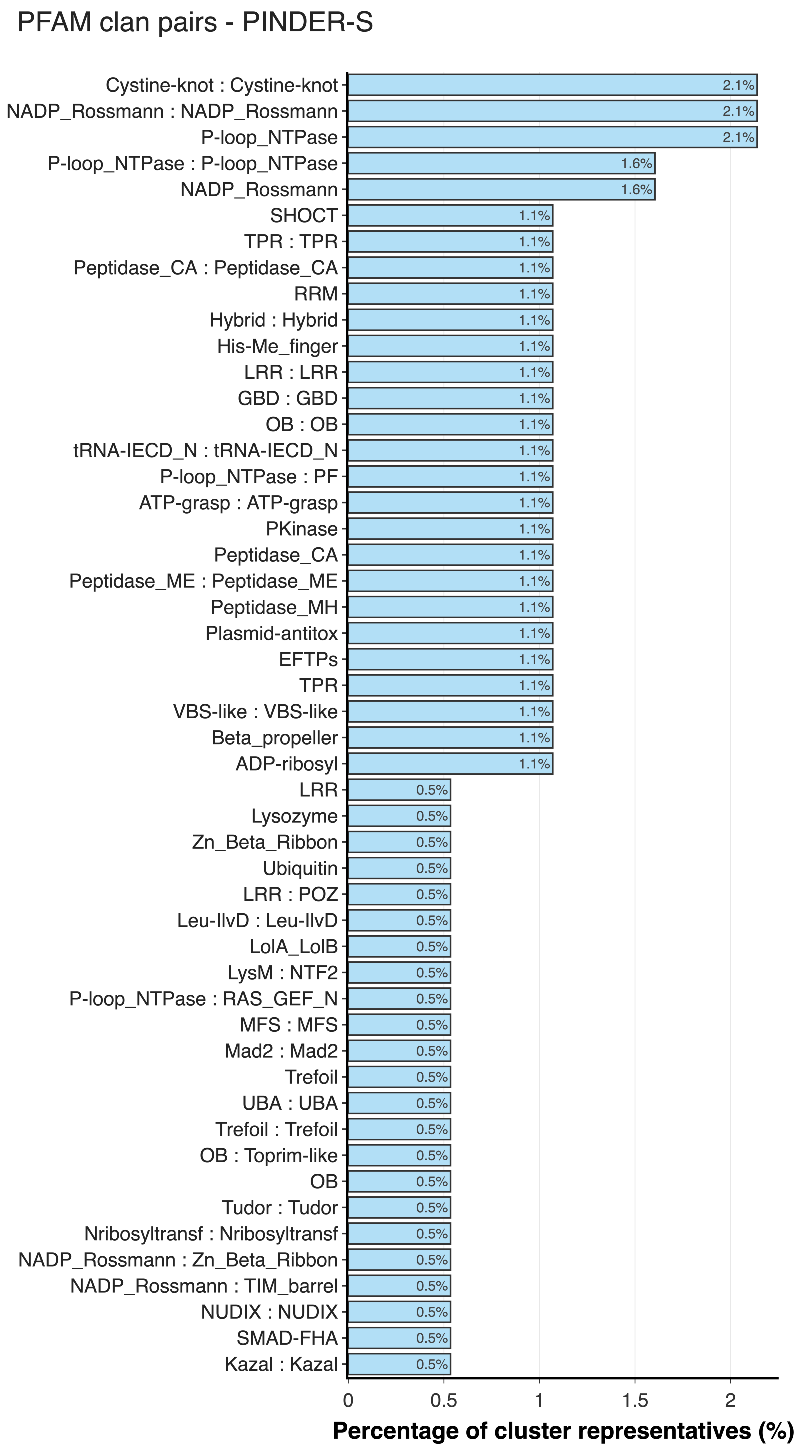


*Figure A.9.* **PFAM Classes - PINDER-S**. Top 50 Pfam classes and paired Pfam classes for interacting chains, ranked and represented by the ratio present in different splits.

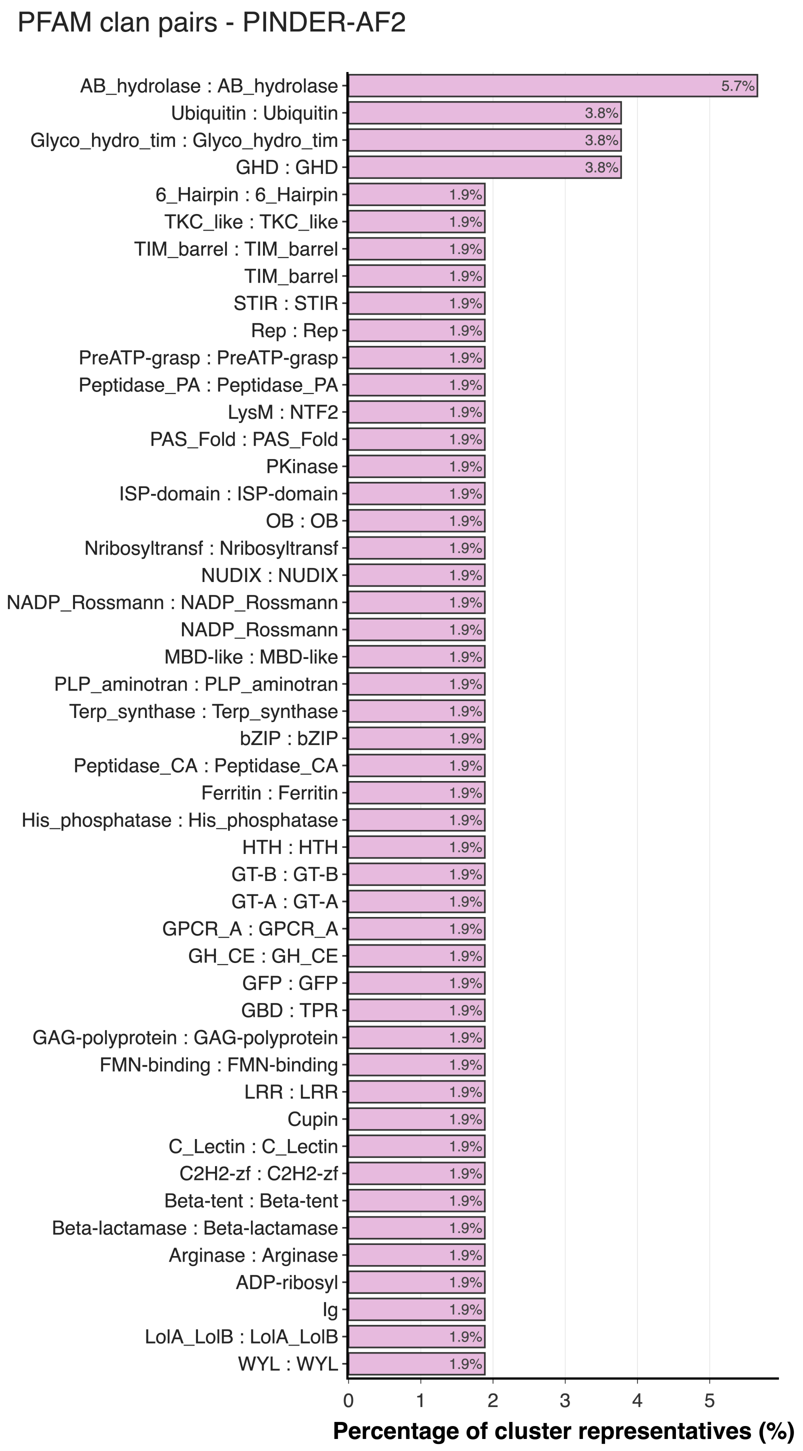


*Figure A.10.* **PFAM Classes - PINDER-AF2**. Top 50 Pfam classes and paired Pfam classes for interacting chains, ranked and represented by the ratio present in different splits.

#### A.3. Additional figures

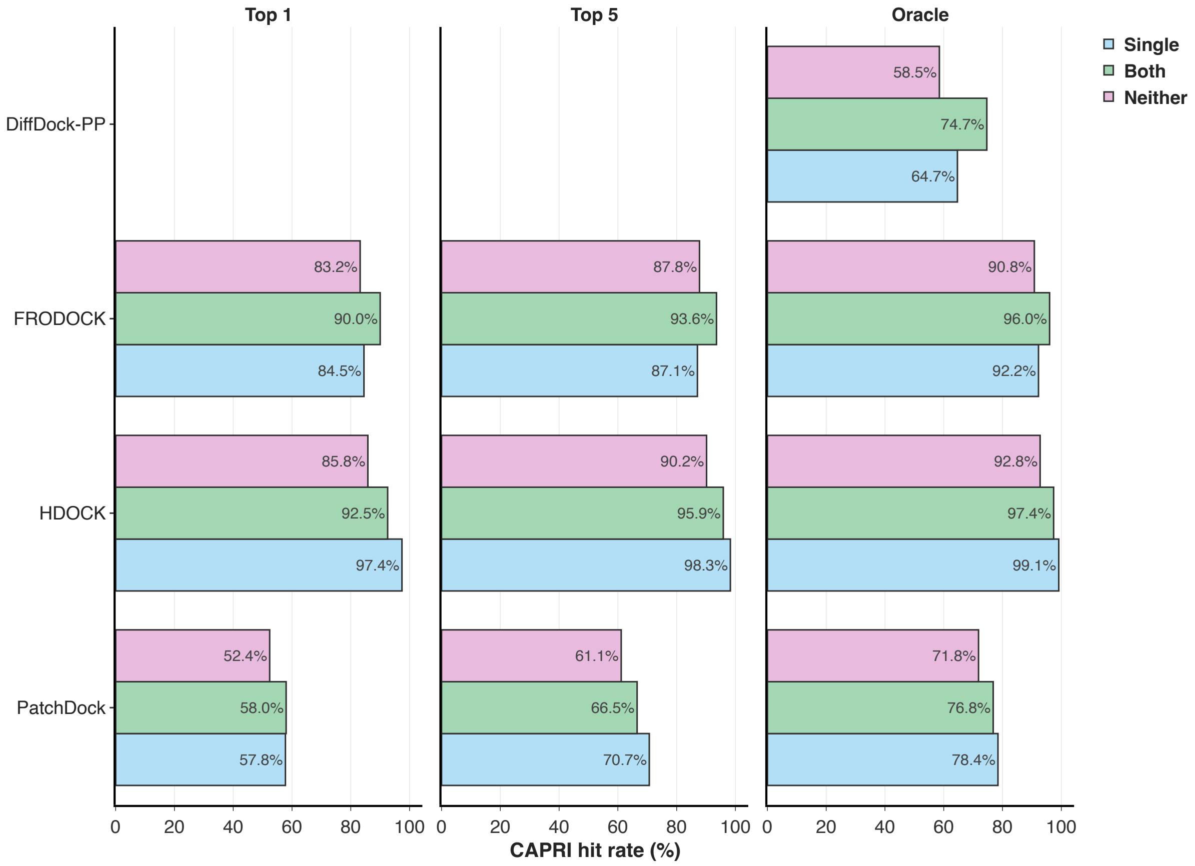


*Figure A.11.* Bar-chart showing performance difference between different categories of chain uniqueness in PINDER-XL set. ”Both” includes systems unique to the test set, ”Single” contains systems with a single chain unique to the test set, and ”Neither” contains systems with neither of the chains unique to the test set.

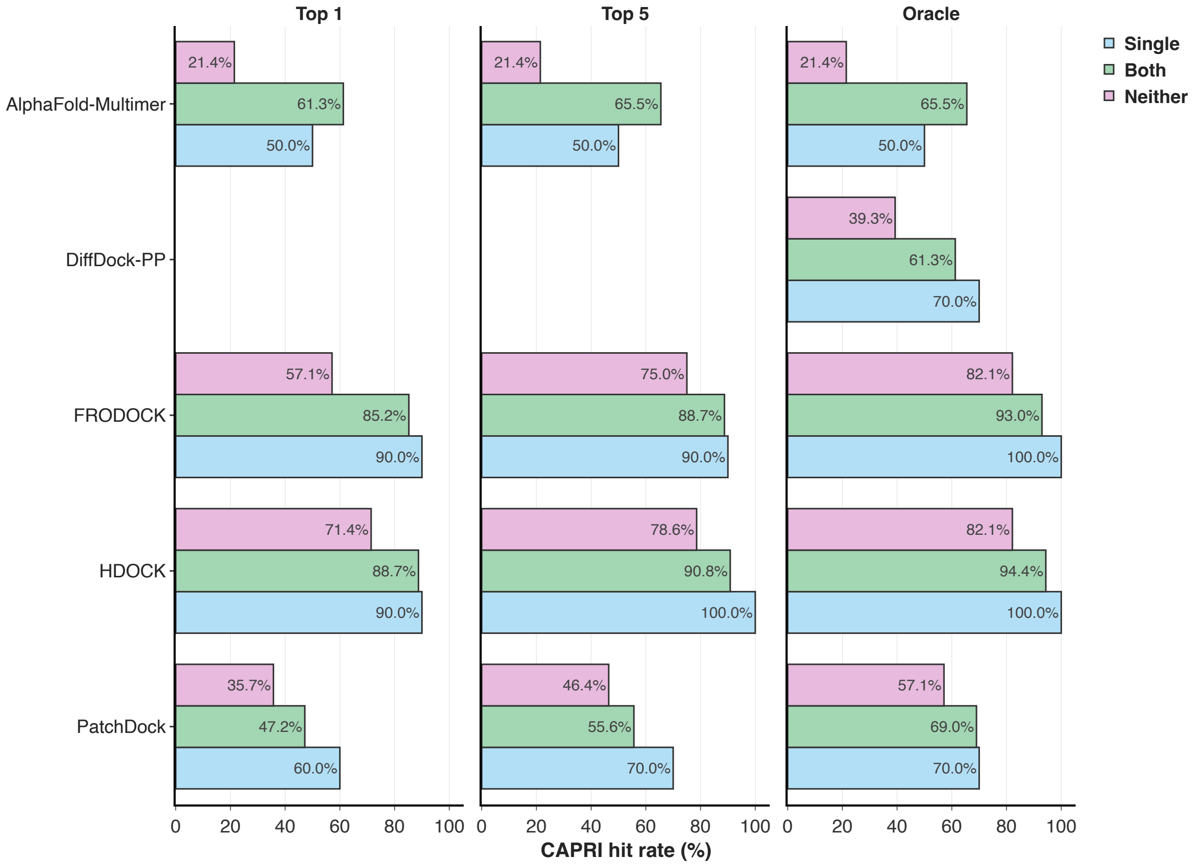


*Figure A.12.* Bar-chart showing performance difference between different categories of chain uniqueness in PINDER-AF2 set. ”Both” includes systems unique to the test set, ”Single” contains systems with a single chain unique to the test set, and ”Neither” contains systems with neither of the chains unique to the test set.

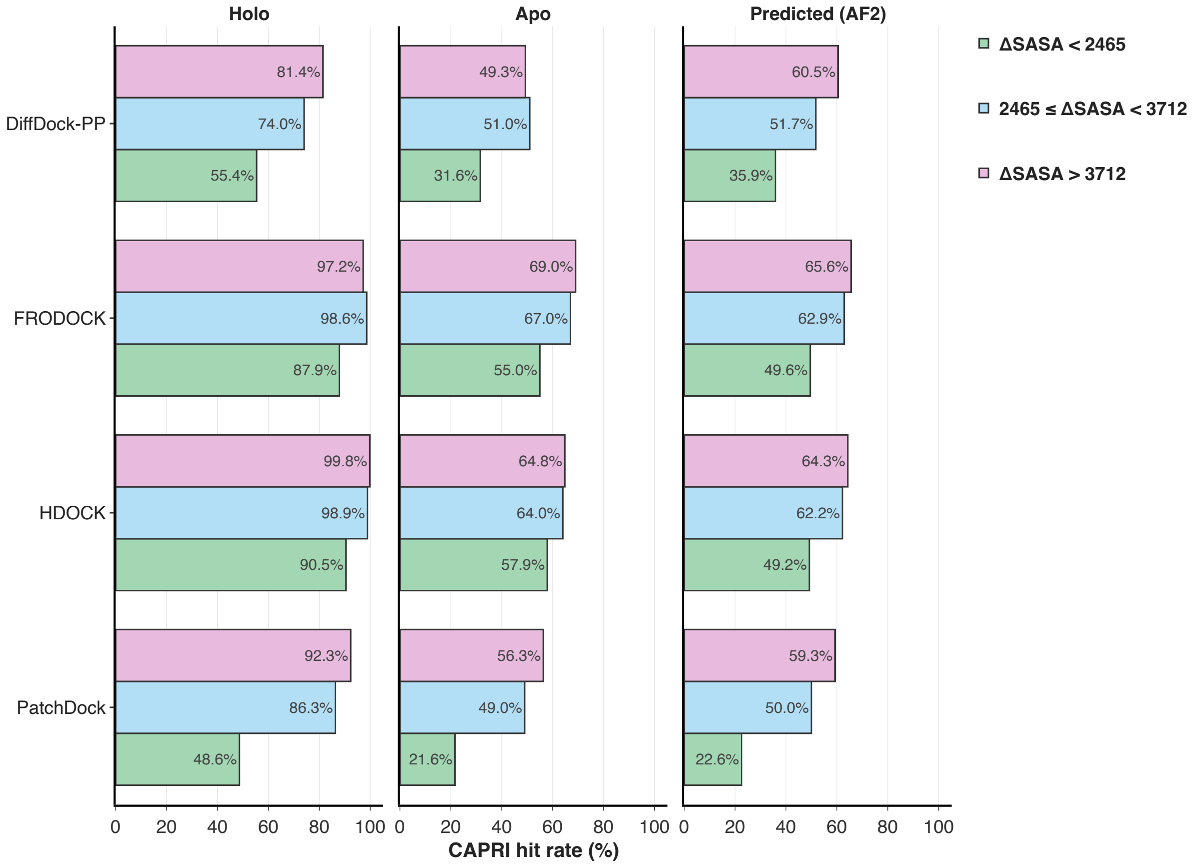


*Figure A.13.* Oracle results of three different buried accessible surface area (dSASA) ranges in PINDER-XL set.

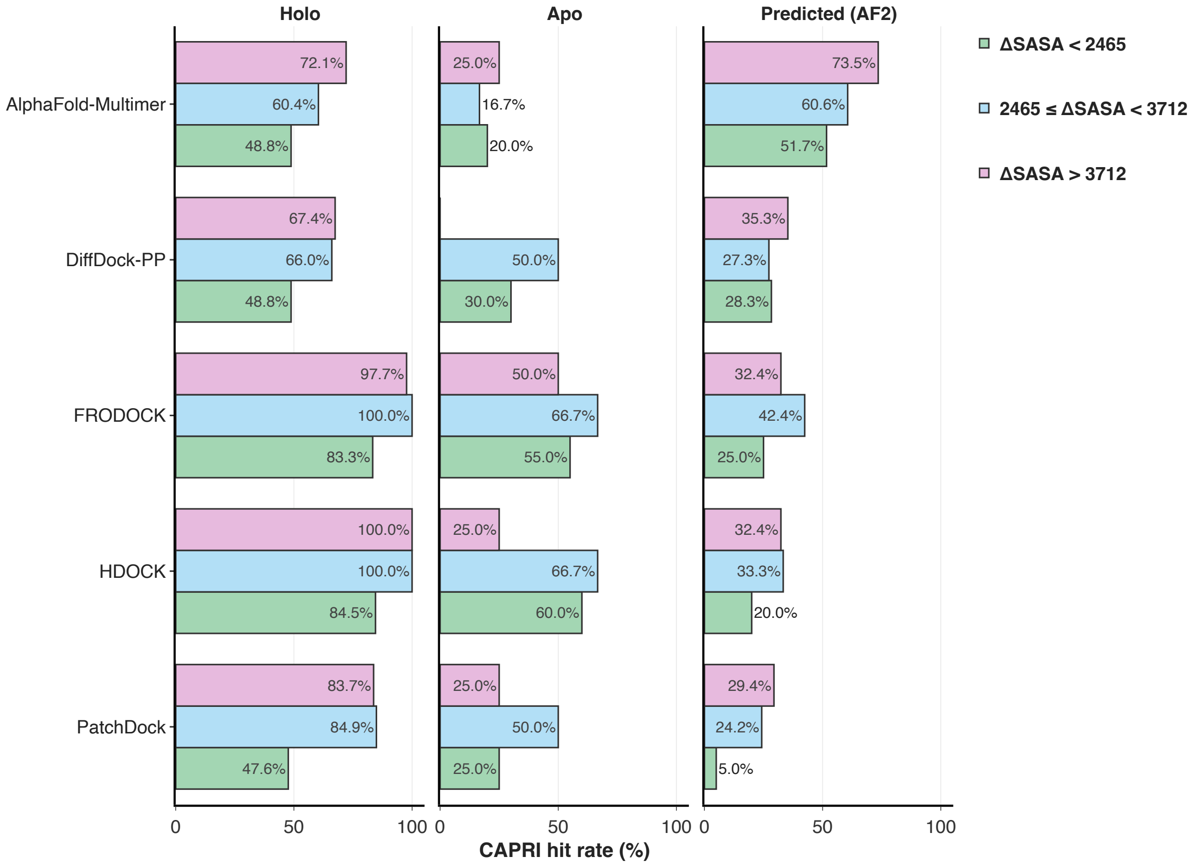


*Figure A.14.* Oracle results of three different buried accessible surface area (dSASA) ranges in PINDER-AF2 set.

##### A.3.1. POSE VALIDITY

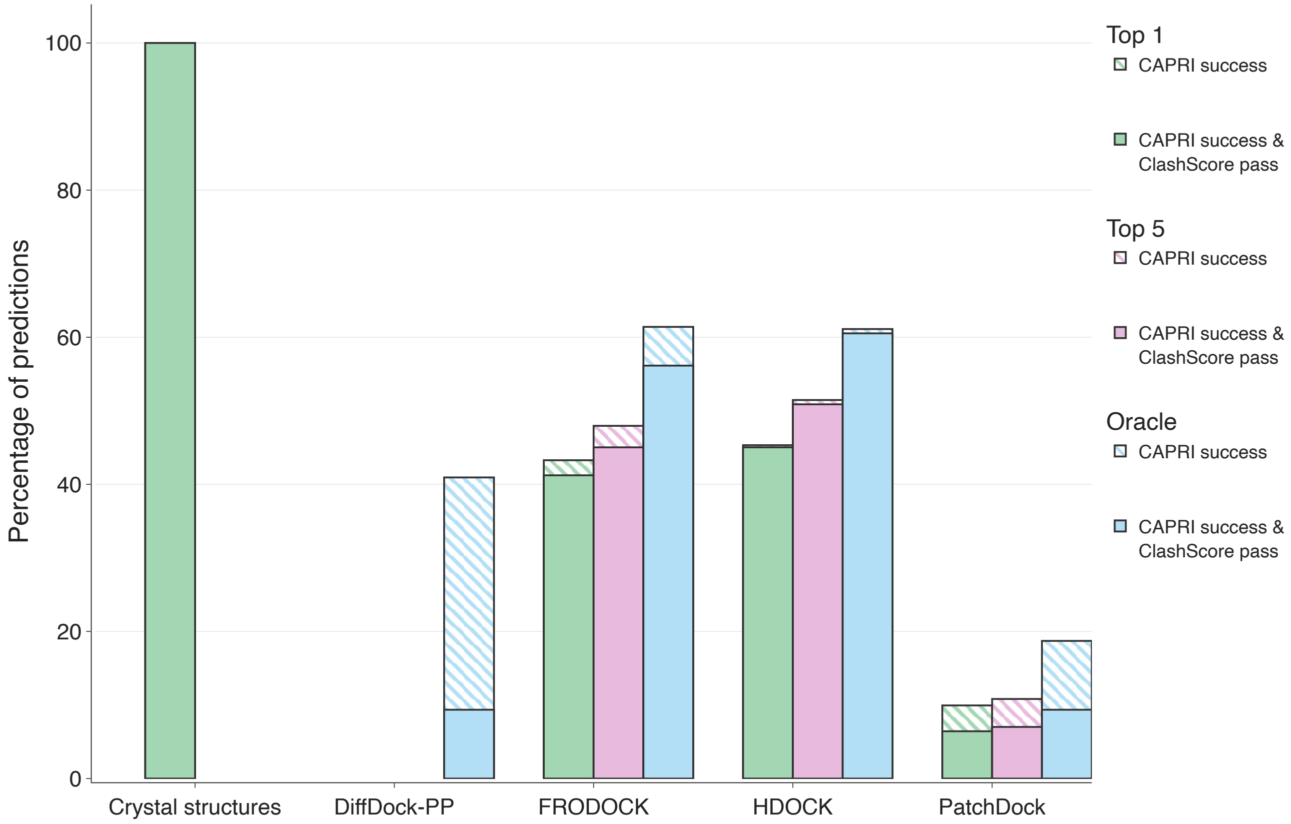


*Figure A.15.* CAPRI success rates of benchmarked methods on *apo* systems across PINDER-XL, with and without low ($< 0.1$) VoroMQA clash scores for oracle, top-1 and best of top-5 poses.

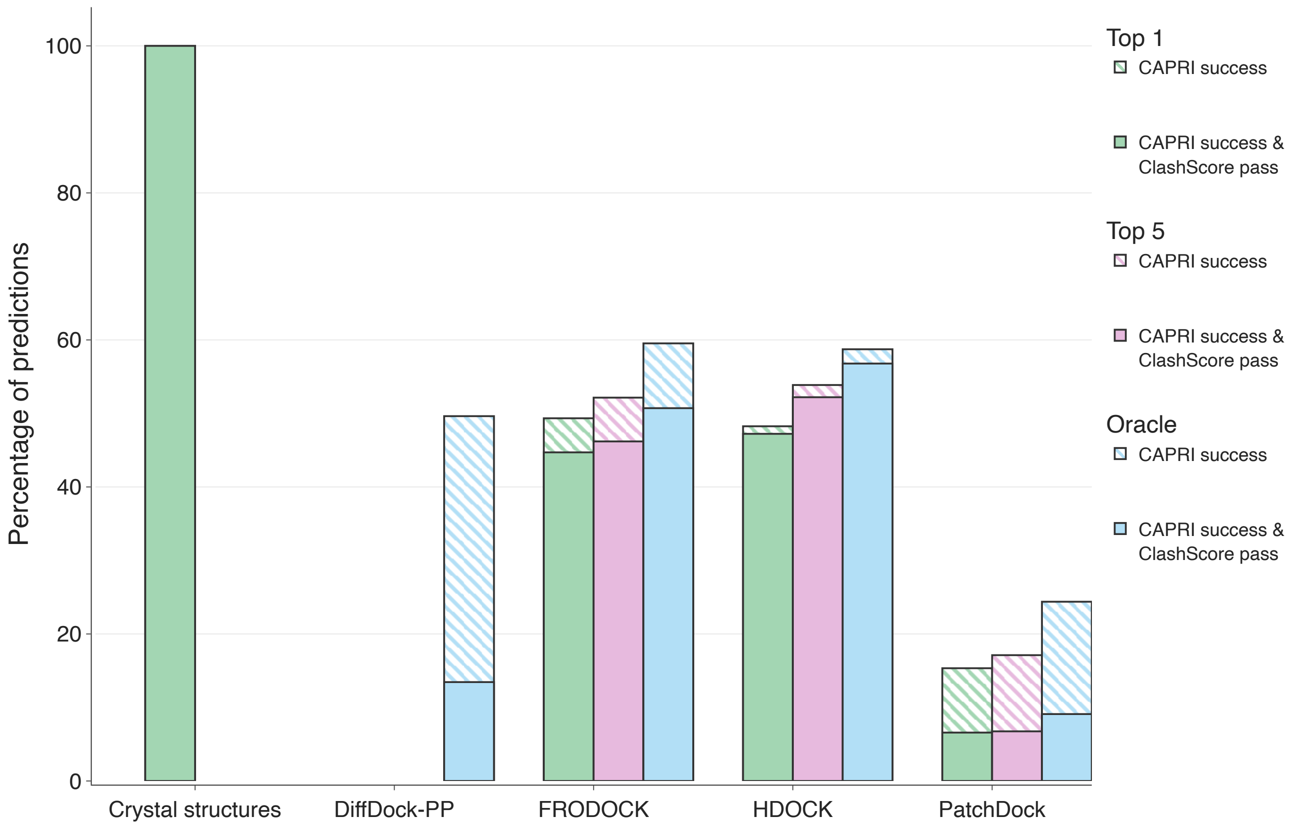


*Figure A.16.* CAPRI success rates of benchmarked methods on predicted systems across PINDER-XL, with and without low ($< 0.1$) VoroMQA clash scores for oracle, top-1 and best of top-5 poses.

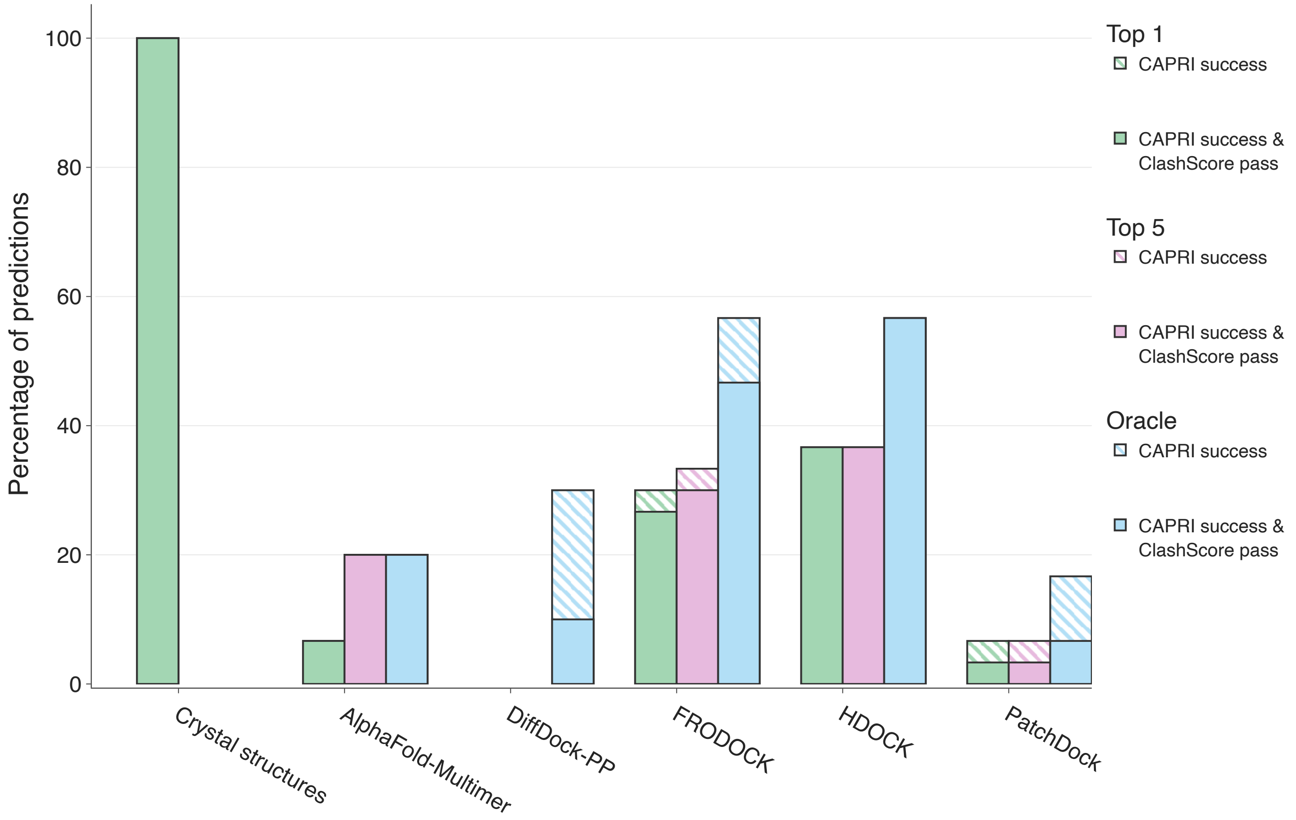


*Figure A.17.* CAPRI success rates of benchmarked methods on *apo* systems across PINDER-AF2, with and without low ($< 0.1$) VoroMQA clash scores for oracle, top-1 and best of top-5 poses.

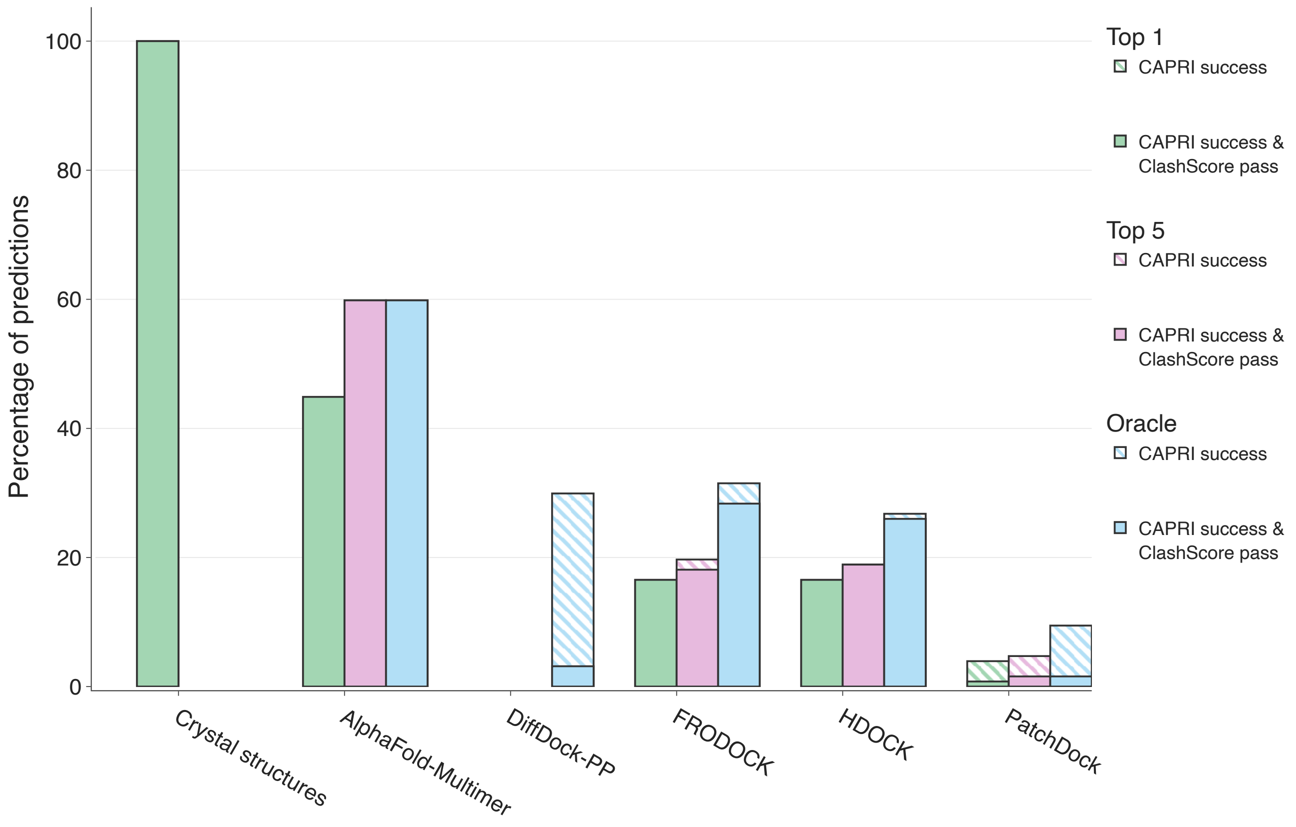


*Figure A.18.* CAPRI success rates of benchmarked methods on predicted systems across PINDER-AF2, with and without low ($< 0.1$) VoroMQA clash scores for oracle, top-1 and best of top-5 poses.

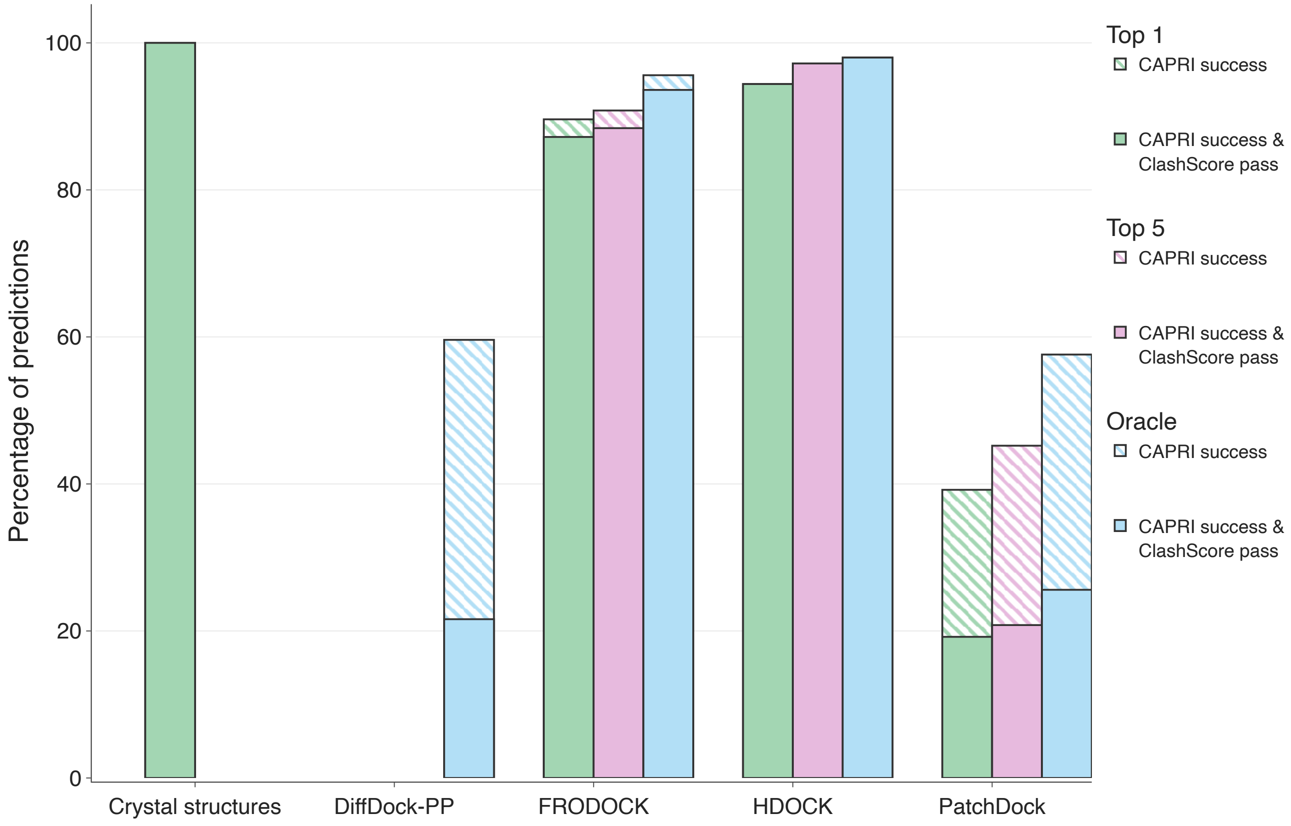


*Figure A.19.* CAPRI success rates of benchmarked methods on *holo* systems across PINDER-S, with and without low ($< 0.1$) VoroMQA clash scores for oracle, top-1 and best of top-5 poses.

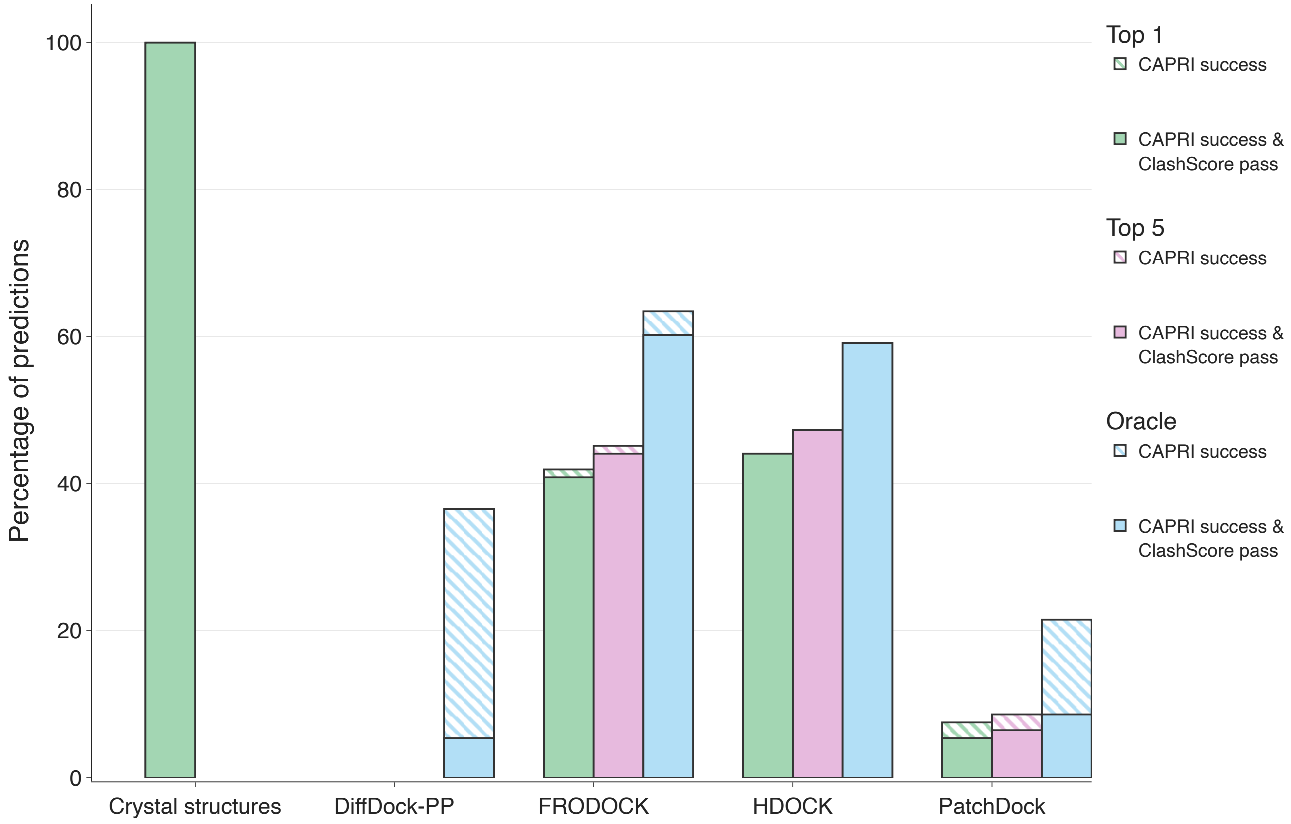


*Figure A.20.* CAPRI success rates of benchmarked methods on *apo* systems across PINDER-S, with and without low ($< 0.1$) VoroMQA clash scores for oracle, top-1 and best of top-5 poses.

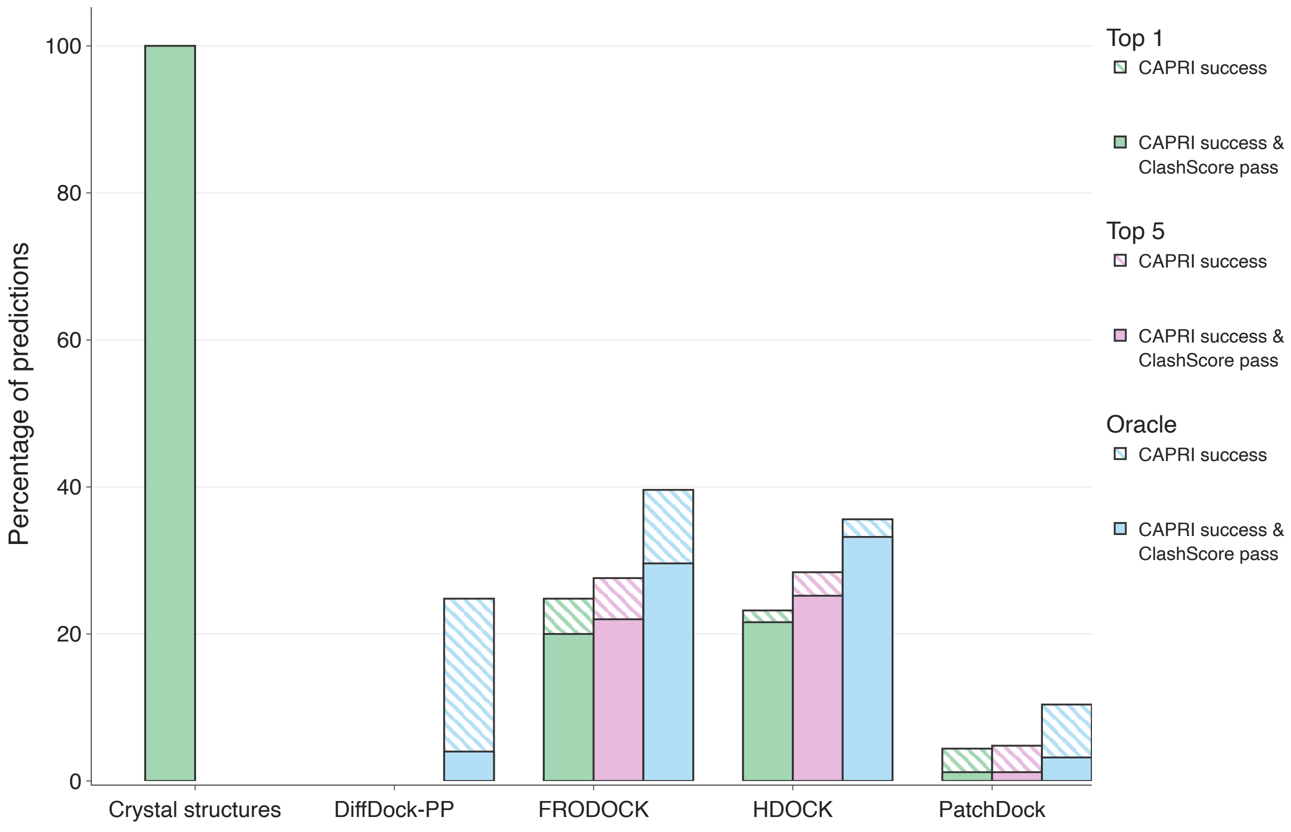


*Figure A.21.* CAPRI success rates of benchmarked methods on predicted systems across PINDER-S, with and without low ($< 0.1$) VoroMQA clash scores for oracle, top-1 and best of top-5 poses.

##### A.3.2. Distribution of clash metrics relative to crystal structures

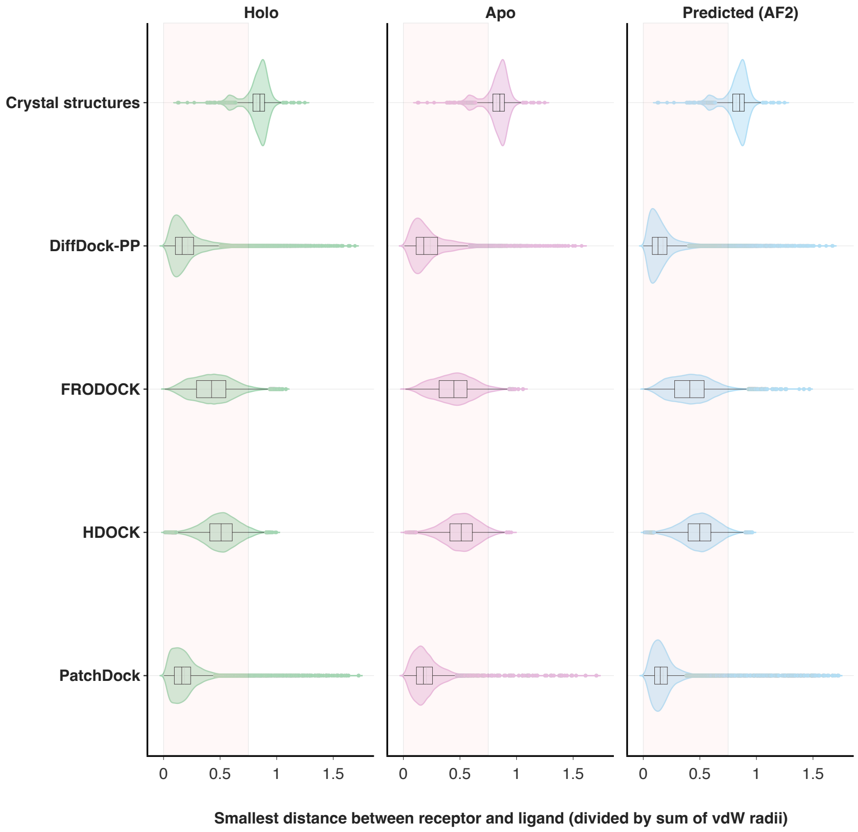


*Figure A.22.* Minimum distances between protein chains across PINDER-XL. Distance is the smallest pairwise distance of heavy atoms of the receptor and ligand protein chains normalized by their sum of van der Waals radii. The red area highlights the rejection zone below the cutoff of 0.75 (Buttenschoen et al., 2024).

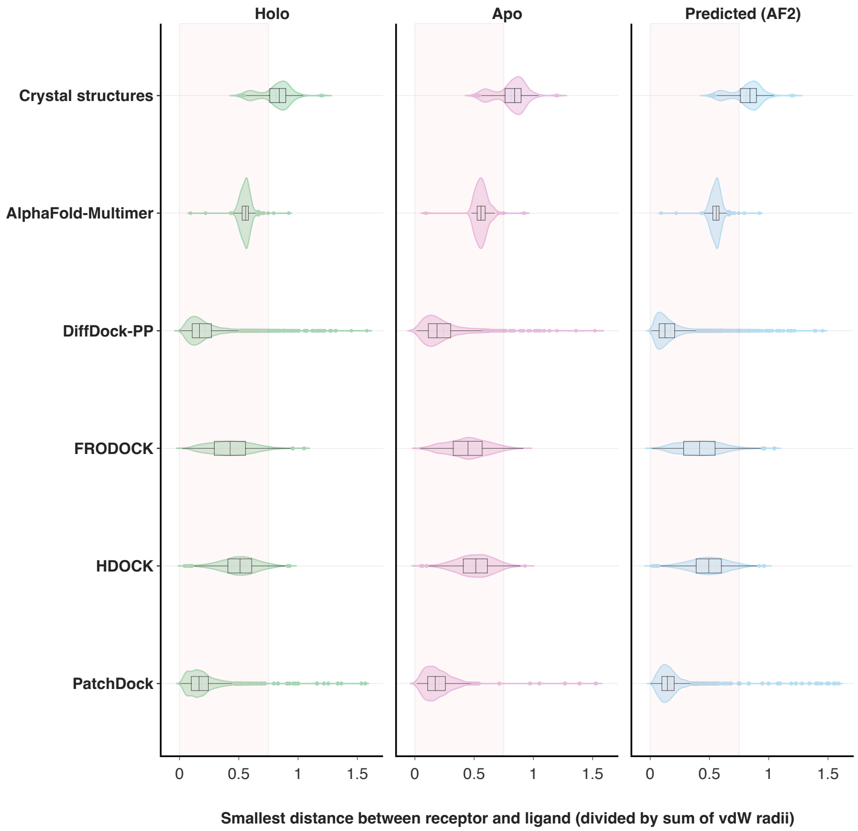


*Figure A.23.* Minimum distances between protein chains across PINDER-AF2. Distance is the smallest pairwise distance of heavy atoms of the receptor and ligand protein chains normalized by their sum of van der Waals radii. The red area highlights the rejection zone below the cutoff of 0.75 (Buttenschoen et al., 2024).

#### A.4. Interface leakage from monomer domains

##### A.4.1. ANNOTATING SIMILAR INTERFACES WITH IALIGN

While the PINDER clustering and de-leaking algorithms are designed to minimize information leakage between clusters and splits, the method is constrained by the thresholds used on the Foldseek and MMseqs2 alignment graphs. The final lDDT and sequence identity thresholds were selected based on an analysis of recall and precision at varying threshold values and selected to minimize false-positive alignments, while maximizing recall of true hits. Although reductions in the thresholds can increase recall, as the threshold gets smaller, the number of false positives quickly increases.

Rather than reducing the lDDT and sequence identity thresholds, we checked for interface similarity with an orthogonal, interface-specific alignment method from iAlign. Compared to Foldseek and MMseqs2, iAlign is not appropriate for the global all-vs-all scale due to performance limitations. Additionally, while iDist (Bushuiev et al., 2024) was introduced to mitigate the performance bottlenecks of iAlign, it was found to be insufficiently accurate to be used for identification of similar PPI interfaces that are not nearly-identical(described in A.4.2).

In order to identify similar interfaces between the dataset splits we pushed iAlign to its sensitivity limits by calibrating the cutoffs on P-value (corresponding each IS-score reported by iAlign). The P-values were adjusted for multiple testing with Benjamini-Hochberg (B-H)(Benjamini & Hochberg, 1995) correction and estimating the number of False Positives (FP) for the PINDER dataset (see Figure A.26).

The PINDER-AF2 set was curated by *removing* any test members with interface similarity to the AlphaFold training set. Any members meeting the interface similarity criteria from PINDER-XL are preserved. Systems from PINDER-XL were not removed in order to: (1) maximize the size of the test dataset, (2) evaluate the effectiveness of the PINDER splitting

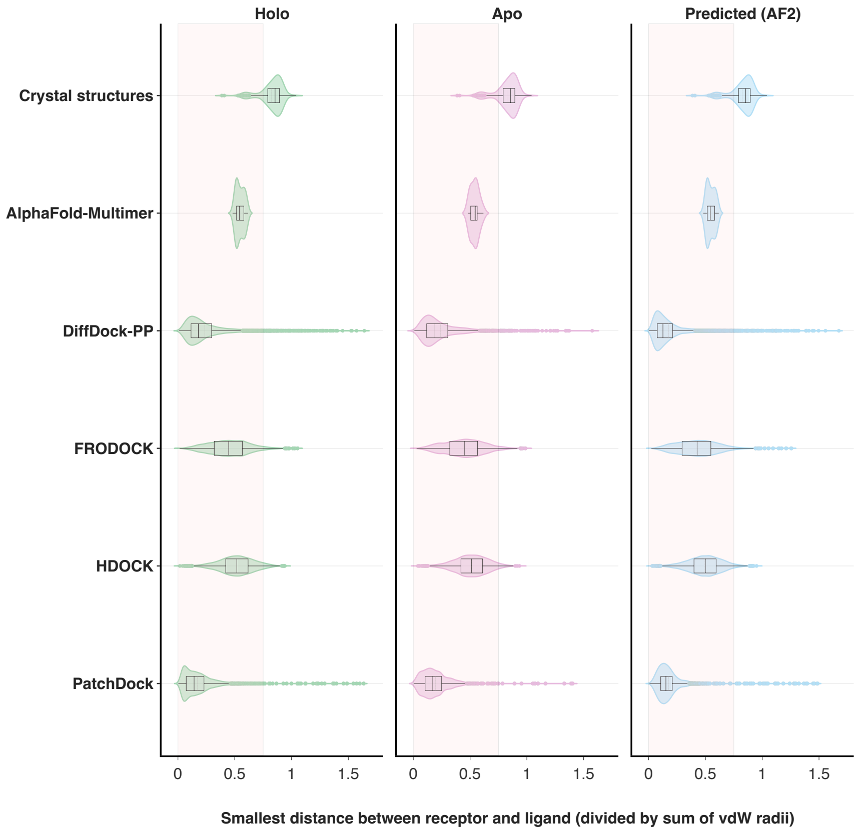


*Figure A.24.* Minimum distances between protein chains across PINDER-S. Distance is the smallest pairwise distance of heavy atoms of the receptor and ligand protein chains normalized by their sum of van der Waals radii. The red area highlights the rejection zone below the cutoff of 0.75 (Buttenschoen et al., 2024).

algorithm's ability to minimize leakage at baseline, and (3) evaluate the impact of any remaining leakage on model generalization capabilities.

##### A.4.2. COMPARISON OF IDIST VS IALIGN FOR INTERFACE DE-LEAKING PURPOSES

Despite its scalability, iDist was found to be insufficiently sensitive for the level of de-leaking we aimed to achieve. To probe its potential, a subset of PINDER-train (60k samples) was compared to the full test set (PINDER-XL) using both iDist and iAlign. At an iDist threshold of $> 0.03$, intended to indicate low similarity, the method still produced a significant number of false negatives. This insensitivity undermines its utility in accurately detecting similar interfaces. Our analysis revealed that iDist could not reliably distinguish similar from dissimilar interfaces across the spectrum of iAlign IS-scores $> 0.3$ which was the threshold utilized in de-leaking. In Figure A.27 we can see iDist values plotted against iAlign values for PINDER system pairs.

##### A.4.3. MAPPING OF FOREIGN DATASETS ONTO PINDER

Considering that PINDER systems cover all possible dimers in the biological assemblies derived from a recent snapshot of RCSB database, in order to assess diversity and leakage quality metrics of the data splits provided in other publicly-available datasets we have mapped PINDER IDs systems onto entries from these foreign datasets: DIPS-EquiDock, ProteinFlow, and PPIRef. Not every foreign system was possible to map with the major reasons being: (a) PPI from asymmetric units that most likely represent crystal packing and are not included in PINDER, which is foreign dataset limitation; (b) oligomeric systems with more than two chains that did not map to PINDER unambiguously, which is PINDER limitation.

**DIPS-EquiDock splits mapping** While the original DIPS dataset used DB5 as a test set, we downloaded the DIPS dataset

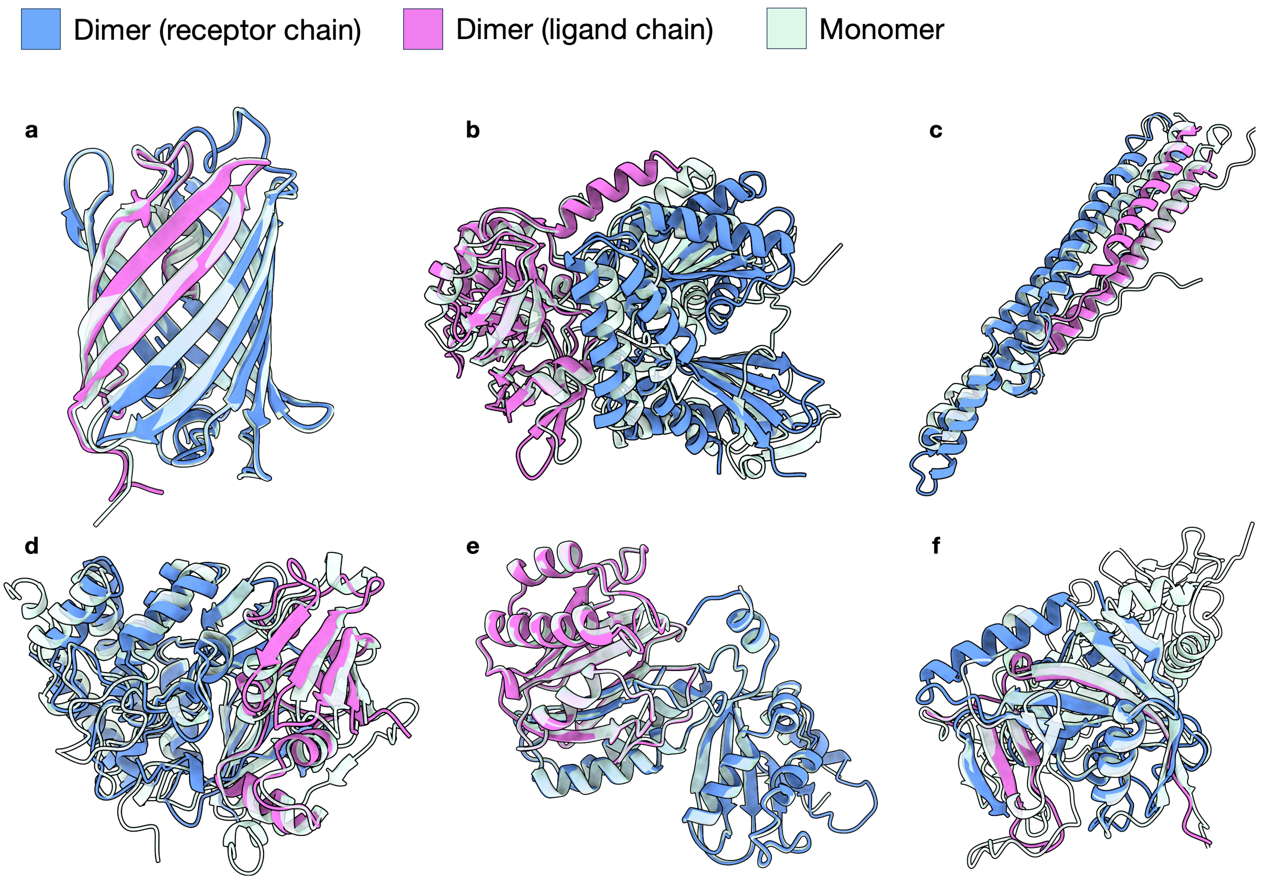


*Figure A.25.* **Examples of monomers with intra-chain contacts similar to dimer interfaces**. AlphaFold-Multimer predictions for **a-e** are all "High" CAPRI quality with DockQ above 0.8. **f** is incorrectly predicted with a DockQ of 0.02. **a**, Monomer of green fluorescent protein (PDB 6LZ2) superimposed to dimer (PDB 3LF4). **b**, Fumarate hydratase 2 monomer (PDB 6UQM) superimposed to two-subunit Fumarate hydratase apo-protein dimer complex (PDB 7XKY). **c**, ESX-1 secretion-associated protein EspB monomer (PDB 8AKO) superimposed to the dimer PE/PPE protein complex (PDB 2G38). **d**, Toxin-antitoxin system toxin HipA family monomer (PDB 4PU4) superimposed to toxin Lpg2370 (HipT Lp) and the antitoxin Lpg2369 (HipS Lp) dimer complex (PDB 7VKB). **e**, Ornithine Acetyltransferase monomer (PDB 1VZ7) superimposed to acyl-enzyme dimer complex in catalytic conditions (PDB 2VZK). **f**, Poly [ADP-ribose] polymerase 2 (PAR) monomer (PDB 6X0L) superimposed to fragmentized poly [ADP-ribose] polymerase tankyrase-2 heterodimer (PDB 8B6M).

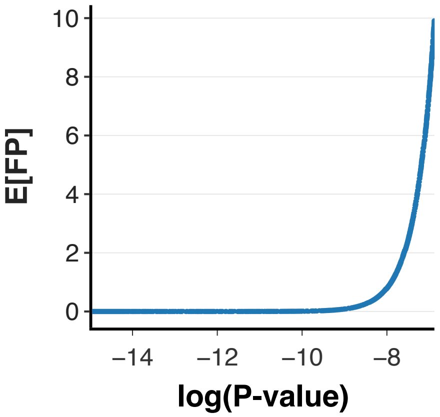


*Figure A.26.* **Calibration of expected False Positives at sensitive iAlign p-value thresholds**

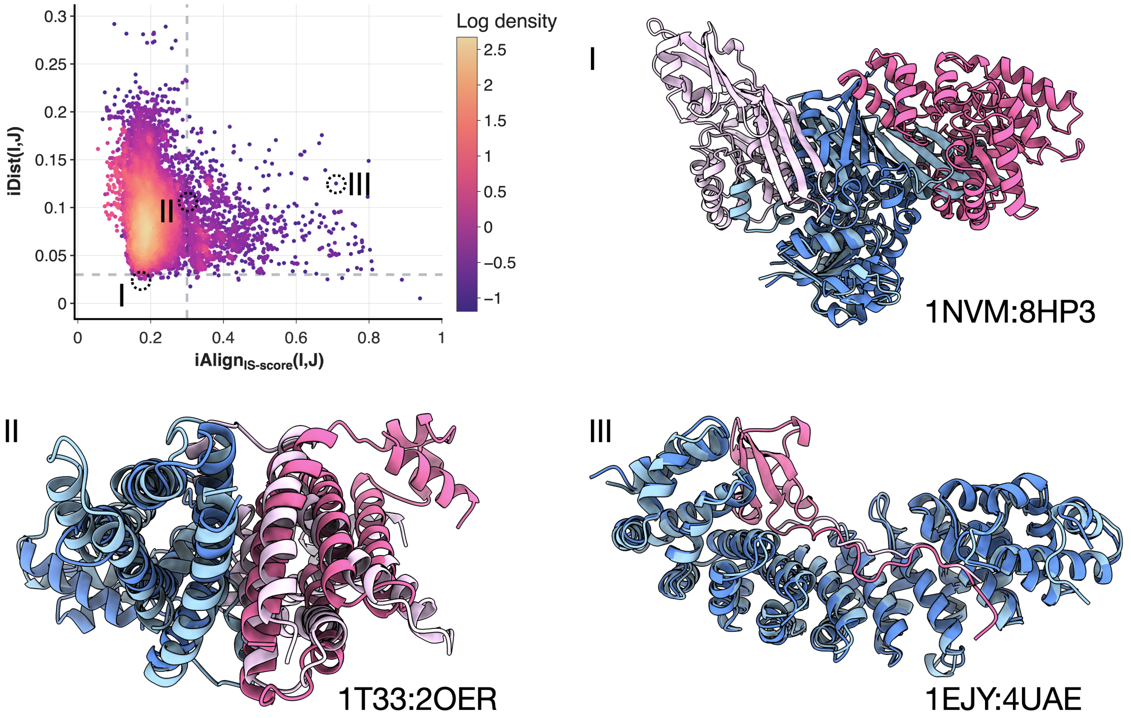


*Figure A.27.*
I: An example of a false positive from iDist that is negative in iAlign.
II: An example of a false negative from iDist that is close to the borderline, but falls within our definition of leakage using iAlign. This illustrates how important it is to consider IS-scores in the range ˜0.25-0.3.
III: An example of a false negative that has an extreme delta between iAlign and iDist. This demonstrates how short peptides or less-obvious leakage may be missed by iDist but detected by iAlign.

used in the EquiDock project with 40,143 entries split into 38,296 in train, 928 in val, and 919 in test. These entries were mapped onto 30,938 PINDER ids in train, 745 in val, and 757 in test, the rest of the chains were not mapped mostly due to interactions extracted from asymmetric units and not from the biological assemblies.

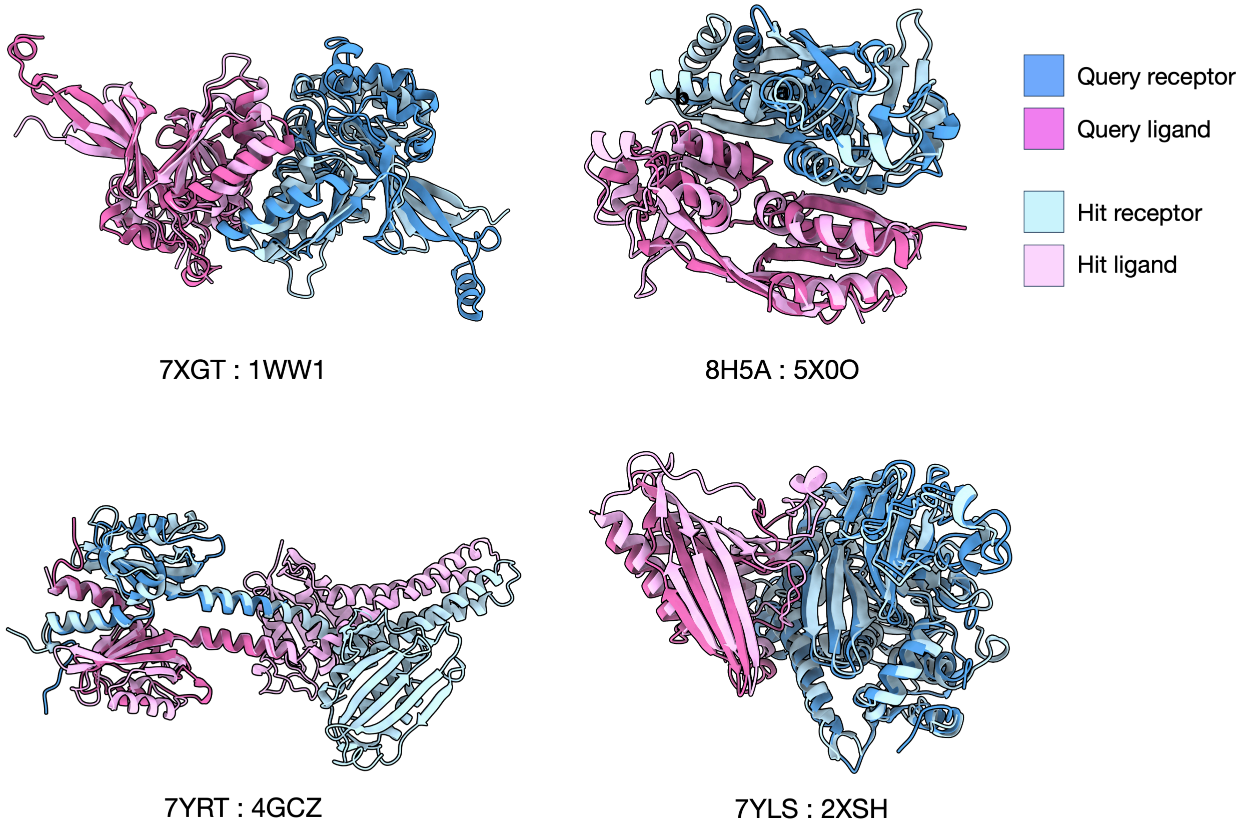


*Figure A.28.* **Examples of interface data leakage between PINDER-XL and AlphaFold-Multimer training set**. All examples were detected during construction of the PINDER-AF2 subset using iAlign and were removed from the eligible PINDER-AF2 systems.

**ProteinFlow splits mapping** We obtained ProteinFlow dataset release "20230102_stable", which contains 9,818 (977) test, 252,680 (19,471) train and 12,466 (1,017) val entries. The numbers in brackets indicate ProteinFlow clusters in each set. Out of which 250,177 train set, 14,357 val set, and 7119 test set ProteinFlow entries were successfully mapped onto PINDER IDs, leaving 679,070 PINDER IDs without a match to ProteinFlow.

**PPIRef splits mapping** While PPIRef provides multiple datasets, we have selected the non-redundant dataset PPIRef50K constructed at 6Å distance cutoff that had the most similar interfaces clustered and de-duplicated using the iDist methodology ("ppiref_6A_filtered_clustered_04"). The dataset includes 51,755 dimers. PPIRef does not provide a split for this dataset, however, considering that a 80/20/20 random splitting was used to train PPIformer, we generated random PPIRef splits with the same proportions, resulting in 41,403 train, 5,176 val, and 5,176 test entries. These generated splits were mapped to PINDER IDs, leading to 25,049 train, 3,092 val, and 3,162 test IDs mapped. The 45% loss in mapping is largely explained by the fact that interactions in PPIRef were extracted from asymmetric units and not biological assemblies.

###### A.4.4. COMPARISON TO OTHER DATASETS

| | | PINDER | | DIPS-EquiDock | | ProteinFlow | | PPIRef | |
|---|---|---|---|---|---|---|---|---|---|
| | Metric | Test (%) | Val (%) | Test (%) | Val (%) | Test (%) | Val (%) | Test (%) | Val (%) |
| Diversity | Structure-based cluster | 5.65 | 5.66 | 0.69 | 0.64 | 1.83 | 1.84 | 6.85 | 6.62 |
| | ECOD single-chain | 16.66 | 19.11 | 3.36 | 2.88 | 6.20 | 6.64 | 20.48 | 20.05 |
| | ECOD chain-pair | 8.16 | 9.17 | 1.64 | 1.42 | 3.72 | 3.91 | 10.89 | 10.49 |
| | Pfam clan | 52.20 | 57.80 | 19.32 | 17.80 | 29.66 | 33.39 | 58.64 | 54.92 |
| | Unique ECOD pairs | 30.95 | 33.15 | 4.76 | 7.26 | 37.28 | 11.72 | 4.43 | 3.96 |
| Leakage | UniProt pair | 2.76 | 3.27 | 42.93 | 48.99 | 3.91 | 5.42 | 36.94 | 35.72 |
| | ECOD pair | 6.34 | 8.02 | 63.94 | 75.54 | 26.07 | 52.34 | 45.17 | 45.28 |
| | iAlign interface pair | 12.63 | 11.13 | 83.36 | 84.03 | 24.57 | 11.99 | 63.47 | 15.62 |

*Table A.1.* **Diversity and leakage metrics for PINDER compared with other publicly available datasets.** Metrics summarizing the diversity and amount of leakage for systems in the the Test (PINDER-XL, for PINDER) and Val sets relative to systems in the Train set are shown as percentages for selected datasets. Metrics are defined and discussed in Section A.2.

Figure A.30 depicts the distributions of various properties in the PINDER test set compared to test splits from other datasets. Overall, while the distributions are similar for planarity, number of intermolecular contacts, and resolution (Figure A.30A, B, D) the PINDER test split has a wider range of PPIs with varying properties, allowing for a more representative and comprehensive evaluation. The physiological probability of the interface, as predicted by PRODIGY-cryst (Jiménez-García et al., 2019) is always high for PINDER test PPIs (Figure A.30C) as we only consider physiological interfaces. We restrict the test set to dimers, with a near-even split between homo and heterodimers in PINDER-S, but a higher proportion of homodimers in the overall PINDER-XL set (Figure A.30E), as considering binary interactions from multimeric complexes can lead to cases where the reported chains have minimal contact and require the presence of other chains for stabilizing the contact (Figure 2B).

##### A.5. PINDER dataloader

PINDER provides its dataloader to facilitate the loading of subsets of the dataset for training and evaluating protein-protein docking models. The dataloader is based on iterators over a core abstraction called "Pinder System", which provides a collection of structural data associated with an entry in the PINDER database. Each Pinder System exposes the ground-truth crystal structure as *holo* receptor and ligand, and where available, the *apo* receptor and ligand, and finally the predicted receptor and ligand (currently from AlphaFold Protein Structure Database). Each structure is represented by a "Structure" object, which provides access to various structural features like coordinates, residues, atoms, and sequence and structural utilities.

The dataloader offers several features to streamline the data loading process. Users can obtain a collection of monomers associated with a PINDER entry, classify system difficulty based on conformational shifts between *apo* and *holo* states, and access various structural features like coordinates, residues, atoms, and sequence and structural utilities. Additionally, the dataloader allows for filtering datasets to construct specific data mixes based on the user specific task, such as the presence

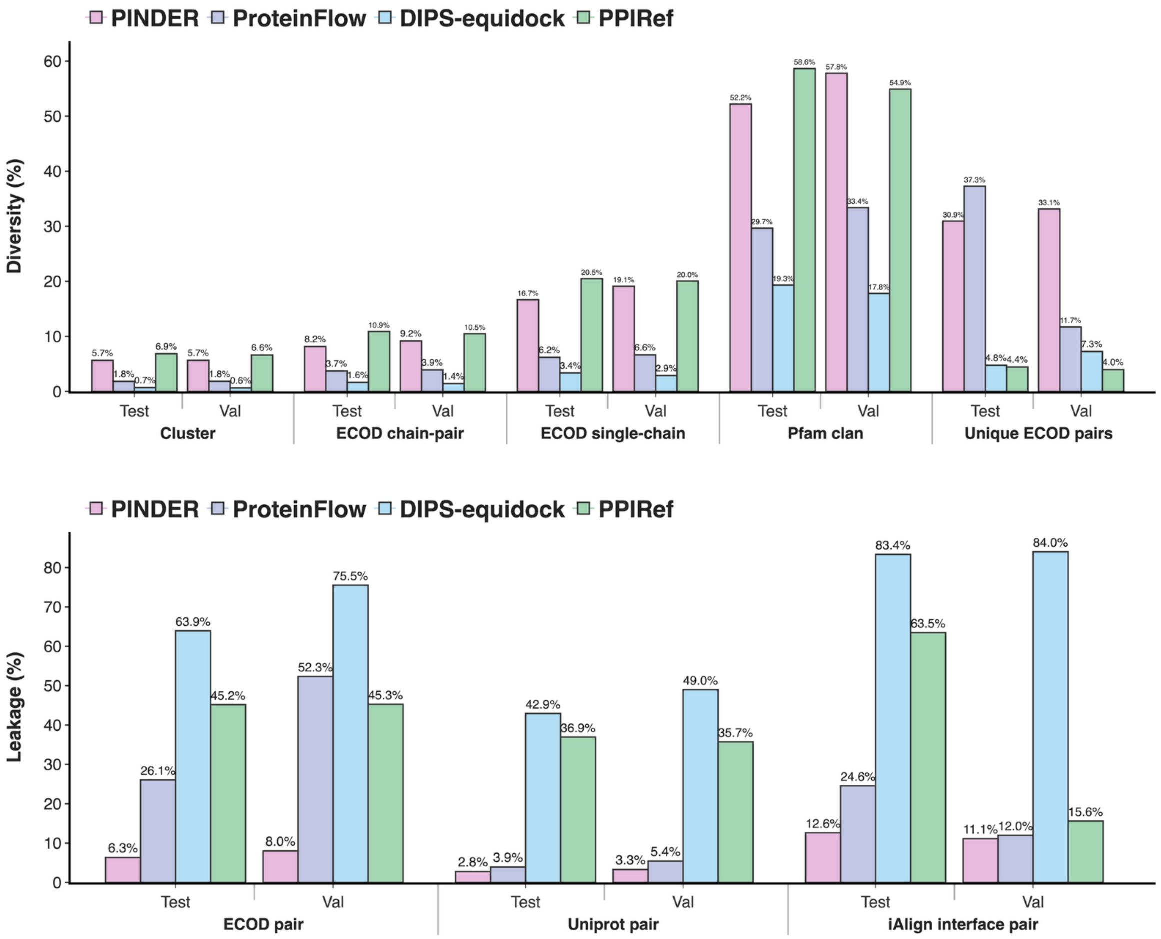


*Figure A.29.* Diversity and leakage metrics for PINDER compared with other publicly available datasets. Metrics summarizing the diversity and amount of leakage for systems in the the Test (PINDER-XL, for PINDER) and Val sets relative to systems in the Train set are shown as percentages for selected datasets. Metrics are defined and discussed in Section A.2.

of certain atom types, sequence identity, or structural properties. Users can also create generators for specific data mixes and apply a collection of filters. The dataloader also supports loading PPI data in the form of PyTorch Geometric hetero-data allowing to create standard PyTorch dataloaders for ”ready to use” integration into machine learning workflows.

##### A.5.1. FILTERING OPTIONS

PINDER provides a wide array of filters to select specific subsets of data based on various criteria. These filters can be applied to both the entire protein complex (base filters) and individual protein chains (subunit filters). Some of the key filter categories include:

1. **Missing Data Filters:** Filter out entries with missing *holo* structures or specific chain types (e.g., *apo*, predicted).

2. **Interface Property Filters:** Filter based on interface characteristics such as the number of contacts, planarity, and buried surface area.

3. **Structural Property Filters:** Filter based on structural properties like elongation, number of connected components, and presence of gaps.

4. **Metadata Filters:** Filter based on metadata annotations such as experimental method, resolution, presence of antibodies or enzymes, and more.

5. **Sequence Similarity Filters:** Filter based on sequence identity between *holo* and *apo*/predicted structures.

###### A.5.2. DATA AUGMENTATION

The PINDER dataloader module enables data augmentation by allowing the loading of multiple conformations (*apo*, *holo*, or predicted) for each protein chain in any combination. This means that for a single PPI, the dataloader can generate multiple training examples by combining different conformations of the receptor and ligand chains. As described in Section 2.2, the *holo* dimers were paired with *apo* and *predicted* monomers, selecting the most structurally diverse representatives as canonical (see Table 8). However all available alternative monomers are used, PINDER expands the total training dataset to a size of 25 million structures. This data augmentation strategy increases the diversity of the training data and exposes models to a wider range of conformational variations, which can improve their generalization capabilities and robustness.

##### A.6. Retraining DiffDock-PP

We re-train DiffDock-PP using the NVIDIA BioNeMo Framework (bio) (FW) with an identical model size of 1.6M parameters with the original DiffDock-PP model (Ketata et al., 2023). A Fused Adam optimizer was used a with learning rate of 0.001 and ReduceLROnPlateau scheduler with patience of 20 epochs. A new feature of DiffDock-PP in the BioNeMo FW is an Adaptive Batch Sampler during training. Due to the variable size of protein-protein complexes, the memory requirements when loading the heterograph can vary. This algorithm pre-computes the memory requirement of each complex, and shuffles the batches to accommodate the memory overhead. Effective batch size is around 20. All models were trained on eight 80GB A100 GPUs for 200 epochs, and fully converge in 24 hours.

###### A.6.1. SAMPLING OF TRAINING EXAMPLES

To create the DiffDock-PP training datasets for the structure and sequence splits, we employed a filtering and sampling scheme, and condensed the training dataset size to a DIPS-plus comparable level, while allowing for efficient training of all models within a reasonable time frame. This involved sampling a non-redundant training set in the following manner:

- Filter - **num_atom_types** $\geq$ **4:** Ensures that all proteins have all backbone atom types.
- Filter - **max_var** $\leq$ **0.98**: Excludes elongated chains helps to focus on more globular protein structures.
- Filter - **length_resolved** $>$ **40:** Avoids short chains and peptides.
- Sample - **top 3 cluster representatives:** The representative selection was based on *apo* and predicted availability, X-RAY determination method, and resolution to prioritize high-quality structures.

By implementing this filtering and sampling strategy, the dataset was effectively reduced to a manageable size, while still retaining a representative and diverse subset of protein structures for the DiffDock-PP training.

##### A.7. Parameters of classical docking methods

###### A.7.1. FRODOCK

The FroDock (version `3.12`) docking process involves several stages, each with specific parameters (A.2). In Stage 1 van der Waals potential map of receptor is created using the command `frodockgrid` with default parameter -m 0, followed by Stage 2 that generates the receptor's desolvation potential map with parameters `-m 1 -t A`. Stages 3 and 4 create both the receptor's and ligand's desolvation potential map using `-m 3`. Stage 5 performs the docking using `frodock` with default parameters. Stage 6 clusters and visualizes top 40 predictions using `frodockcluster --nc 40` and `frodockview -r 1-40`, also with default parameters.

###### A.7.2. HDOCK

HDock docking protocol involved running HDOCKlite `v1.0` with the default parameters. The only specified parameter was the number of solutions: `-nmax 40`. Default parameters include an angle interval of 15° that is used for rotational sampling, and a spacing of `1.2Å` adopted for FFT-based translational search. For each rotation, the top `10` translations with the best shape complementarities from the FFT search are optimized by knowledge-based scoring functions. HDocklite does not support templates.

##### A.7.3. PatchDock

The PatchDock docking protocol involves running several stages with specific parameters (see table A.3). The first stage generates the parameters using `buildParams.pl` with an RMSD value of `4.0`. The second stage creates the receptor electrostatic potential map using `patch_dock`. The final stage generates the top `40` solutions using `transOutput.pl`.

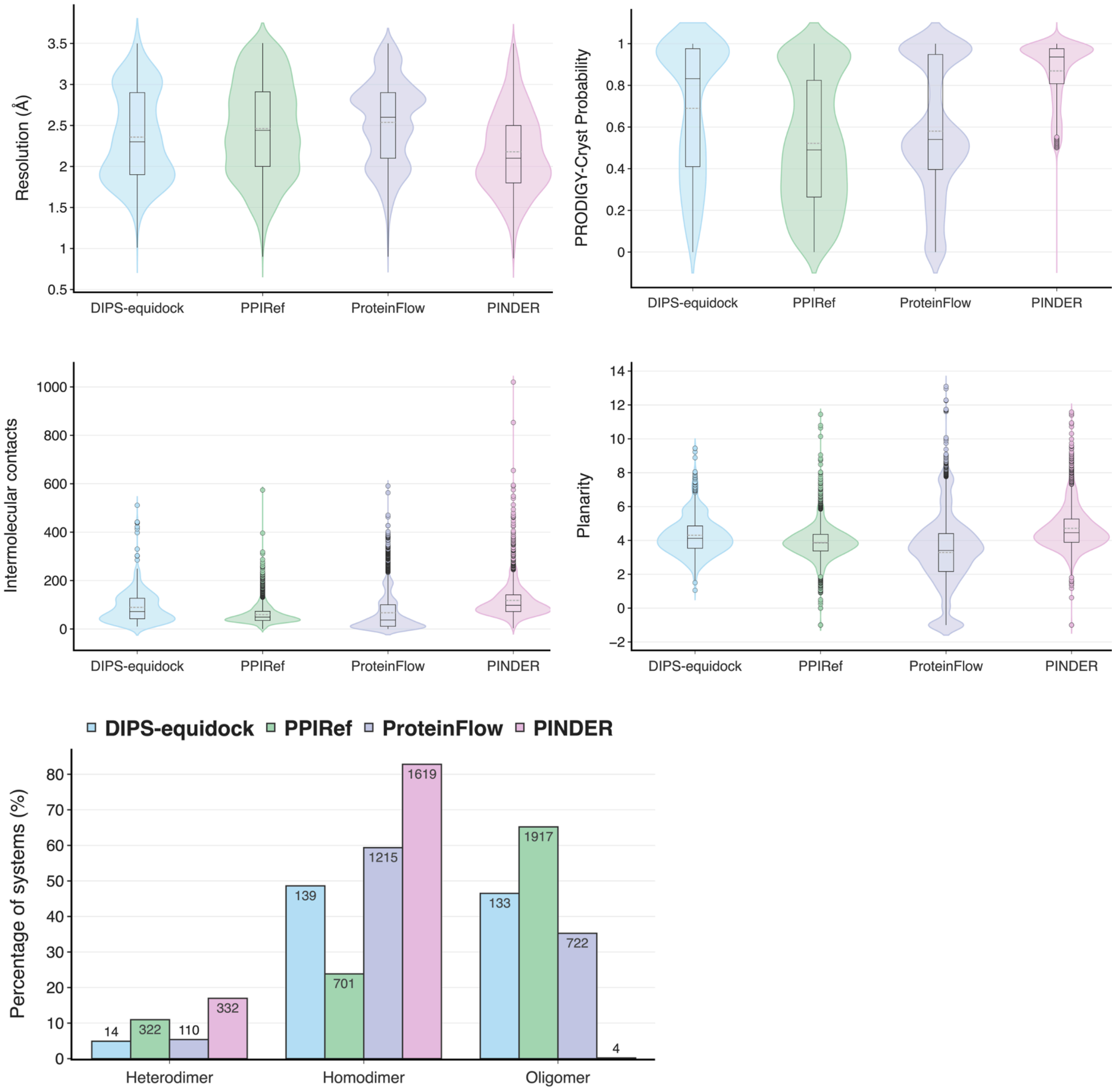


*Figure A.30.* Violin plots depicting, for the test PPIs of the different datasets, the distributions of (A) Resolution; (B) Physiological interface probability; (C) Number of intermolecular contacts; (D) Planarity and (E) Percentage of different oligomeric states

#### A.8. Predicted structure case studies

Based on the PINDER leaderboard per system metrics, we sampled and visualized example cases from all evaluated methods for each input monomer used. Cases were selected based on the highest average DockQ (easiest), lowest average DockQ (hardest), highest variance DockQ, and examples where a single method performed better than the rest.

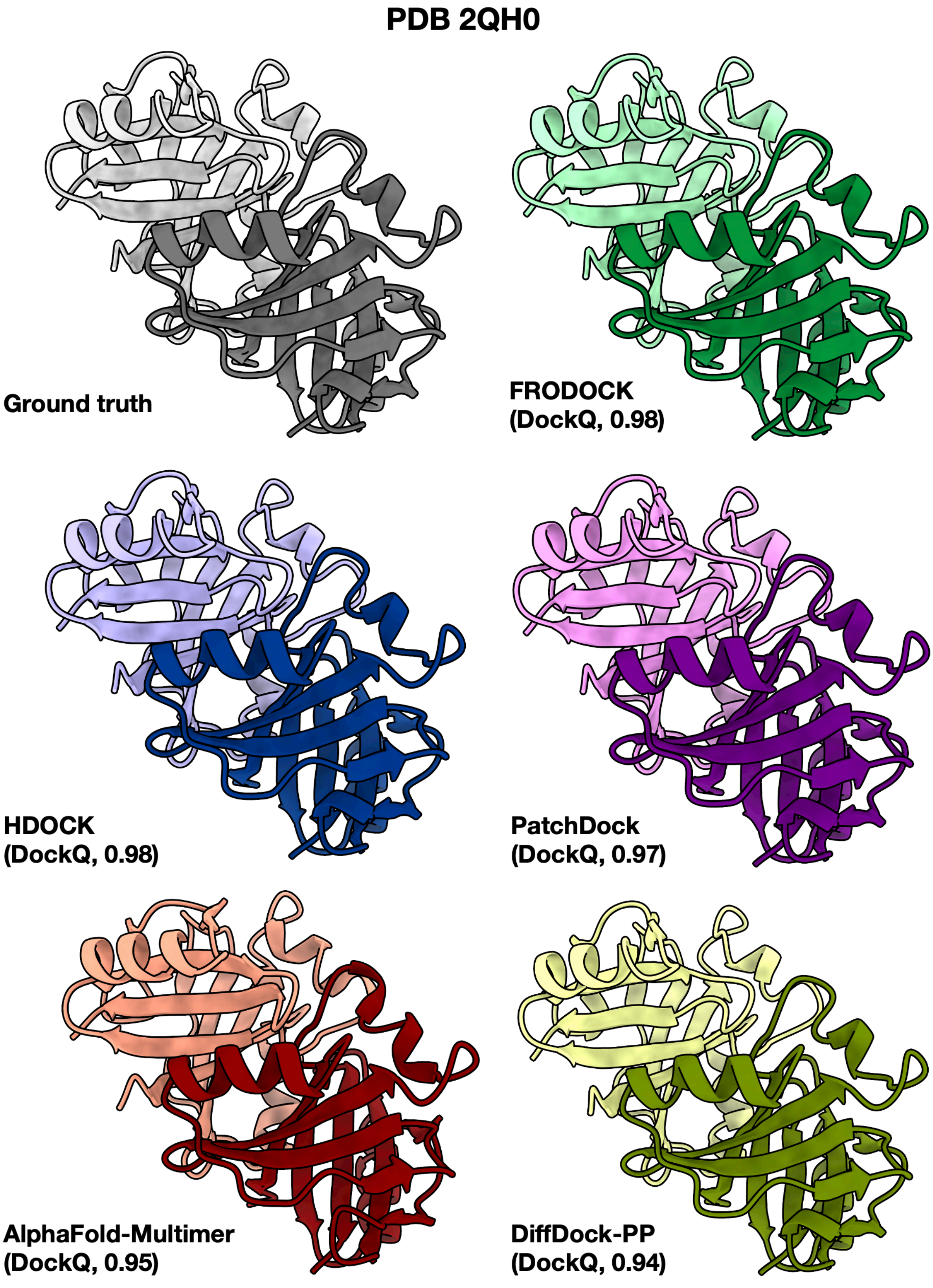


*Figure A.31.* Easiest system using bound *holo* structures, based on highest mean oracle DockQ across methods.

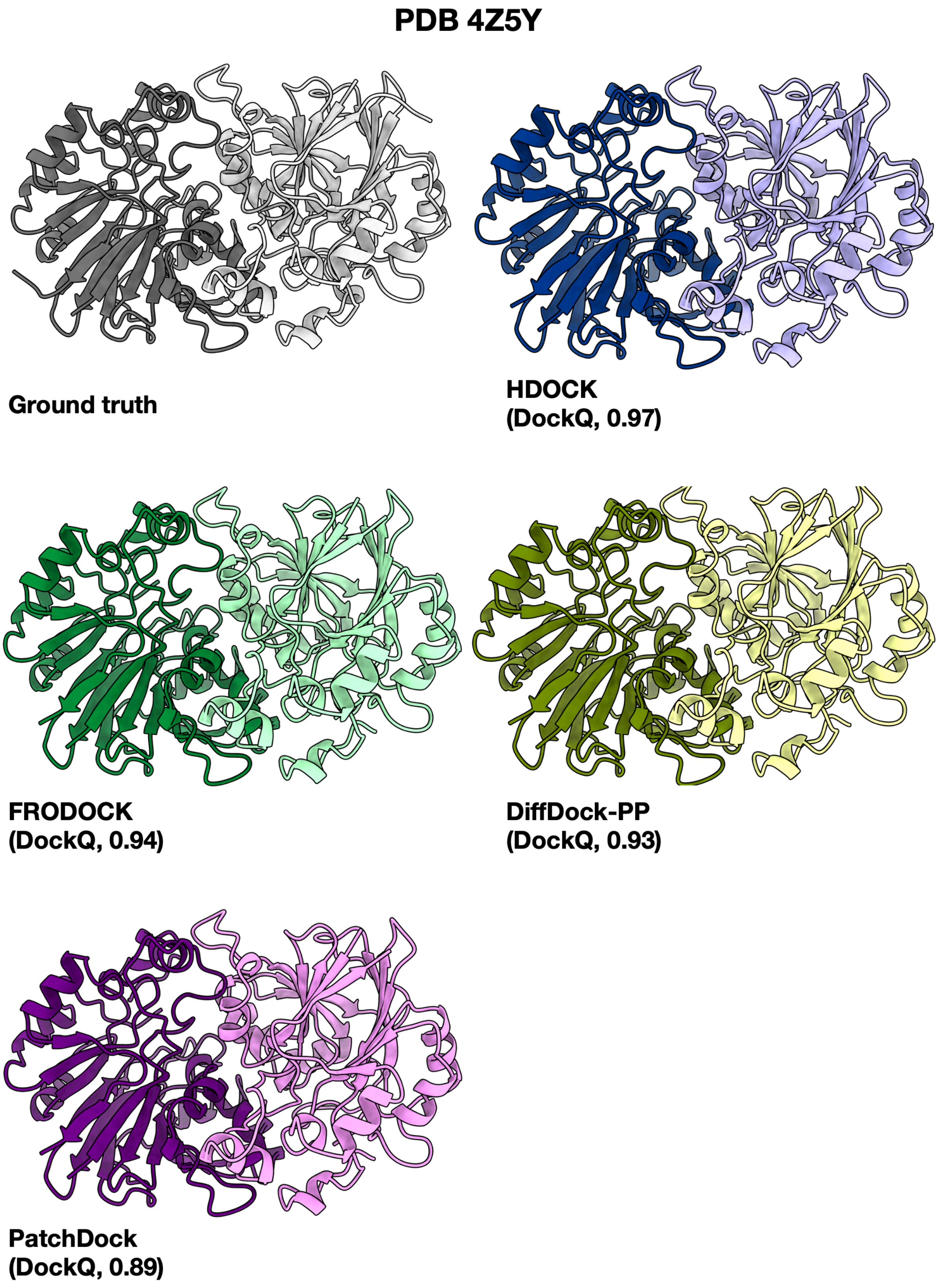


*Figure A.32.* Easiest system using unbound *apo* input structures, based on highest mean oracle DockQ across methods.

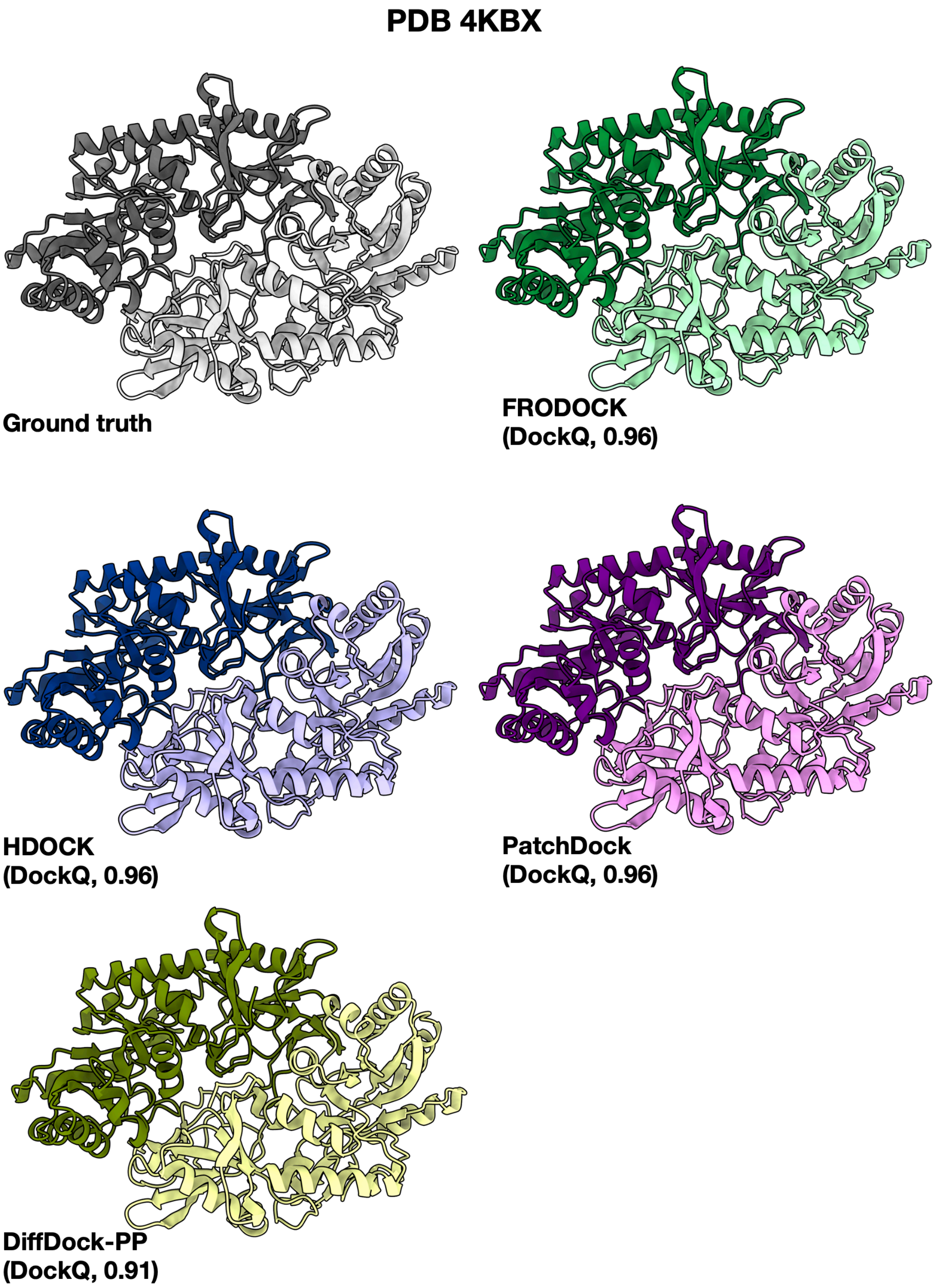


*Figure A.33.* Easiest system using unbound predicted input structures, based on highest mean oracle DockQ across methods.

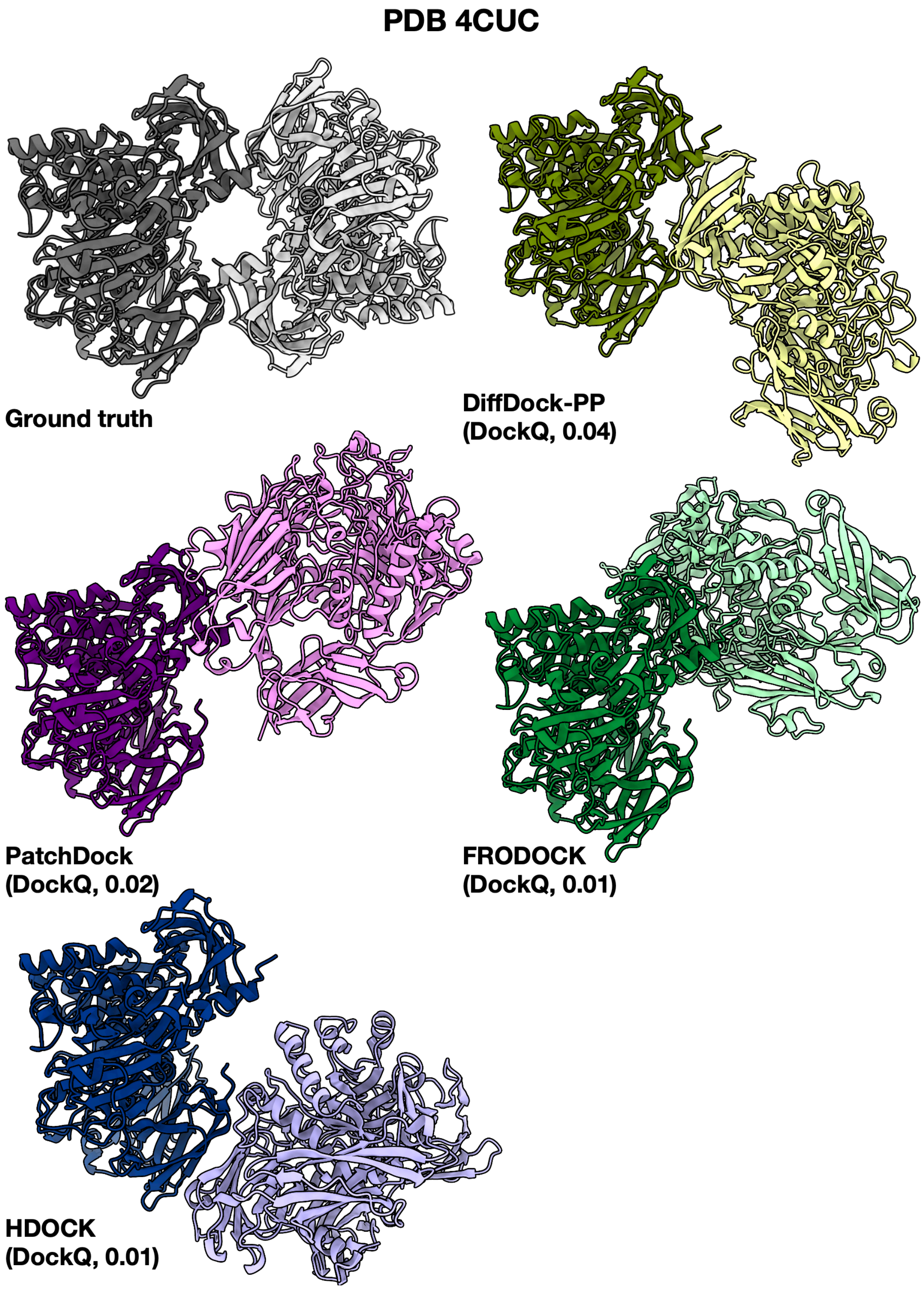


*Figure A.34.* Hardest system using unbound *apo* structures, based on lowest mean oracle DockQ across methods.

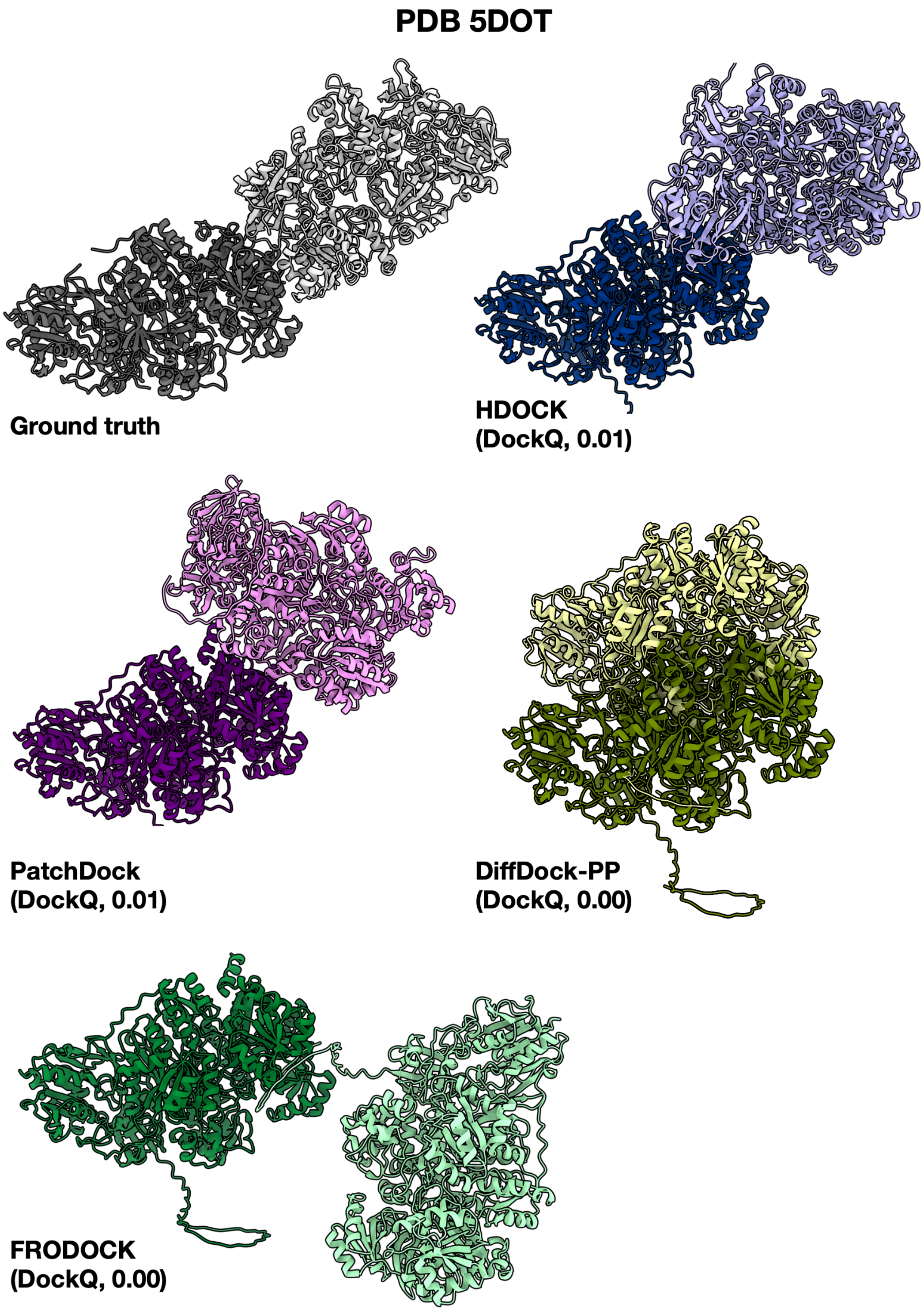


*Figure A.35.* Hardest system using unbound predicted structures, based on lowest mean oracle DockQ across methods.

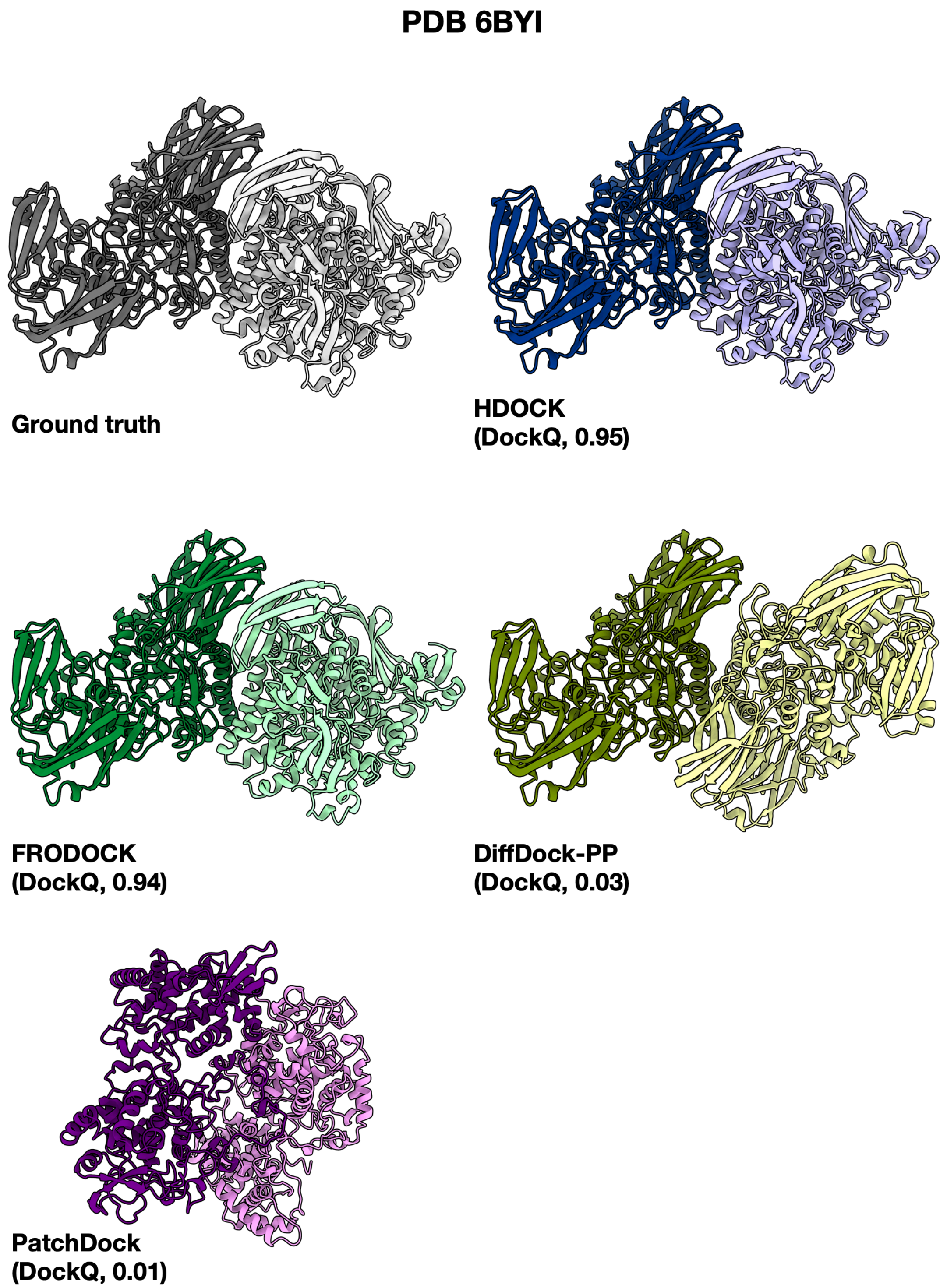


*Figure A.36.* System with highest variation in oracle DockQ across methods using unbound *apo* input structures.

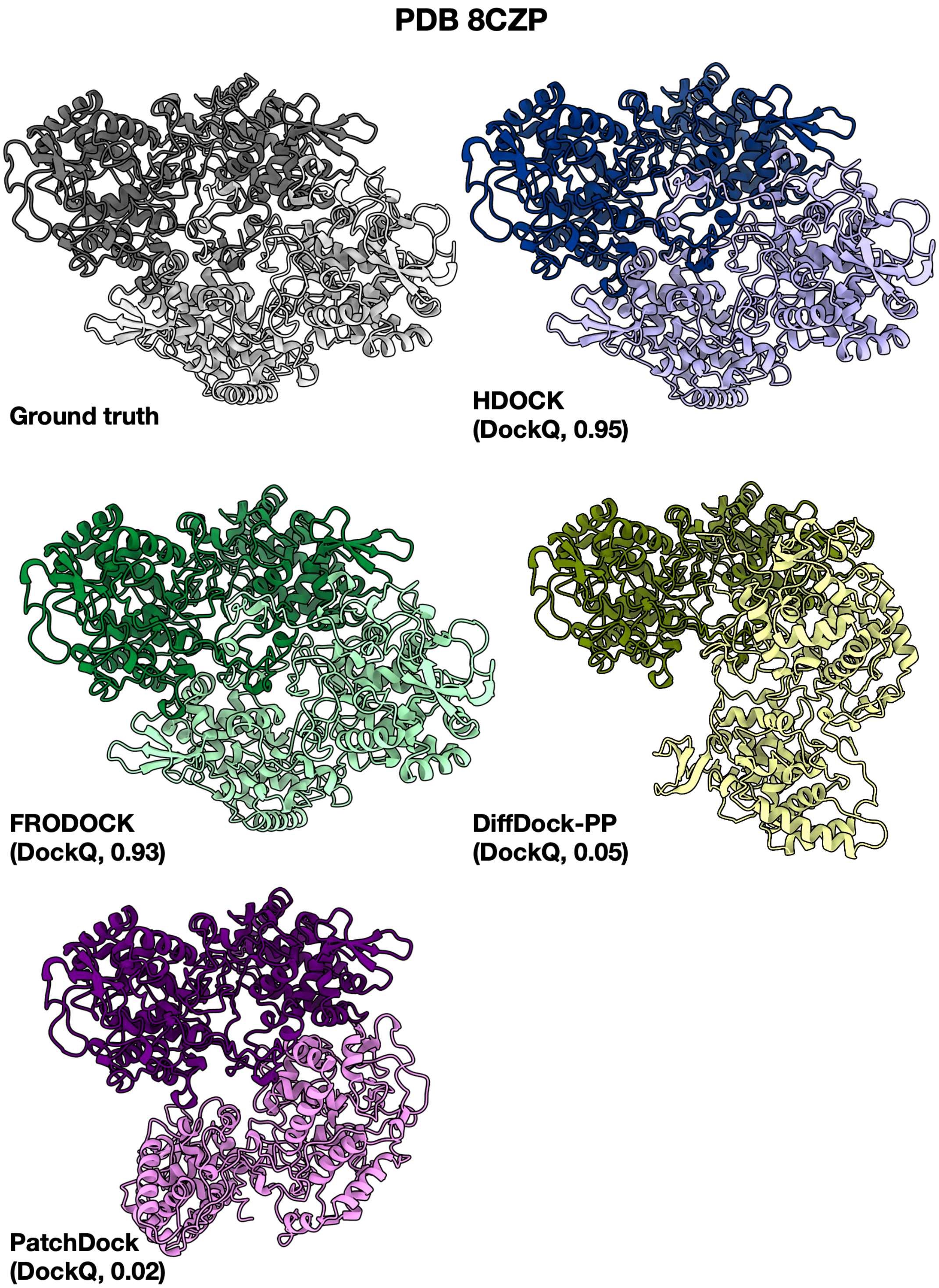


*Figure A.37.* System with highest variation in oracle DockQ across methods using unbound predicted input structures.

##### A.9. Singleton Success

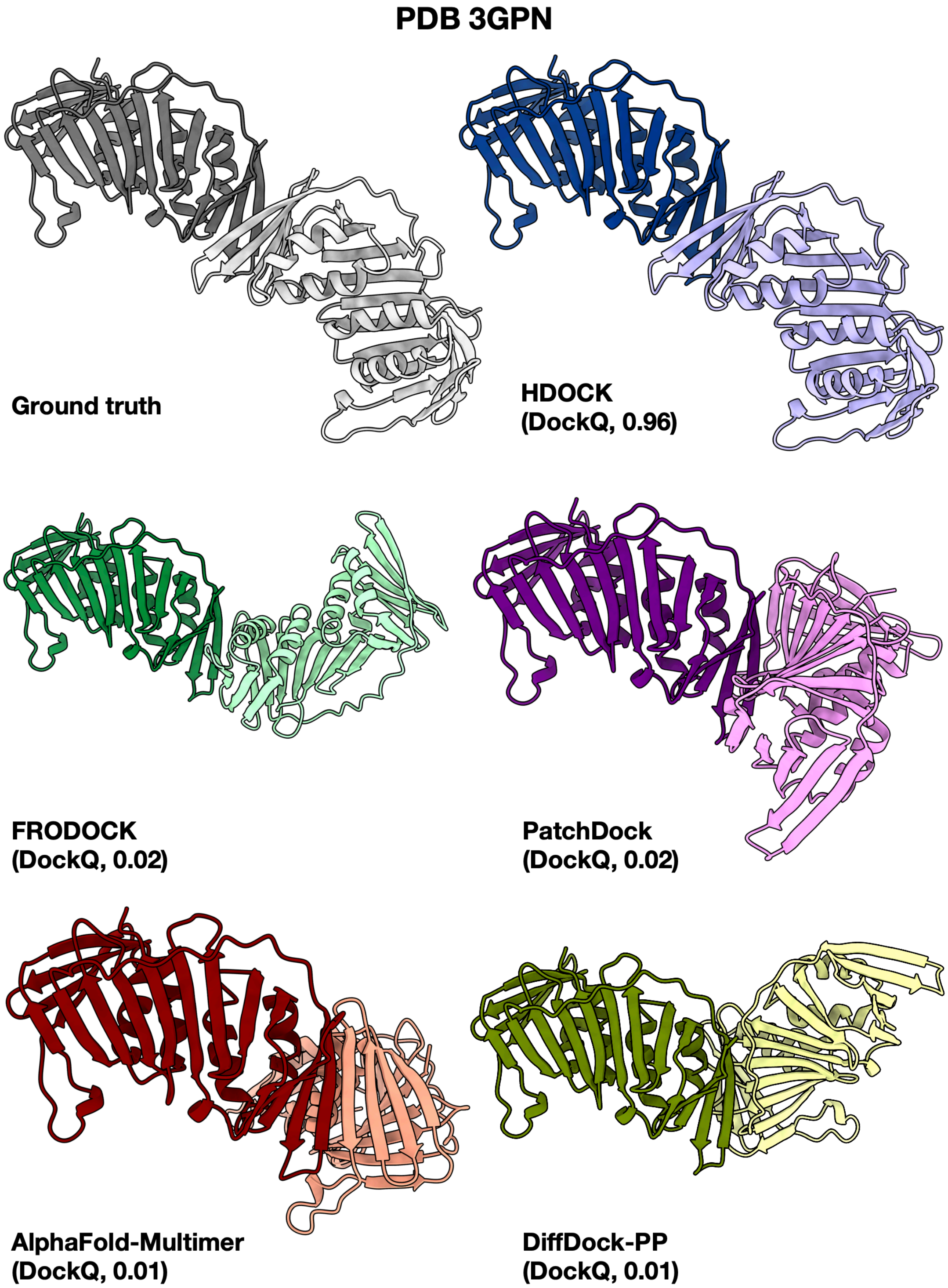


*Figure A.38.* System with only one method (HDOCK) producing a successful prediction using bound *holo* input structures.

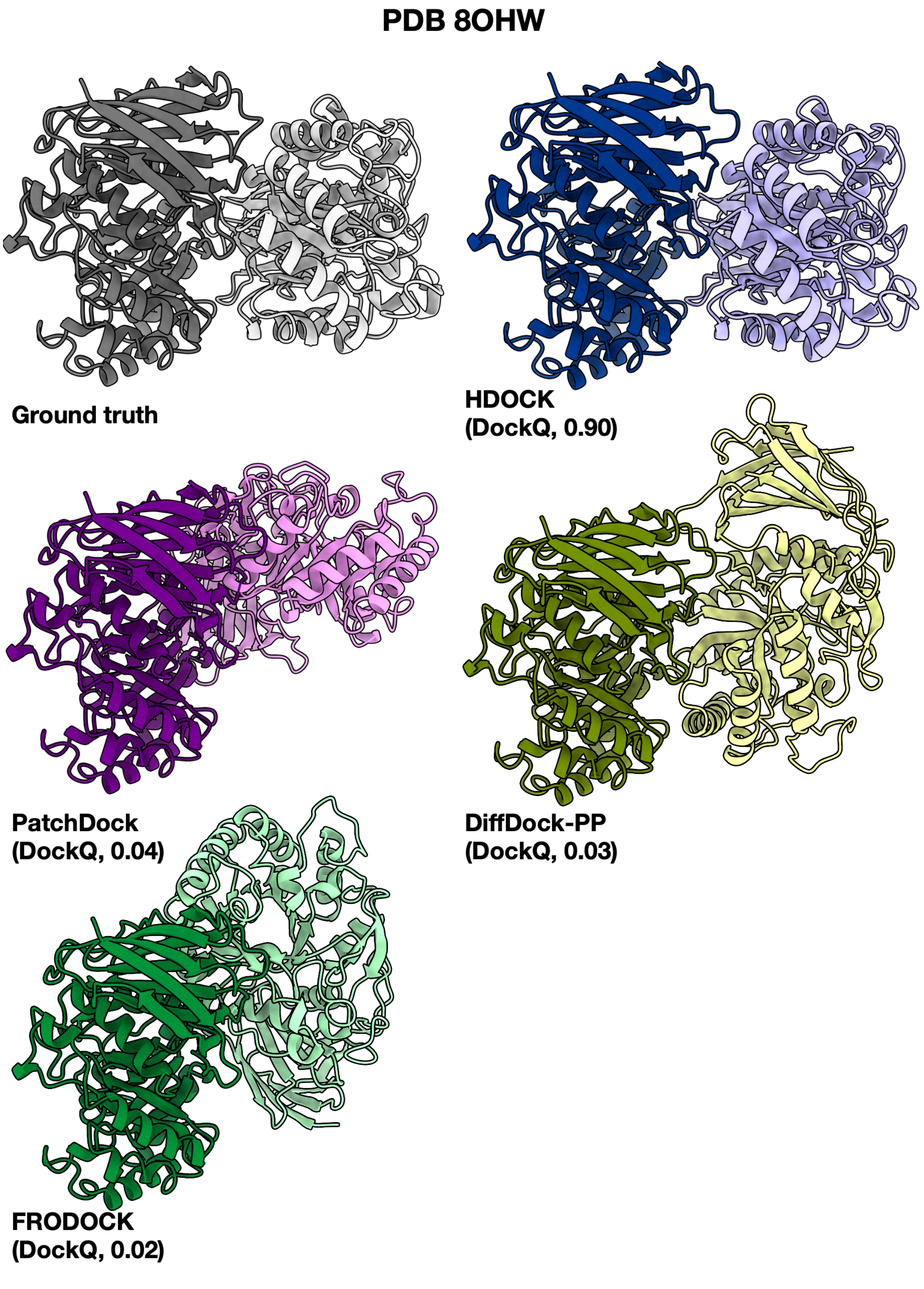


*Figure A.39.* System with only one method (HDOCK) producing a successful prediction using unbound *apo* input structures.

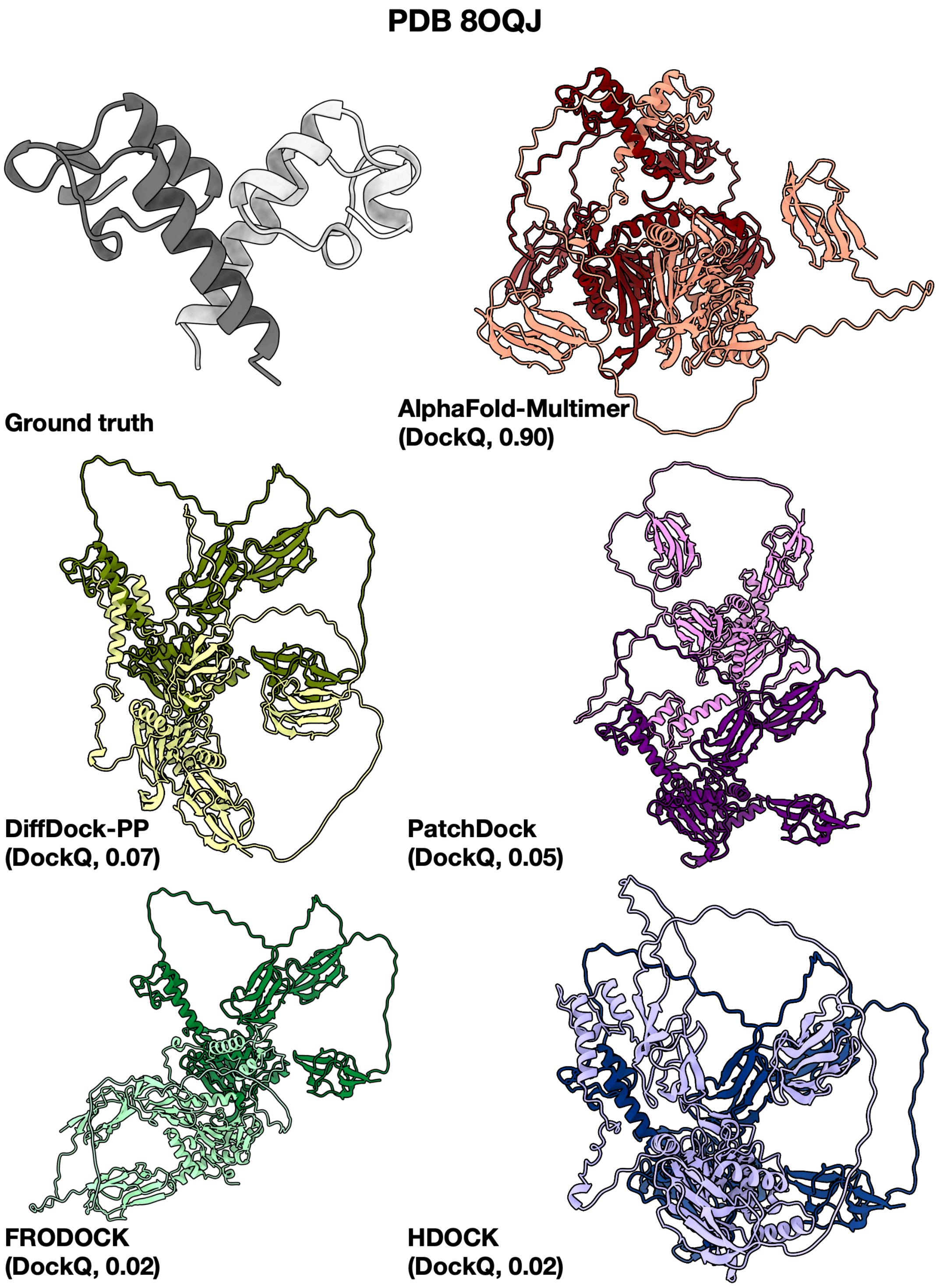


*Figure A.40.* System with only one method (AlphaFold-Multimer) producing a successful prediction using unbound predicted input structures.

##### A.10. Annotations

Here we detail a full list of annotations collected for systems included in PINDER.

##### A.10.1. Computed annotations

- **label**: Classification of the interface as likely to be biologically-relevant or a crystal contact, annotated using PRODIGY-cryst. PRODIGY-cryst uses machine learning to compute bio-relevant/crystal contact propensity based on Intermolecular contact types and Interfacial link density.
- **probability**: Probability that the protein complex is a true biological complex.
- **length1**: The number of amino acids in the first (receptor) chain.
- **length2**: The number of amino acids in the second (ligand) chain.
- **length_resolved_1**: The structurally resolved (CA) length of the first (receptor) chain in amino acids.
- **length_resolved_2**: The structurally resolved (CA) length of the second (ligand) chain in amino acids.
- **number_of_components_1**: The number of connected components in the first (receptor) chain (contiguous structural fragments)
- **number_of_components_2**: The number of connected components in the second (receptor) chain (contiguous structural fragments)
- **link_density**: Density of contacts at the interface as reported by PRODIGY-cryst. Interfacial link density is defined as the number of interfacial contacts normalized by the maximum possible number of pairwise contacts for that interface. Values range between 0 and 1, with higher values indicating a denser contact network at the interface.
- **planarity**: Defined as the deviation of interfacial $C_\alpha$ atoms from the fitted plane. This interface characteristic quantifies interfacial shape complementarity. Transient complexes have smaller and more planar interfaces than permanent and structural scaffold complexes.
- **max_var_1**: The maximum variance of coordinates projected onto the largest principal component. This allows the detection of long end-to-end stacked complexes, likely to be repetitive with small interfaces (receptor chain).
- **max_var_2**: The maximum variance of coordinates projected onto the largest principal component. This allows the detection of long end-to-end stacked complexes, likely to be repetitive with small interfaces (ligand chain).
- **num_atom_types**: Number of unique atom types in structure. This is an important annotation to identify complexes with only $C_\alpha$ or backbone atoms.
- **n_residue_pairs**: The number of residue pairs at the interface.
- **n_residues**: The number of residues at the interface.
- **buried_sasa**: The buried solvent accessible surface area upon complex formation.
- **intermolecular_contacts**: The total number of intermolecular contacts (pair residues with any atom within a 5Å distance cutoff) at the interface. Annotated using PRODIGY-cryst.
- **charged_charged_contacts**: Denotes intermolecular contacts between any of the charged amino acids (E, D, H, K). Annotated using PRODIGY-cryst.
- **charged_polar_contacts**: Denotes intermolecular contacts between charged amino acids (E, D, H, K, R) and polar amino acids (N, Q, S, T). Annotated using PRODIGY-cryst.
- **charged_apolar_contacts**: Denotes intermolecular contacts between charged amino acids (E, D, H, K) and apolar amino acids (A, C, G, F, I, M, L, P, W, V, Y). Annotated using PRODIGY-cryst.
- **polar_polar_contacts**: Denotes intermolecular contacts between any of the charged amino acids (N, Q, S, T). Annotated using PRODIGY-cryst.
- **apolar_polar_contacts**: Denotes intermolecular contacts between apolar amino acids (A, C, G,F, I, M, L, P, W, V, Y) and polar amino acids (N, Q, S, T). Annotated using PRODIGY-cryst.

- **apolar_apolar_contacts**: Denotes intermolecular contacts between any of the charged amino acids (A, C, G, F, I, M, L, P, W, V, Y). Annotated using PRODIGY-cryst.

- **interface_atom_gaps_4A**: Number of interface atoms within a 4Å radius of a residue gap. A Gap is determined by residue numbering; regions where one or more of the expected residue index is missing is marked as a gap.

- **missing_interface_residues_4A**: Number of interface residues within a 4Å radius of a residue gap. A Gap is determined by residue numbering; regions where one or more of the expected residue index is missing is marked as a gap.

- **interface_atom_gaps_8A**: Number of interface atoms within an 8Å radius of a residue gap. A Gap is determined by residue numbering; regions where one or more of the expected residue index is missing is marked as a gap.

- **missing_interface_residues_8A**: Number of interface residues within an 8Å radius of a residue gap. A Gap is determined by residue numbering; regions where one or more of the expected residue index is missing is marked as a gap.

- **ECOD_intersection_R**: The number of interface residues that could be mapped to the RCSB-derived ECOD domain protein family name(s) corresponding to the receptor dimer chain. If multiple ECOD domain annotations were found, each domain intersection is delimited with a comma.

- **ECOD_intersection_L**: The number of interface residues that could be mapped to the RCSB-derived ECOD domain protein family name(s) corresponding to the ligand dimer chain. If multiple ECOD domain annotations were found, each domain intersection is delimited with a comma.

- **chain_1_residues**: The residue numbers corresponding to the interface residues of the first (receptor) chain, delimited with a comma.

- **chain_2_residues**: The residue numbers corresponding to the interface residues of the second (ligand) chain, delimited with a comma.

- **n_interchain_bond_atoms**: The number of atoms that form bonds between the two interacting chains. These atoms participate in covalent or non-covalent bonds that contribute to the stability of the interface between the receptor and ligand chains.

- **interchain_res**: The number of residues that form the interface between the two interacting chains. This metric helps quantify the extent of the interaction surface between the receptor and ligand chains.

- **disulfide_bonds**: The number of disulfide bonds formed between cysteine residues of the two interacting chains. Disulfide bonds are covalent bonds that can contribute significantly to the stability of protein interfaces.

- **interchain_bond_or_clash**: The number of bonds or clashes between atoms of the two interacting chains. This includes both stabilizing interactions, such as hydrogen bonds or ionic interactions, and destabilizing interactions, such as steric clashes.

- **potential_transient**: Indicates whether the interaction between the two chains is likely to be transient. Transient interactions are typically less stable and more dynamic compared to permanent interactions, often seen in signaling pathways or regulatory mechanisms.

- **potential_disulfide**: Indicates the potential for the formation of disulfide bonds between the two interacting chains. This is based on the proximity and orientation of cysteine residues, which may form disulfide bonds under the right conditions.

- **neff**: Paired Neff values for the test system members. Calculated using the paired MSA.

- **chain1_neff**: Calculated Neff corresponding to the first (receptor) chain.

- **chain2_neff**: Calculated Neff corresponding to the second (ligand) chain.

##### A.10.2. EXTERNAL AND DERIVED ANNOTATIONS

- **method**: The experimental method for structure determination (XRAY, CRYO-EM, etc.).
- **date**: Date of deposition into RCSB PDB.
- **release_date**: Date of initial release into RCSB PDB.
- **resolution**: The resolution of the experimental structure.
- **assembly**: Which bioassembly is used to derive the structure. 1, 2, 3 means first, second, and third assembly, respectively. All PINDER dimers are derived from the first biological assembly.
- **assembly_details**: How the bioassembly information was derived. Is it author-defined or from another source.
- **oligomeric_details**: Description of the oligomeric state of the protein complex.
- **oligomeric_count**: The oligomeric count associated with the dataset entry.
- **biol_details**: The biological assembly details associated with the dataset entry.
- **asym_id_1**: The first asymmetric identifier (author chain ID)
- **asym_id_2**: The second asymmetric identifier (author chain ID)
- **entity_id_R**: The RCSB PDB 'entity_id' corresponding to the receptor dimer chain.
- **entity_id_L**: The RCSB PDB 'entity_id' corresponding to the ligand dimer chain.
- **pdb_strand_id_R**: The RCSB PDB 'pdb_strand_id' (author chain) corresponding to the receptor dimer chain.
- **pdb_strand_id_L**: The RCSB PDB 'pdb_strand_id' (author chain) corresponding to the ligand dimer chain.
- **ECOD_names_R**: The RCSB-derived ECOD domain protein family name(s) corresponding to the receptor dimer chain. If multiple ECOD domain annotations were found, the domains are delimited with a comma.
- **ECOD_names_L**: The RCSB-derived ECOD domain protein family name(s) corresponding to the ligand dimer chain. If multiple ECOD domain annotations were found, the domains are delimited with a comma.
- **ECOD_ids_R**: The RCSB-derived ECOD domain protein family name(s) corresponding to the receptor dimer chain. If multiple ECOD domain annotations were found, the domains are delimited with a comma.
- **ECOD_ids_L**: The RCSB-derived ECOD domain protein family identifier(s) corresponding to the ligand dimer chain. If multiple ECOD domain annotations were found, the identifiers are delimited with a comma.
- **resi_pdb_R**: The RCSB structure residue numbers corresponding to the receptor chain, delimited with a comma.
- **resi_pdb_L**: The RCSB structure residue numbers corresponding to the ligand chain, delimited with a comma.
- **resi_auth_R**: The RCSB author-defined structure residue numbers corresponding to the receptor chain, delimited with a comma.
- **resi_auth_L**: The RCSB author-defined structure residue numbers corresponding to the ligand chain, delimited with a comma.
- **resi_uniprot_R**: The UniProt-defined residue numbers corresponding to the receptor chain, delimited with a comma.
- **resi_uniprot_L**: The UniProt-defined residue numbers corresponding to the ligand chain, delimited with a comma.
- **sequence**: RCSB-defined entity sequence for each asym identifier.
- **organism**: RCSB-defined source organism for each asym identifier.
- **tax_id**: RCSB-defined source taxonomy identifier for each asym identifier.

- **contains_enzyme**: Whether the dimer contains a protein with an enzyme classification (EC) identifier.
- **EC_R**: RCSB-derived enzyme classification (EC) identifier(s) corresponding to the receptor chain.
- **EC_L**: RCSB-derived enzyme classification (EC) identifier(s) corresponding to the ligand chain.
- **Hchain**: SAbDab Hchain mapped to PINDER.
- **Lchain**: SAbDab Lchain mapped to PINDER.
- **model**: SAbDab model mapped to PINDER.
- **antigen_chain**: SAbDab antigen chain mapped to PINDER.
- **antigen_type**: SAbDab antigen type mapped to PINDER.
- **antigen_het_name**: SAbDab antigen heterogen name mapped to PINDER.
- **antigen_name**: SAbDab antigen name mapped to PINDER.
- **short_header**: SAbDab short header mapped to PINDER.
- **compound**: SAbDab compound mapped to PINDER.
- **heavy_species**: SAbDab heavy species mapped to PINDER.
- **light_species**: SAbDab light species mapped to PINDER.
- **antigen_species**: SAbDab antigen species mapped to PINDER.
- **r_free**: SAbDab ”r free” mapped to PINDER.
- **r_factor**: SAbDab ”r factor” mapped to PINDER.
- **scfv**: SAbDab ”scfv” annotation mapped to PINDER.
- **engineered**: SAbDab ”engineered” annotation mapped to PINDER.
- **heavy_subclass**: SAbDab heavy subclass mapped to PINDER.
- **light_subclass**: SAbDab light subclass mapped to PINDER.
- **light_ctype**: SAbDab light ctype mapped to PINDER.
- **affinity**: SAbDab affinity mapped to PINDER.
- **delta_g**: SAbDab delta g mapped to PINDER.
- **affinity_method**: SAbDab affinity method mapped to PINDER.
- **temperature**: SAbDab temperature mapped to PINDER.
- **pmid**: SAbDab pmid mapped to PINDER.
- **pinder_Hchain_ids**: SAbDab pinder Hchain ids mapped to PINDER.
- **pinder_Hchain_monomers**: SAbDab pinder Hchain monomers mapped to PINDER.
- **pinder_Lchain_ids**: SAbDab pinder Lchain ids mapped to PINDER.
- **pinder_Lchain_monomers**: SAbDab pinder Lchain monomers mapped to PINDER.
- **pinder_antigen_chain_ids**: SAbDab antigen chain ids mapped to PINDER.
- **pinder_antigen_chain_monomers**: SAbDab antigen chains mapped to PINDER monomers.

##### A.10.3. PINDER ASSIGNED ANNOTATIONS

- **chain1_id**: The Receptor chain identifier associated with the dimer entry. Should all be chain 'R'.
- **chain2_id**: The Ligand chain identifier associated with the dimer entry. Should all be chain 'L'.
- **chain_1**: New chain id generated post-bioassembly generation, to reflect the asym_id of the bioassembly and also to ensure that there is no collision of chain ids, for example in homooligomers (receptor chain).
- **chain_2**: New chain id generated post-bioassembly generation, to reflect the asym_id of the bioassembly and also to ensure that there is no collision of chain ids, for example in homooligomers (ligand chain).
- **assembly**: Which bioassembly is used to derive the structure. 1, 2, 3 means first, second, and third assembly, respectively. All PINDER dimers are derived from the first biological assembly.
- **complex_type**: The type of the complex in the dataset entry (homomer or heteromer).
- **resi_R**: The PINDER structure residue numbers corresponding to the receptor chain, delimited with a comma.
- **resi_L**: The PINDER structure residue numbers corresponding to the ligand chain, delimited with a comma.
- **pinder_af2_timesplit**: Whether the PINDER ID is a member of PINDER-XL and is released after the AlphaFold training cutoff date.
- **ialign_af2_similar**: Whether the ID is similar to a member in the AlphaFold training set, as determined by iAlign.
- **ialign_train_similar**: Whether the ID is similar to a member in the PINDER-Train split, as determined by iAlign.
- **ialign_val_similar**: Whether the ID is similar to a member in the PINDER-Val split, as determined by iAlign.

##### A.10.4. DIMER QUALITY CRITERIA FOR SELECTING *proto-test*

Here we detail a full list of quality annotations used to define the *prototest* split prior to de-leaking. Dimer systems are assigned into the *proto-test* group if all of the following conditions are met:

- method = "X-RAY"
- oligomeric_count = 2
- label = "BIO"
- resolution $\leq$ 3.5 Å
- length1 $\geq$ 40
- length2 $\geq$ 40
- interface_atom_gaps_4A = 0
- num_atom_types $\geq$ 3
- number_of_components_1 = 1
- number_of_components_2 = 1
- max_var_1 $\leq$ 0.98
- max_var_2 $\leq$ 0.98

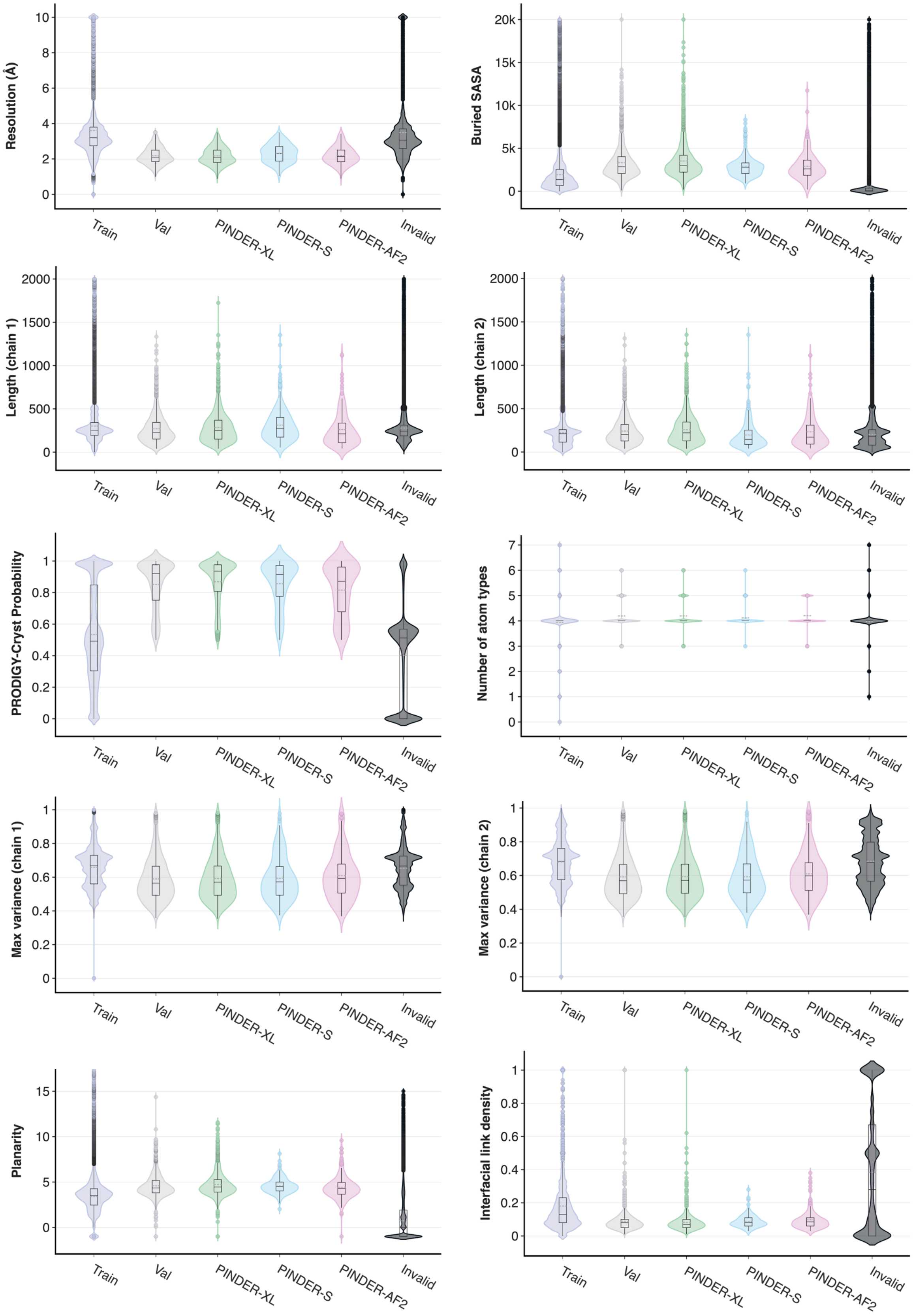


*Figure A.41.* Distribution of metadata annotation values for systems in PINDER Train (purple), Val (gray), XL (green), S (blue), AF2 (pink), and Invalid (black) splits. Selected system annotations include resolution (Å), buried solvent-accessible surface area (SASA), chain lengths, PRODIGY-cryst (Jiménez-García et al., 2019) score, number of atom types, coordinate variance projected along the largest principal component, interface planarity, and link density reported by PRODIGY-cryst.

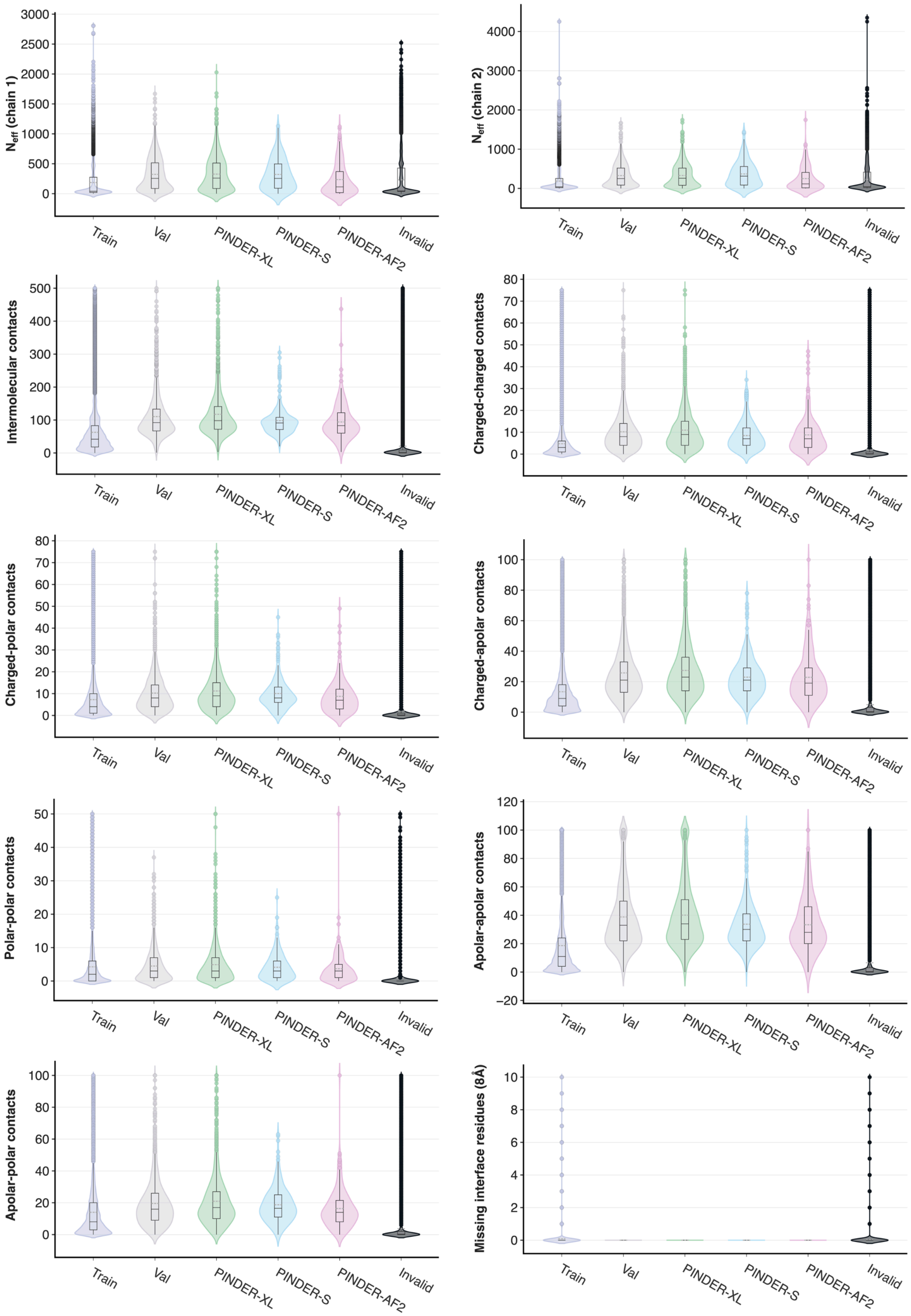


*Figure A.42.* Distribution of additional metadata annotation values for systems in PINDER Train (purple), Val (gray), XL (green), S (blue), AF2 (pink), and Invalid (black) splits. Selected system annotations include number of effective sequences ($N_{\text{eff}}$), number of intermolecular contacts of various types, and number of missing interface residues.

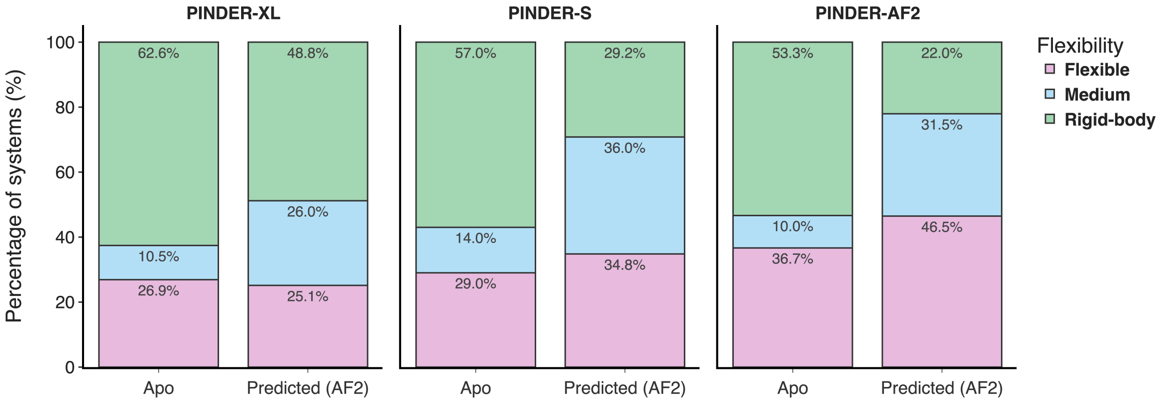


*Figure A.43.* Percent of *apo* and predicted (AFDB) structures in the PINDER-XL, PINDER-S and PINDER-AF2 sets classified by the difficulty of using the *apo* monomer conformations as inputs.

| Program | Parameter | Value | Description |
|---|---|---|---|
| `frodockgrid` | `-m` | 0, 1, 3 | Type of map (0: van der Waals potential map, 1: Electrostatic potential map, 3: Desolvation potential + PDB file with ASA in Occupancy column + ASA projection map.) |
| `frodockgrid` | `-t` | O | Type of complex - A: Antibody/antigen or O: Other |
| `frodockgrid` | `--gs` | 1.0 | Grid spacing |
| `frodockgrid` | `--wr` | 1.4 | Water radius |
| `frodockgrid` | `--sr` | 1.95 | Solvent radius |
| `frodockgrid` | `--fp` | 0 | Fixed points |
| `frodockgrid` | `--an` | 0 | Anisotropy |
| `frodockgrid` | `--rm` | 1.0 | Radius multiplier |
| `frodock` | `--bw` | 32 | Bandwidth |
| `frodock` | `--lmax` | 0.23 | Maximum ligand length |
| `frodock` | `--lmin` | 0.43 | Minimum ligand length |
| `frodock` | `--th` | 10.0 | Threshold |
| `frodock` | `--lw` | 1.0 | Ligand weight |
| `frodock` | `--st` | 2.0 | Step size |
| `frodock` | `-W` | 1.0 | Weight |
| `frodock` | `--np` | 4 | Number of processors |
| `frodock` | `--rd` | 12.0 | Radius |
| `frodock` | `--td` | 0 | Translation distance |
| `frodockcluster` | `-d` | 5.0 | Distance threshold |
| `frodockcluster` | `--nc` | 40 | Number of clusters |
| `frodockcluster` | `--ns` | all solutions | Number of solutions |
| `frodockcluster` | `-c` | 1.0 | Clustering coefficient |
| `frodockview` | `-r` | 1-40 | Range of solutions |

*Table A.2.* FroDock parameters

| Parameter Name | Sub-Parameter | Parameter Value | Description |
|---|---|---|---|
| receptorSeg | low_patch_thr, high_patch_thr | 10.0, 20.0 | Min and max patch parameters |
| receptorSeg | prune_thr | 1.5 | Minimal distance between points inside the patch |
| receptorSeg | knob, flat, hole | 1, 0, 1 | Types of patches to dock (1-use, 0-do not use) |
| receptorSeg | hot_spot_filter_type | 0 | Hot spot filter type: None - 0, Antibody - 1, Antigen - 2, Protease - 3, Inhibitor - 4, Drug - 5 |
| ligandSeg | low_patch_thr, high_patch_thr | 10.0, 20.0 | Min and max patch parameters |
| ligandSeg | prune_thr | 1.5 | Minimal distance between points inside the patch |
| ligandSeg | knob, flat, hole | 1, 0, 1 | Types of patches to dock (1-use, 0-do not use) |
| ligandSeg | hot_spot_filter_type | 0 | Hot spot filter type: None - 0, Antibody - 1, Antigen - 2, Protease - 3, Inhibitor - 4, Drug - 5 |
| scoreParams | small_interfaces_ratio | 0.3 | Ratio of the low scoring transforms to be removed |
| scoreParams | max_penetration | -5.0 | Maximal allowed penetration between molecules surfaces |
| scoreParams | ns_thr | 0.5 | Normal score threshold |
| scoreParams | rec_as_thr, lig_as_thr | 0.0, 0.0 | Minimal ratio of the active site area in the solutions |
| scoreParams | patch_res_num | 1500 | Number of results to consider in each patch |
| scoreParams | w1, w2, w3, w4, w5 | -8, -4, 0, 1, 0 | Scoring weights for ranges: [-5.0,-3.6],[-3.6,-2.2],[-2.2,-1.0],[-1.0,1.0],[1.0-up] |
| desolvationParams | energy_thr | 500.0 | Remove all results with desolvation energy higher than threshold |
| desolvationParams | cut_off_ratio | 1.0 | Ratio of low energy results to be kept |
| clusterParams | rotationVoxelSize | 0.1 | Rotation voxel size |
| clusterParams | discardClustersSmaller | 4 | Discard clusters smaller than this value |
| clusterParams | rmsd | 2.0 | RMSD value |
| clusterParams | final_clustering_rmsd | 4.0 | Final clustering RMSD value |
| baseParams | min_base_dist | 4.0 | Minimum base distance |
| baseParams | max_base_dist | 13.0 | Maximum base distance |
| baseParams | num_patches | 2 | Number of patches for base |
| matchingParams | geo_dist_thr | 1.5 | Geometric distance threshold |
| matchingParams | dist_thr | 1.5 | Distance threshold |
| matchingParams | angle_thr | 0.4 | Angle threshold |
| matchingParams | torsion_thr | 0.5 | Torsion threshold |
| matchingParams | angle_sum_thr | 0.9 | Angle sum threshold |
| receptorGrid | gridStep | 0.5 | Grid step |
| receptorGrid | maxDistInDistFunction | 6.0 | Maximum distance in distance function |
| receptorGrid | vol_func_radius | 6.0 | Volume function radius |
| ligandGrid | gridStep | 0.5 | Grid step |
| ligandGrid | maxDistInDistFunction | 6.0 | Maximum distance in distance function |
| ligandGrid | vol_func_radius | 6.0 | Volume function radius |
| receptorMs | density | 10.0 | Density |
| receptorMs | probe_radius | 1.8 | Probe radius |
| ligandMs | density | 10.0 | Density |
| ligandMs | probe_radius | 1.8 | Probe radius |

*Table A.3.* PatchDock Parameters
